# Detailed characterization of the UMAMITs provides insight into their evolution, functional properties as amino acid transporters and role in the plant

**DOI:** 10.1101/2021.02.18.431885

**Authors:** Chengsong Zhao, Réjane Pratelli, Shi Yu, Brett Shelley, Eva Collakova, Guillaume Pilot

**Affiliations:** School of Plant and Environmental Sciences, Virginia Tech, Blacksburg, VA 24060, USA

## Abstract

Amino acid transporters play a critical role in distributing amino acids within the cell compartments and between the plant organs. Despite this importance, relatively few amino acid transporter genes have been characterized and their role elucidated with certainty. Two main families of proteins encode amino acid transporters in plants: the Amino Acid-Polyamine-Organocation superfamily, containing mostly importers, and the Usually Multiple Acids Move In and out Transporter family, apparently encoding exporters, totaling about 100 genes in Arabidopsis alone. Knowledge on UMAMITs is scarce, focused on six Arabidopsis genes and a handful of genes from other species. To get insight into the role of the members of this family and provide data to be used for future characterization, we studied the evolution of the UMAMITs in plants, and determined the functional properties, the structure, and the localization of the 44 Arabidopsis UMAMITs. Our analysis showed that the AtUMAMIT are essentially localized at the tonoplast or the plasma membrane, and that most of them are able to export amino acids from the cytosol, confirming a role in intra- and inter-cellular amino acid transport. As an example, this set of data was used to hypothesize the role of a few AtUMAMITs in the plant and the cell.

## Introduction

Along with sugars and carboxylic acids, amino acids are major players in cell metabolism. They are essential nitrogen carriers used in a large number of metabolic reactions and in translocating nitrogen between the various organs of most multicellular organisms. In plants, amino acids are also used as precursors of specialized metabolites and as a dynamic, readily available storage form of nitrogen. Amino acid transporters, *i.e.* membrane proteins specialized in the transfer of amino acids from one side of the membrane of cells and organelles to the other, play critical roles in controlling the flow of nitrogen between the various compartments, cells, and organs of the plant. Similar to metabolic activity, amino acid transporter activity is controlled and adjusted to the growth conditions and the response to biotic and abiotic stresses.

The first amino acid transporters discovered in plants belong to the Amino Acid-Polyamine-Organocation (APC) and the Amino Acid/Auxin Permease (AAAP) families (2.A.3 and 2.A.18 respectively in the Transporter Classification Database - TCDB; http://www.tcdb.org/), both families being members of the APC superfamily (Vastermark *et al.*, 2014). Arabidopsis and rice genomes contain a total of 63 and 85 APC superfamily member genes respectively (Zhao *et al.*, 2012; Pratelli and Pilot, 2014). With a few exceptions, all the proteins from the APC superfamily that are functionally characterized are importers, *i.e.* catalyze the transport of amino acids across membranes towards the cytosol (either from the outside of the cell, or from intracellular compartments, like the vacuole). The existence of exporters, catalyzing the transport out of the cytosol, was experimentally proven in the 80’s; however, the first amino acid transporter with export properties was not isolated and characterized until 20 years after amino acid importers were (Ladwig *et al.*, 2012). This protein, SiAR1/AtUMAMIT18, belongs to the Usually Multiple Acids Move In and out Transporter family (UMAMIT, or P-DME from TCDB; 2.A.7.4), whose founding member is the gene MtN21 (MtrunA17_Chr3g0081511), a “nodulin” whose expression is induced during nodulation of *Medicago truncatula* roots (Gamas *et al.*, 1996; Denancé *et al.*, 2014b).

Arabidopsis UMAMITs are the most characterized. AtUMAMIT11, 14, 24, 25, 28, 29 and SiAR1/AtUMAMIT18 fulfill roles in seed filling, phloem loading and unloading (Ladwig *et al.*, 2012; Müller *et al.*, 2015; Besnard *et al.*, 2016; Besnard *et al.*, 2018). A Ph.D. thesis reported the localization of several other AtUMAMITs in the seed tissues (AtUMAMIT01, 02, 10, 15, 21, 23, 25, 34 and 41) (Müller, 2016), but no genetic study has been performed to test for their function in this organ. Interestingly, this Ph.D. work reported that some of those AtUMAMITs are induced by infestation by nematodes (Müller, 2016). This parallels the fact that other AtUMAMITs appear to play a role during infection by pathogens, *i.e. Ralstonia solanacearum*, *Xanthomonas campestris* or *Verticillium dahliae* (WAT1/AtUMAMIT05) (Denancé *et al.*, 2013), and *Phytophthora parasitica*, *Pseudomonas syringae* or *Golovinomyces cichoracearum* (AtUMAMIT36/RTP1) (Pan *et al.*, 2016), possibly by modulating the activity of the salicylic acid pathway. AtUMAMIT05 has also been shown to affect auxin content and lignin deposition (Ranocha *et al.*, 2010; Denancé *et al.*, 2013; Ranocha *et al.*, 2013; Denancé *et al.*, 2014a). This relationship between UMAMITs and auxin seems conserved across species, because it has also been identified in pine and cotton, where the AtUMAMIT05 homologs 5NG4 and GsWAT1 were found induced by auxin (Busov *et al.*, 2004; Tang *et al.*, 2019). In addition, GsWAT1 was found involved in pathogen response and lignin deposition. A study of the *Pinus sylvestris* 5NG4 and MtN21-like-a/b proteins (respectively homologs of AtUMAMT05 and 09) revealed that the corresponding genes are responding to auxin and the presence of the mycorrhizal fungus *Laccaria bicolor* (Heller *et al.*, 2008; Heller *et al.*, 2012), making them candidates for genes involved in symbiosis. The yet under-explored UMAMIT family thus appears to be the “missing link in nitrogen cycling” (Okumoto and Pilot, 2011), but seems also involved in other pathways related to aromatic amino acids, reminiscent of the AUX1 and LAX proteins from the AAAP amino acid transporter family that function as auxin importers (Yang *et al.*, 2006).

Contrary to the Amino Acid Permease (AAP) subfamily of amino acid transporters, whose functional properties have been extensively studied (Rentsch *et al.*, 2007; Tegeder and Rentsch, 2010), little is known about the substrate diversity or the transport mechanism of the UMAMITs. AtUMAMIT14 and 18 have been shown to transport amino acids in a bidirectional fashion, probably downstream from their electrochemical gradient, making them facilitators (Ladwig *et al.*, 2012; Müller *et al.*, 2015). Yet, those studies have not explored the specificity for each amino acid and if other compounds are potential substrates, a justified question since AtUMAMIT05 can transport auxin (Ranocha *et al.*, 2013). Despite the appeal and expected importance of this gene family in relation to nitrogen nutrition, little has been done to understand its structure, origin, and answer the question whether all the members are amino acid facilitators, similar to AtUMAMIT14 and 18. A few publications have analyzed the phylogenic relationships of the UMAMITs from some species (Busov *et al.*, 2004; Ladwig *et al.*, 2012; Denancé *et al.*, 2014b; Müller *et al.*, 2015), but the studies have remained limited in terms of number of species used and conclusions reached.

The goal of the present work was to provide an extensive overview of the family, using phylogenic, structural and expression data, complemented with study of sub-cellular localization and preliminary exploration of the functional properties of all the AtUMAMITs. We specifically wanted to answer the questions whether (1) all the AtUMAMITs are amino acid exporters/facilitators; (2) they are addressed to the plasma membrane, the tonoplast or other membranes; (3) there is any correlation between position in the phylogenic tree and the protein properties or expression; and (4) any information on these genes could be gathered that would help the plant research community in better understanding nitrogen flow in the plant.

## Results

### Organization of the UMAMIT family across species

Mining the Arabidopsis genome identified 47 loci encoding UMAMIT proteins. Three of them encoded proteins missing transmembrane domains and were hence qualified as pseudogenes: AtUMAMIT16-Ψ, 39-Ψ and 43-Ψ; Supplemental Figure 1; Supplemental File 1A). Interestingly, these pseudogenes produced transcripts detected by RNAseq analyses, but ranked among the five most lowly-expressed genes of the family (Supplemental Figure 2). Similarly, mining the rice genome identified 53 UMAMIT loci, including two pseudogenes (OsUMAMIT32-Ψ and 53-Ψ; Supplemental Figure 1; Supplemental File 1B). The Arabidopsis and rice loci were numbered according to the position of the genes in the phylogenic tree (see below), with pseudogenes indicated by the Greek letter “Ψ” (we released the names of the AtUMAMITs in the TAIR database in 2008). These pseudogenes were not used in subsequent analyses.

Phylogenic relationships between the UMAMITs were first determined using four representative species whose genomes were available (*Physcomitrella patens*, *Selaginella moellendorffii*, *Arabidopsis thaliana*, *Oryza sativa*), and the two conifers *Pinus pinaster* and *Picea abies* (pine and fir) (Supplemental Figure 1). Maximum likelihood analysis grouped the genes into 10 clades (named A to J): clades C and D contained the most members, clade A contained all *Physcomitrella* and *Selaginella* sequences, while clade H was exclusively composed of pine and fir sequences (Figure 1). The AtUMAMIT proteins shared relatively low sequence identity (~32%) with one another, except for proteins from the same clade (50-90% identity) (Supplemental File 1D). The position of the AtUMAMIT loci in the Arabidopsis genome identified several clusters of genes (particularly evident for genes from clade C, D and J) which mirrored the sequence identities, suggesting that these genes arose from recent duplication events (Supplemental Figure 3). The phylogenic study was expanded to include UMAMITs from 38 plant species (Supplemental File 1E) totaling 1466 protein sequences (Supplemental File 1F). Some patterns arose from the maximum likelihood analysis: (1) Conifer sequences were absent from clades E, F, G, I and J, but made a clade on their own, clade H. (2) Some *Physcomitrella* and *Selaginella* sequences were not contained in clade A but grouped together, and represent highly divergent sequences. (3) Clade J contained only dicot sequences. (4) No *Brassicaceae* sequence was identified in clade G. (5) Clades C and D contained 46% of the sequences of the tree. (6) Monocot sequences were enriched in clades A, D, E and I (1.4, 1.4, 2.2 and 1.3 times respectively) compared to their proportion in the 1466 sequences (Supplemental Figure 4; Supplemental File 1G).

**Figure 1:**
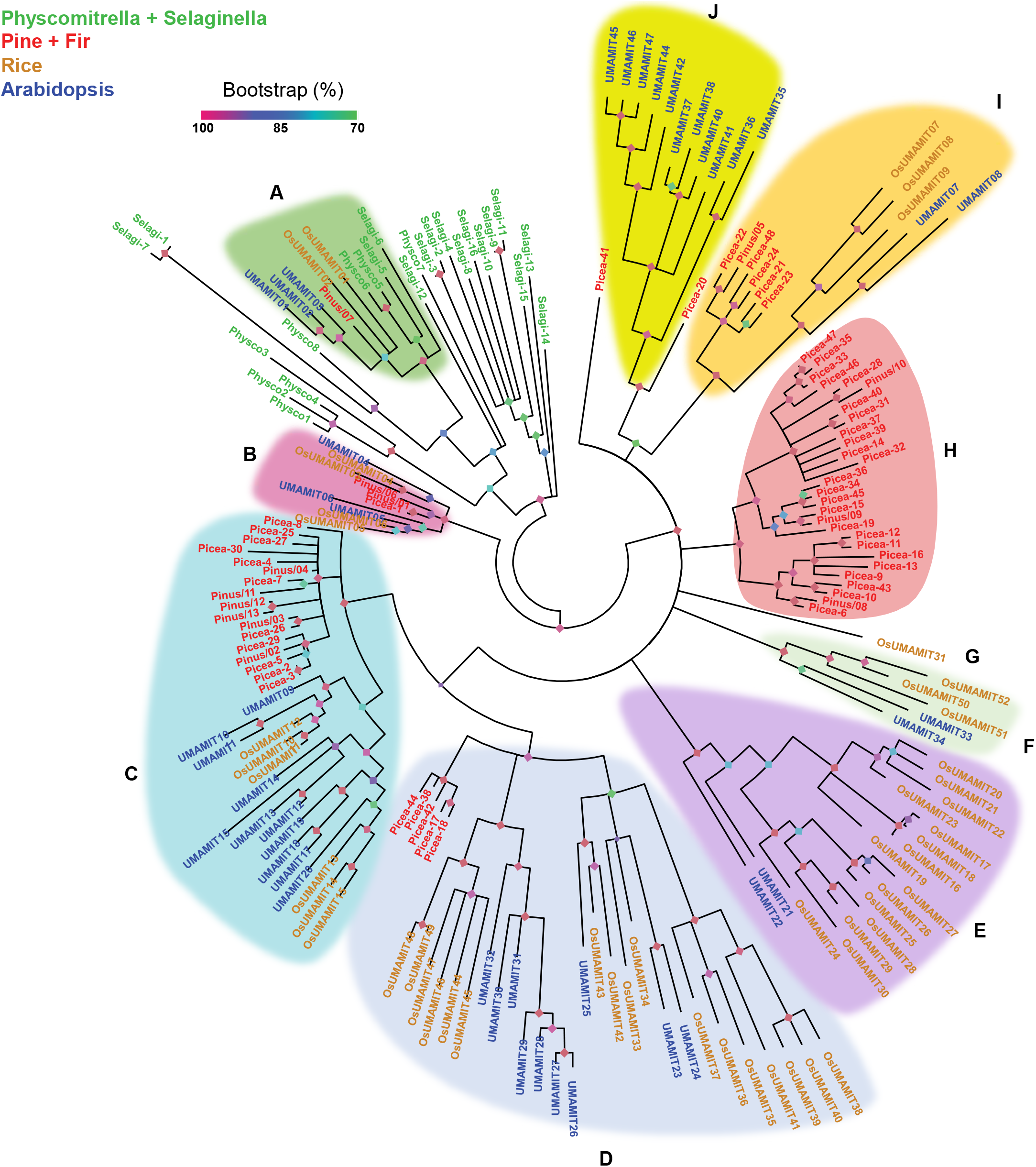
Phylogenetic tree of UMAMIT proteins from six selected species. Ten clades were identified (A to J), with clade G represented here by only one member (see Supplemental Figure 4 for more species constituting this clade). Most *Physcomitrella* and *Selaginella* sequences could not be grouped into one clade in this tree. The tree was constructed using RaxML 8.1.3, with gamma model for rate of heterogeneity, the JTT substitution matrix and 10,000 bootstraps. The sequences from *Physcomitrella patens*, *Selaginella moellendorffii*, *Pinus pinaster*, *Picea abies*, *Arabidopsis thaliana* and *Oryza sativa* used for the tree construction are given in Supplemental File 1C.

Analysis of the gene structure from the 180 sequences used for the first phylogenic analysis identified six intron insertion sites, with 69% of the genes having six introns (Figure 2A, Supplemental File 2A). A few genes from *Selaginella*, rice and fir seemed to have one additional intron (this could not be ascertained for pine and *Selaginella* because of the medium quality of the genome sequences). Gene structure was not particularly conserved within clades, with the exception of clades D-2, D-3 and D-5, whose Arabidopsis and rice genes have all the introns at position 4 missing (Supplemental File 2B). These observations suggest that an ancestral *UMAMIT* gene had 6 introns, and some introns were lost after speciation during the evolution.

**Figure 2:**
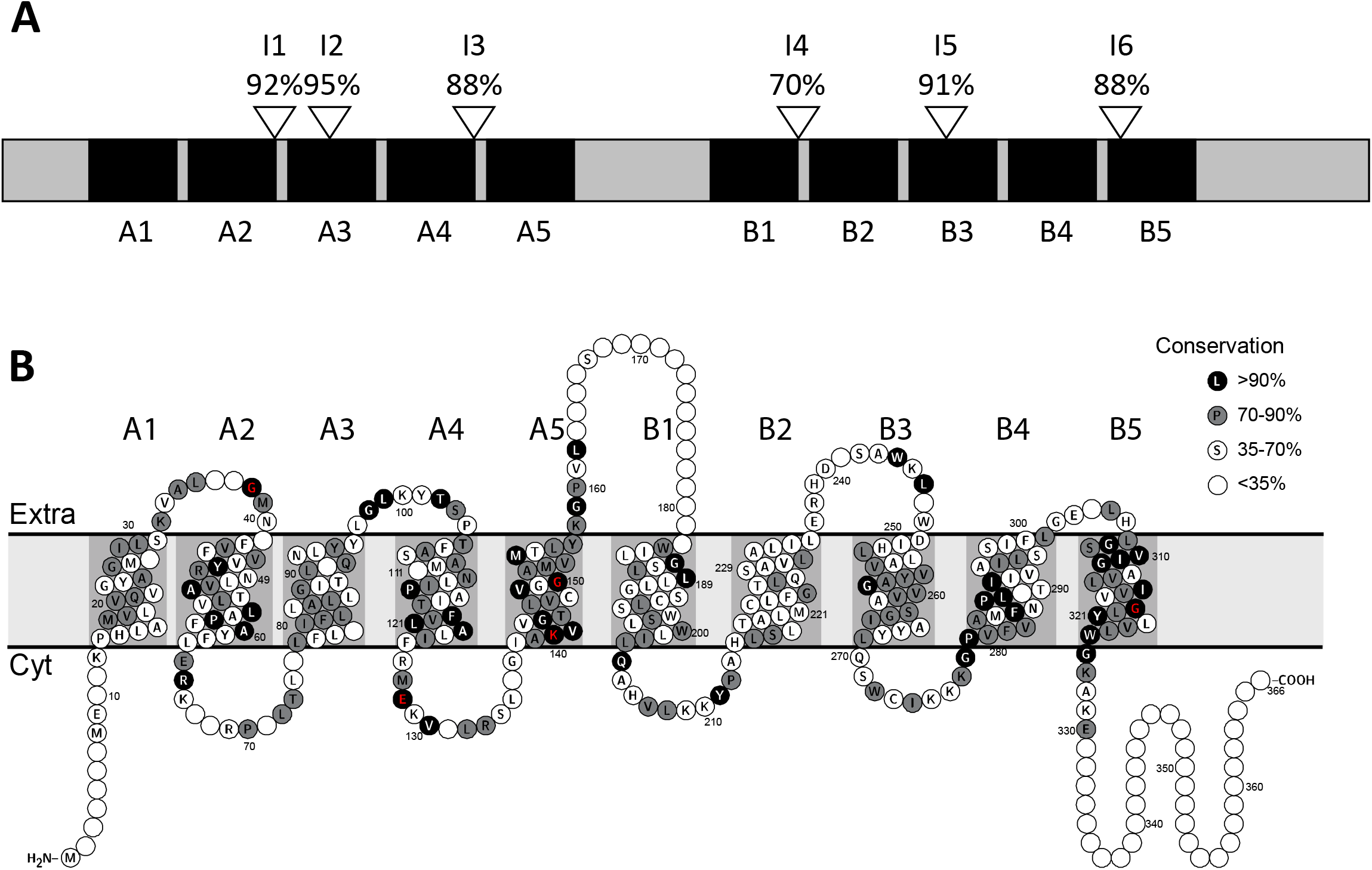
Gene and protein structure of the UMAMIT proteins from Arabidopsis, rice, pine and Physcomitrella. A. Schematic of the protein structure, with the position of the six main intron positions (I1 to I6). The numbers correspond to the percentage of genes (out of 180) with an intron at the corresponding positions. More details are available in Supplemental File 2A. B. Percentages of conservation at each position of a consensus protein sequence of the UMAMITs. Residues mutagenized in UMAMIT14 are indicated in red, localized on the diagram at positions 38 (G33), 128 (E122), 140 (K134), 150 (G144) and 319 (G315). Consensus sequence and percentages were computed by AlignX (Vector NTI) from the multiple sequence alignment of Supplemental Figure 1, and the two dimensional model was constructed by Protter http://wlab.ethz.ch/protter/start/ (Omasits *et al.*, 2014). A1 to A5, transmembrane domains of the first structural repeat, B1 to B5, transmembrane domains of the second structural repeat. Extra, extra-cytosolic side; Cyt, cytosolic side.

**Figure 3:**
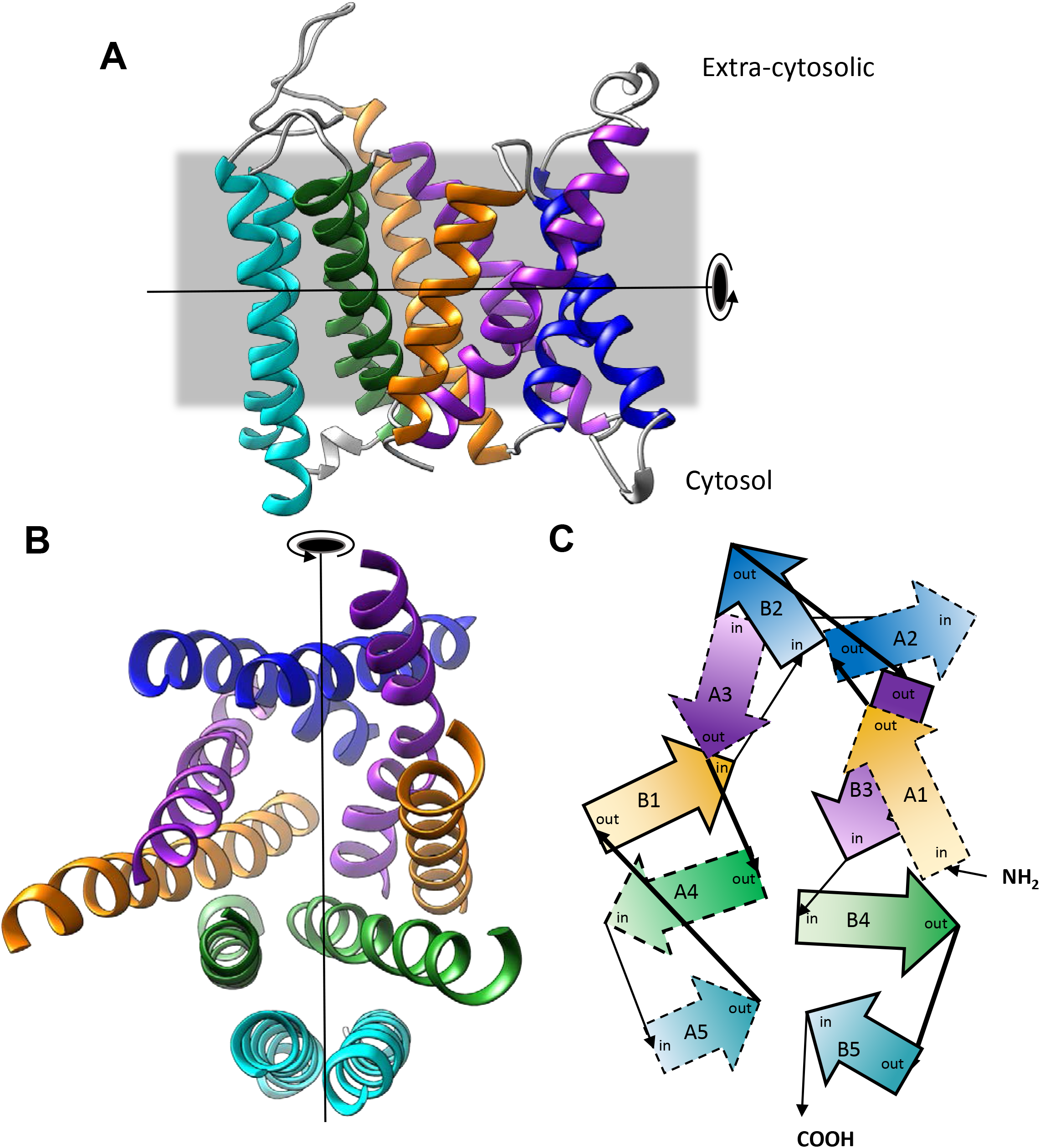
Homology model of the AtUMAMIT14 protein. A. Side view of the homology model created using SwissModel, extra-cytosolic side on the top. Membrane position (gray rectangle) was determined using the PPM tool (Lomize *et al.*, 2012). Colors correspond to the homologous helices in the protein (*e.g.* A1 and B1 in orange) to show the arrangement of the transmembrane domains of the two structural repeats. Loops are colored in gray. B. Same model, seen from the extra cytosolic side (top of model in A), with loops hidden. C. Diagram of model in B, with the direction of the transmembrane domains indicated by arrows. Extra-cytosolic loops are indicated by thick strokes, cytosolic loops indicated by thin strokes. The symmetry axis is represented by a line with the rotating arrow.

### UMAMIT protein structure

The three membrane protein topology prediction tools TMHMM (Krogh *et al.*, 2001), PolyPhobius (Kall *et al.*, 2007) and TOPCONS (Tsirigos *et al.*, 2015) predicted the UMAMITs to have 10 helical transmembrane (TM) domains. Early work on the Plant Drug and Metabolite Exporter family suggested that the structure arose from the duplication of a 5 TM ancestor, with the two halves showing a similar structure and sequence (Jack *et al.*, 2001). In good agreement with this hypothesis, the first 5 TMs showed sequence similarity to the last 5 TMs of the rice and Arabidopsis UMAMITs (Supplemental Figure 5). TM1-5 and TM6-10 were named TM A1-5 and B1-5, respectively, to emphasize the homology of the helices from each structural domain. Placing conserved residues on a topological drawing of the UMAMIT proteins showed that the most highly conserved TMs are A5 and B5, with inter-TM loops generally displaying low sequence conservation (Figure 2B).

The structure of AtUMAMIT14, used as a model for the UMAMIT proteins, was predicted by homology modeling using the Phyre2 (Kelley *et al.*, 2015), Robetta (Song *et al.*, 2013), IntFOLD4 (McGuffin *et al.*, 2019) and SwissModel (Waterhouse *et al.*, 2018) servers. The similarity of the output structures determined by TM-SCORE (Xu and Zhang, 2010) showed that the models were close to one another (scores between 0.65 and 0.77), with the models from Robetta and SwissModel having the highest quality scores determined by QMEAN (Studer *et al.*, 2014). Consequently, the model from SwissModel was used for further study. It became apparent that the homologous helices (*e.g.* A1 and B1, A2 and B2, etc.) were positioned next to each other in inverted orientations so that the protein displays a quasi-symmetry around an axis lying within the plane of the membrane (Figure 4). Based on the crystal structure of a triose-phosphate/phosphate translocator and a nucleotide sugar transporter used as a template for modeling, UMAMIT may form dimers, with TMs A5 and B5 at the interface between the two subunits (Supplemental Figure 6). Interestingly, the sequence of these two TMs is the most conserved amongst the UMAMITs (see above, Figure 2B)

**Figure 4:**
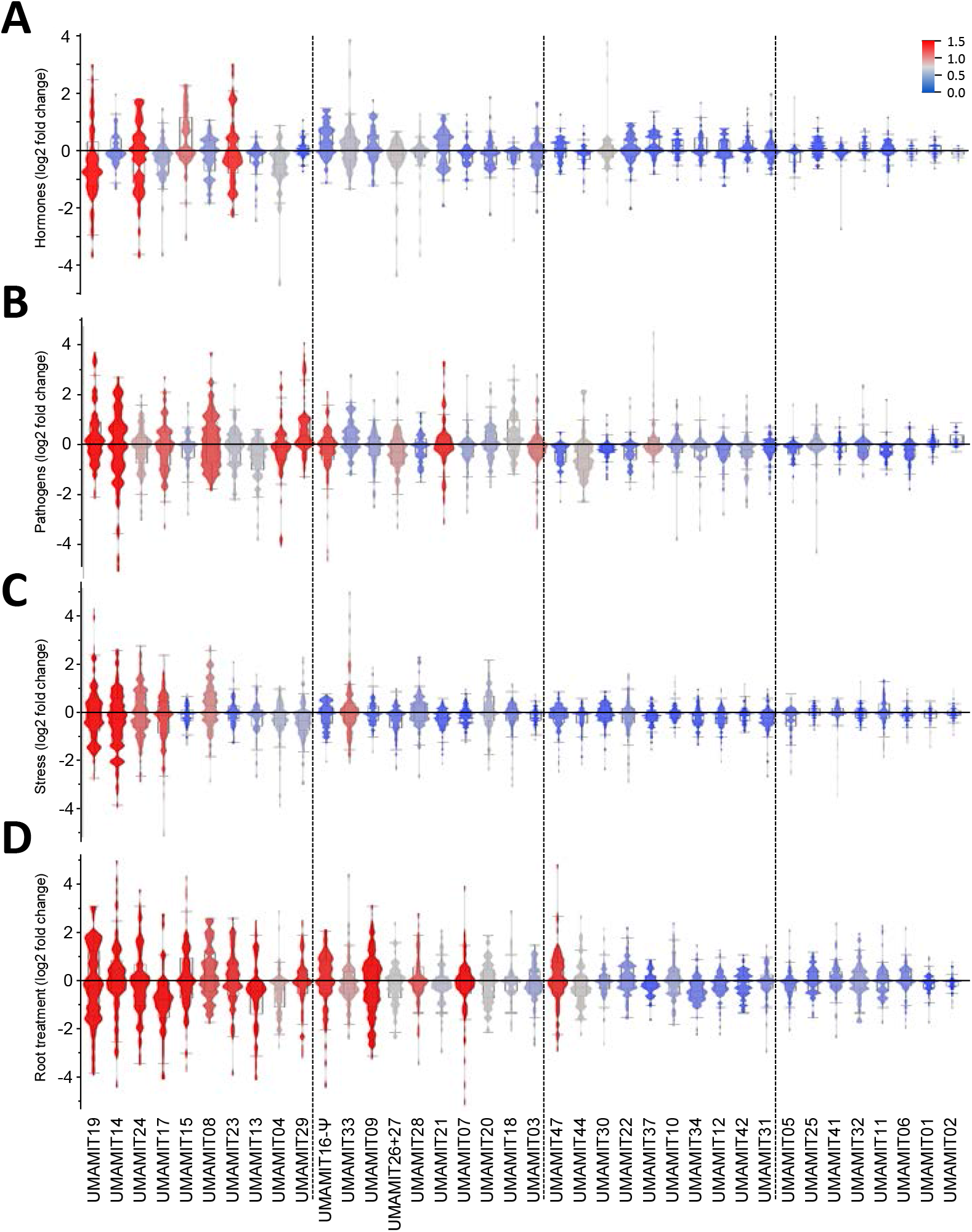
Summary of the changes in the AtUMAMIT mRNA content in response to various conditions. Violin plot of the change in mRNA content of each of the AtUMAMIT genes. Each data point corresponds to one condition from the AtGenexpress data set, analyzed using the Affymetrix ATH1 chip, which is missing six AtUMAMITs, and has a probe that recognizes both AtUMAMIT26 and AtUMAMIT27. Data used for this graph are available in Supplemental File 3. Background boxes correspond to the minimum, 25^th^ percentile, median, 75^th^ percentile and maximum (as defined by JMP software). Areas are colored according to the variance of the dataset for each gene (from 0 to 1.5). A. Whole seedlings treated by various phytohormones, hormone inhibitors and drugs. B. Leaf or seedlings infected or treated by *Erysiphe orontii*, *Hyaloperonospora arabidopsidis*, *Phytophthora infestans*, *Pseudomonas syringae* and elicitors. C. Roots and shoots wounded or grown under cold, osmotic, salt, drought, oxidative, genotoxic, wounding, heat and UV-B conditions. D. Root tissues subjected to Pi and Fe starvation, addition of nitrogen or salt, or pH change.

### Localization of expression of the *AtUMAMITs*

To get a glimpse on the diversity of the expression patterns of the UMAMITs, we focused on the Arabidopsis family, for which a profusion of public data is available, notably the AtGenExpress dataset (Schmid *et al.*, 2005) (Table 1). Unfortunately, the ATH1 Affymetrix chip used for these analyses lacked probes for six members (from clade J), namely *AtUMAMIT35, 36, 38, 40, 45* and *46*. The mRNA accumulation patterns were very diverse, but could be divided into seven groups, composed of genes predominantly expressed in hypocotyl/stem, root, stamen, silique, embryo, pollen or throughout the plant (Supplemental File 3A). In roots, availability of spatiotemporal data (Brady *et al.*, 2007; Cartwright *et al.*, 2009) enabled defining the expression of the genes down to the tissue types. As above, genes could be grouped according to their expression patterns, into procambium-, phloem-, phloem/xylem-, xylem- and peripheral layers-expressed genes (Table 1; Supplemental File 3B). Finally, about 10 *AtUMAMITs* were found expressed at high levels in seeds, mainly in the seed coat (Supplemental File 3C). PCA analysis of these data show that the expression profiles of several AtUMAMITs in the plant, root or seed are distinct from the bulk of the other genes, suggesting that they fulfil unique functions – e.g. *AtUMAMIT05* and *41* ubiquitously expressed in the plant, *AtUMAMIT07* specific of the pollen, and many Clade C- and D-*AtUMAMITs* in the seed (Supplemental Figure 7).

**Table 1:**
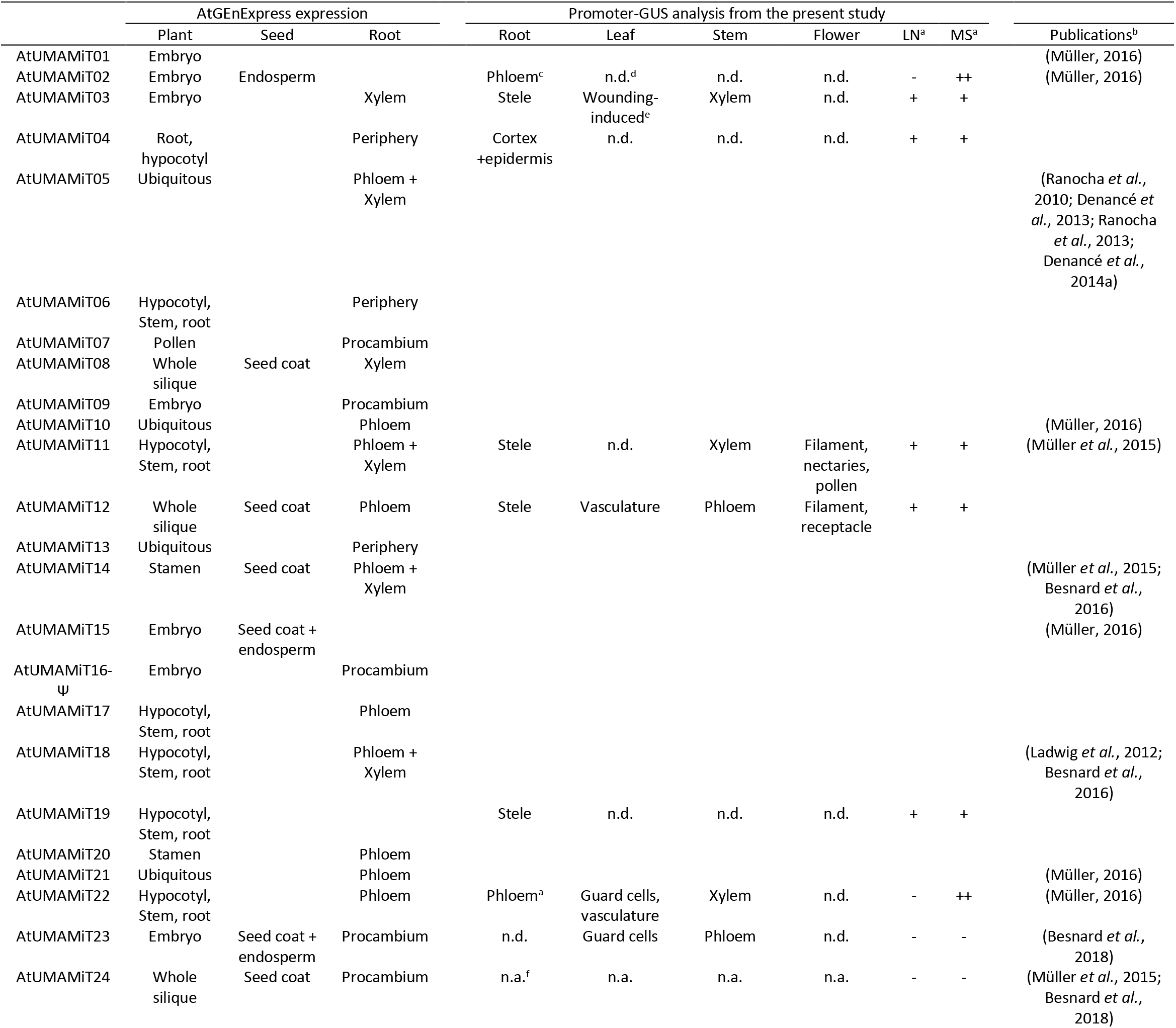
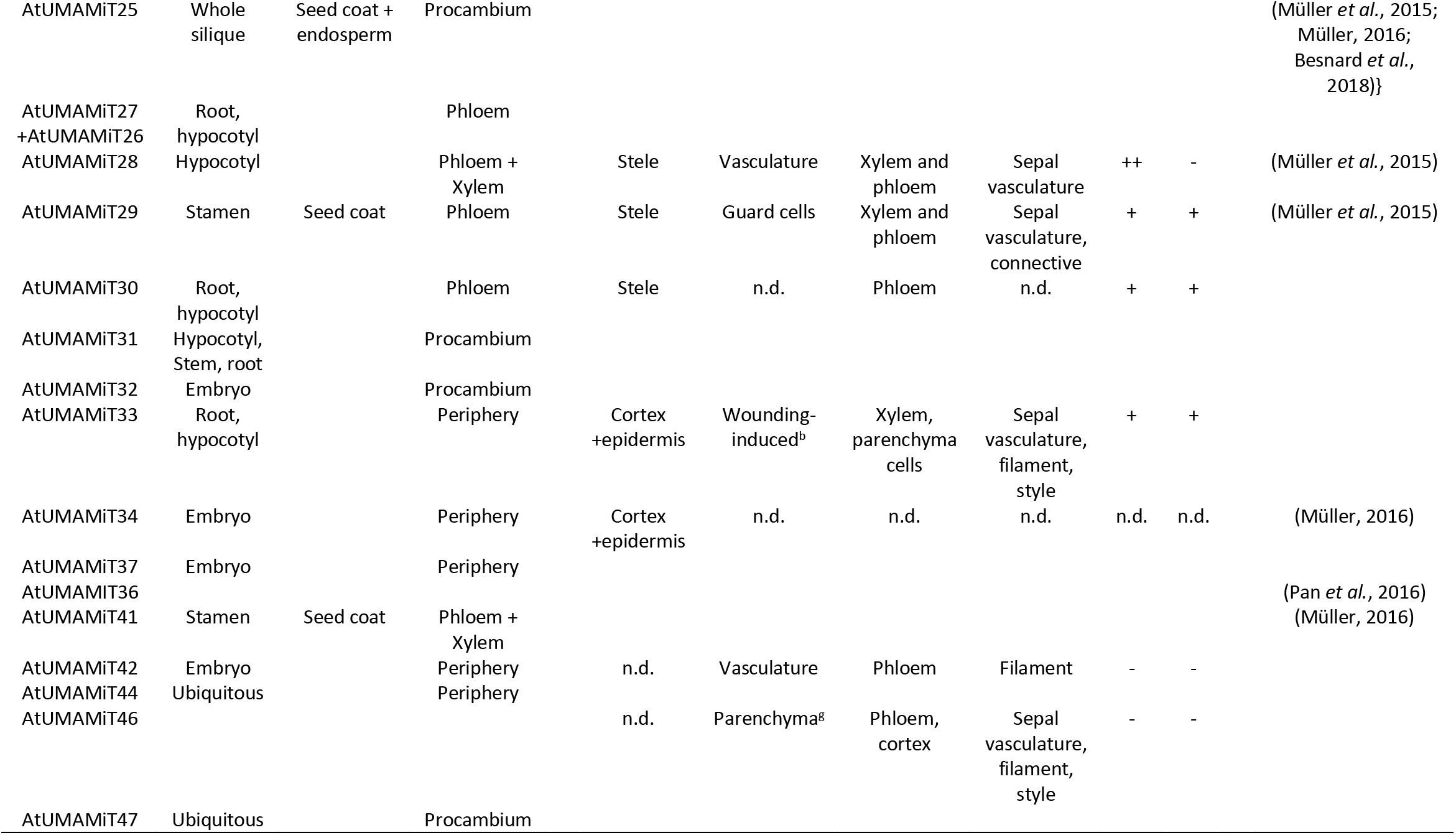
Localization of the promoter activity for the selected AtUMAMITs using the translational GUS reporter assay. ^a^ plants were grown on low (LN) or high (MS) nitrogen media. “-“, no staining; “+”, light staining; “++”, strong staining. ^b^ publication or thesis reporting a study of the AtUMAMIT in question. ^c^ phloem localization was inferred by the pattern of the staining, no cross-section was performed (only the phloem, but not the xylem reaches the dividing zone in the root). ^d^ “n.d.” not detected. ^e^ since the leaves were cut in large pieces before fixing and staining, development of a strong staining at the margin of the leaves is an indication of a wounding-responsive gene. ^f^ “n.a.”, not assayed ^g^ parenchyma localization was inferred from the diffuse staining on the leaf, no cross section was performed.

To confirm the precise localization of the expression obtained from publicly available microarray data, 16 genes were chosen, based on their mRNA levels in various organs: *AtUMAMIT02, 03, 04, 11, 12, 19, 22, 23, 24, 28, 29, 30, 33, 34, 42* and *46* (note that expression of *AtUMAMIT11*, *24*, *28* and *29* has been independently reported (Müller *et al.*, 2015)). The expression patterns of these 16 genes were investigated using the transcriptional β-glucuronidase (GUS) reporter assay (Supplemental Figures 8), but due to unexpected difficulties of getting reliable staining in the siliques and seeds (data not shown), we focused on the roots, vegetative parts and flowers. The histochemical staining mostly agreed with the AtGeneExpress and translatome datasets for expression in the root, leaves, flower and/or vasculature (Table 1; Supplemental Figure 9), confirming the microarray data using a different, complementary assay. This approach also provides information on the localization of several AtUMAMIT in shoot phloem (AtUMAMIT12, 23, 28, 29, 42, 46), xylem parenchyma (AtUMAMT03, 11, 22, 28, 29, 33) and guard cells (AtUMAMIT22, 23, 29).

### Response to treatments of the *AtUMAMITs*

The change in mRNA content of the AtUMAMITs in response to treatments was studied by mining the AtGenExpress dataset, including the response of plant organs and specific root tissues to hormone, pathogen and abiotic treatments (arabidopsis.org/portals/expression/microarray/ATGenExpress.jsp and references therein). Expectedly, mRNA levels of most genes changed in response to several treatments (Supplemental File 3D, E, F and G). Graphing the distribution of the data points and the variance (Figure 4) identified *AtUMAMIT19, 14, 24* and *17* among the genes most subject to regulation by the environment, while AtUMAMIT01 and 02 were the least sensitive. For instance, *AtUMAMIT19* mRNA levels respond strongly to various stresses, being induced by many pathogens, ABA, ABA-involving stresses and phosphate starvation, while repressed in roots by nitrogen (Supplemental File 3). PCA analysis further helped differentiate the highly responding genes from one another, and showed that the expression of many *AtUMAMITs* from Clades C and D is affected by various treatments (Supplemental Figure 10). A slight downward trend was observed between variance in expression change and expression level, showing that the most highly expressed genes (e.g. AtUMAMIT05 and 41) are those least susceptible to change in expression at the mRNA level (Supplemental Figure 11). It is possible that the corresponding proteins are involved in housekeeping processes common to all cells.

During the histochemical staining of the AtUMAMITp-GUS lines above, we noticed that *AtUMAMIT03* and *33* promoters were induced at the place where leaves were cut before staining (Supplemental Figure 8), suggesting that these promoters are induced by wounding. These observations are in good agreement with AtGenExpress data for AtUMAMIT33, but not for AtUMAMIT03 (Supplemental File 3E). Finally, to test the effects of nitrogen supply on the *AtUMAMIT* promoter activity, plants were grown on ½ MS medium or on minimal medium for 10 days, corresponding to high and low nitrogen conditions respectively, and staining intensities were compared. The promoter activity of most tested *AtUMAMITs* was not influenced by the nature of the growth medium, except for *AtUMAMIT02* and *22* which were induced in the root stele by high nitrogen conditions, and for *AtUMAMIT28* which was suppressed (Supplemental Figure 12; Table 1).

### Putative subcellular localizations of the AtUMAMITs

To estimate the diversity of subcellular localization of the AtUMAMIT proteins in the cell, the 44 cDNAs was fused to the GFP coding sequence to lead to AtUMAMIT-GFP fusion proteins, and expressed in *Nicotiana benthamiana* epidermis cells. Fifteen AtUMAMITs were targeted to the plasma membrane, 19 to the tonoplast, but 31 were found in more than one location, mostly the ER (9 proteins) or Golgi/endosomes (29 proteins) (Table 2; Supplemental Figure 13). No signal could be detected for AtUMAMIT07, 24, 27 and 31. Our localization results agree with previous literature concerning AtUMAMIT05, 14, 18, 25, 29 and 36, which have been previously localized as GFP fusions expressed in *N. benthamiana* or Arabidopsis (Ranocha *et al.*, 2010; Ladwig *et al.*, 2012; Müller *et al.*, 2015; Besnard *et al.*, 2016; Pan *et al.*, 2016; Besnard *et al.*, 2018). The localizations in the ER or Golgi/endosomal structures might correspond to the proteins being synthesized, sorted or *en route* to other locations, becoming visible by the high expression levels achieved in *N. benthamiana* cells. Some localizations might also be artifactual: for instance, AtUMAMIT11 and 28 were detected in the ER and dots, while previous reports unequivocally showed them being plasma membrane-resident using the same *N. benthamiana* expression system (Müller *et al.*, 2015). Proteins over-expressed in non-native cells are sometimes not efficiently targeted to the correct membrane, because specific effectors are missing in heterologous cells or for other diverse reasons (for example AtUMAMIT24; (Besnard *et al.*, 2018)). In our hands, the expression vector (including promoter, terminator, spacers and cloning approach) also influences the expression level and hence the localization of the fluorescence in *N. benthamiana* cells (G. Pilot, unpublished results). Consequently, subcellular localization inferred from heterologous expression must always be used with care; this study nevertheless provides a first glimpse into the diversity of the localization of the AtUMAMITs in the cell.

**Table 2:**
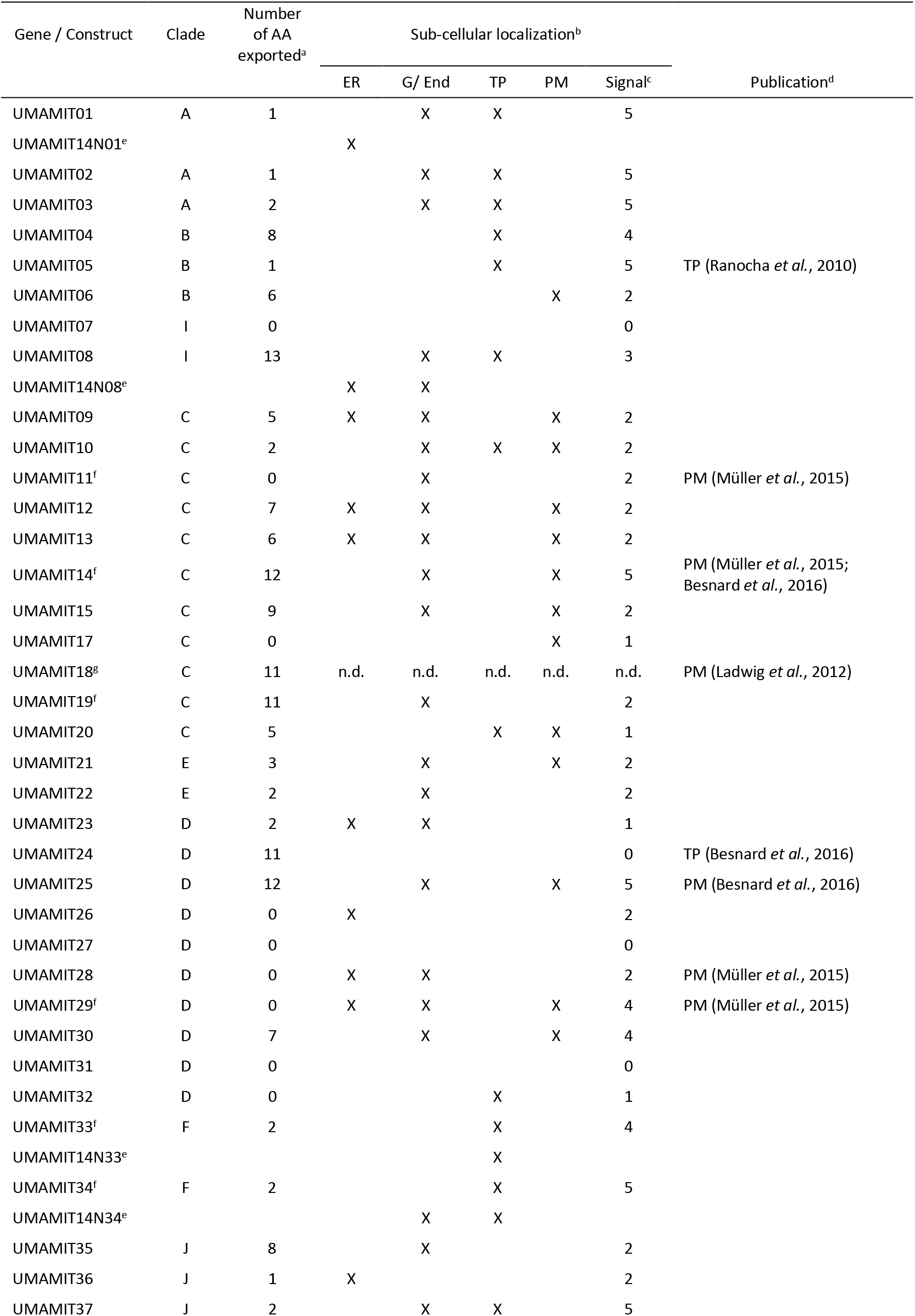
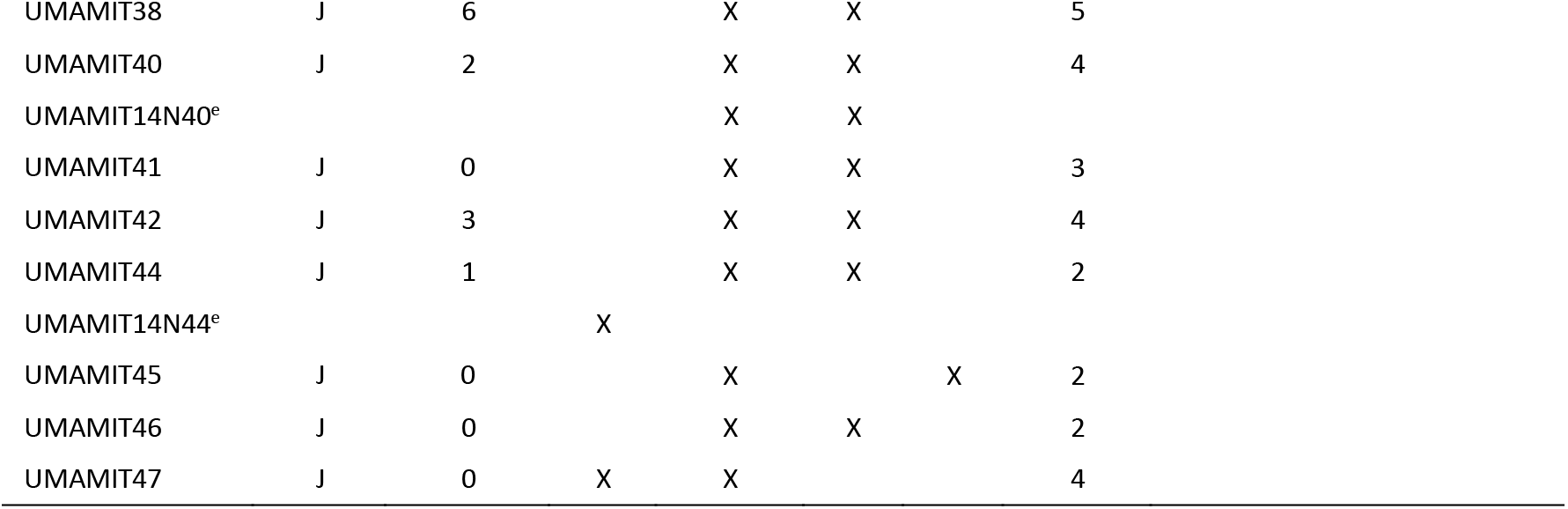
Summary of AtUMAMIT export activity in yeast and of the subcellular localization of the AtUMAMIT-GFP constructs expressed in N. benthamiana epidermis. ^a^ Number of amino acids exported at a level at least 1.5 times the level of the empty plasmid, from yeast export assay. ^b^ ER: endoplasmic reticulum; G/End: Golgi/endosomes; TP: tonoplast; PM: plasma membrane. See Supplemental Figure 12 for pictures. ^c^ Intensity of the GFP signal, on a scale from 0 (no signal) to strong (5). ^d^ Article reporting previous localization result. ^e^ The N-terminus of the AtUMAMIT in question was replaced with that of AtUMAMIT14. See Supplemental Figure 14 for pictures. ^f^ The genomic rather than the cDNA construct has been used for localization. We found that for several genes, including the introns in the CDS improved the signal, possibly by enhancing the expression. ^g^ Localization of AtUMAMIT18 was not tested. n.d.: not determined.

We took advantage of this diversity to probe sub-cellular addressing signals. Several motifs have been identified that target plant membrane proteins to the tonoplast, the di-leucine motif (D/E)X3-5L(L/I) and the tyrosine motif YXXΦ (where Φ is a hydrophobic residue) (Komarova *et al.*, 2012; Wolfenstetter *et al.*, 2012; Pedrazzini *et al.*, 2013). While several of these motifs appeared either in N- or C-termini of AtUMAMITs, no obvious relationship between presence/absence of the motifs and tonoplast/other membrane localization was identified (Supplemental Figure 14). To test whether a tonoplast-targeting signal was present in the N-terminus of some AtUMAMITs, we exchanged the N-termini of these tonoplast-localized AtUMAMIT01, 08, 33, 34, 40 and 44 with that of the plasma membrane-localized AtUMAMIT14. The resulting constructs were expressed in *N. benthamiana* epidermis cells, and the localization was determined by confocal microscopy. Exchanging the N-terminus of AtUMAMIT33 and 40 had no effect on the tonoplastic localization of the proteins (Supplemental Figure 15 and Table 2), suggesting that a tonoplast targeting signal is not present in their N-terminus. On the contrary, the same procedure led to complete retention of AtUMAMIT01, 08, and 44 in the ER and endosomes, and partial localization of AtUMAMIT34 in endosomes (Supplemental Figure 15 and Table 2), which may indicate that a signal for ER exit is present in the N-termini of AtUMAMIT01, 08 and 44, or that two signals, one in the N-terminus and one elsewhere in the protein, are necessary for targeting to the tonoplast, and the loss of one of those signals renders the protein unable to be correctly addressed, remaining in the ER.

### Functional properties of the AtUMAMIT proteins

The functional characterization of the AtUMAMITs reported so far revealed that they are able to export amino acids out of the cytosol (Ladwig *et al.*, 2012; Müller *et al.*, 2015; Besnard *et al.*, 2016; Besnard *et al.*, 2018). To determine which other AtUMAMITs bear this property, the corresponding cDNAs were expressed in the yeast strain 22Δ10α, which is deficient in the uptake of most proteogenic amino acids (Besnard *et al.*, 2016). Amino acid export activity from the cytosol was tested by measuring the amounts of each amino acid secreted in the medium by the cells (Figure 5). AtUMAMIT04, 09, 12, 14, 15, 18, 19 and 25 led to the highest amino acid secretion, on average twice or more the amount secreted by yeast cells containing the empty vector (Supplemental File 4). Several AtUMAMITs exported more than 8 amino acids (*i.e.* AtUMAMIT04, 08, 14, 15, 18, 19, 24, 25 and 35), and some proteins appeared more specific and exported with high efficiency only a few amino acids (*e.g.* Gln for AtUMAMIT09, Asp for AtUMAMIT19 and Arg for AtUMAMIT12; Table 2). PCA analysis of the exported amino acid profiles showed that AtUMAMIT08, 12, 14, 18, 15, 19, 24 and 25 are efficient exporters showing excreted amino acid profiles different from one another and the other proteins (Supplemental Figure 16).

**Figure 5:**
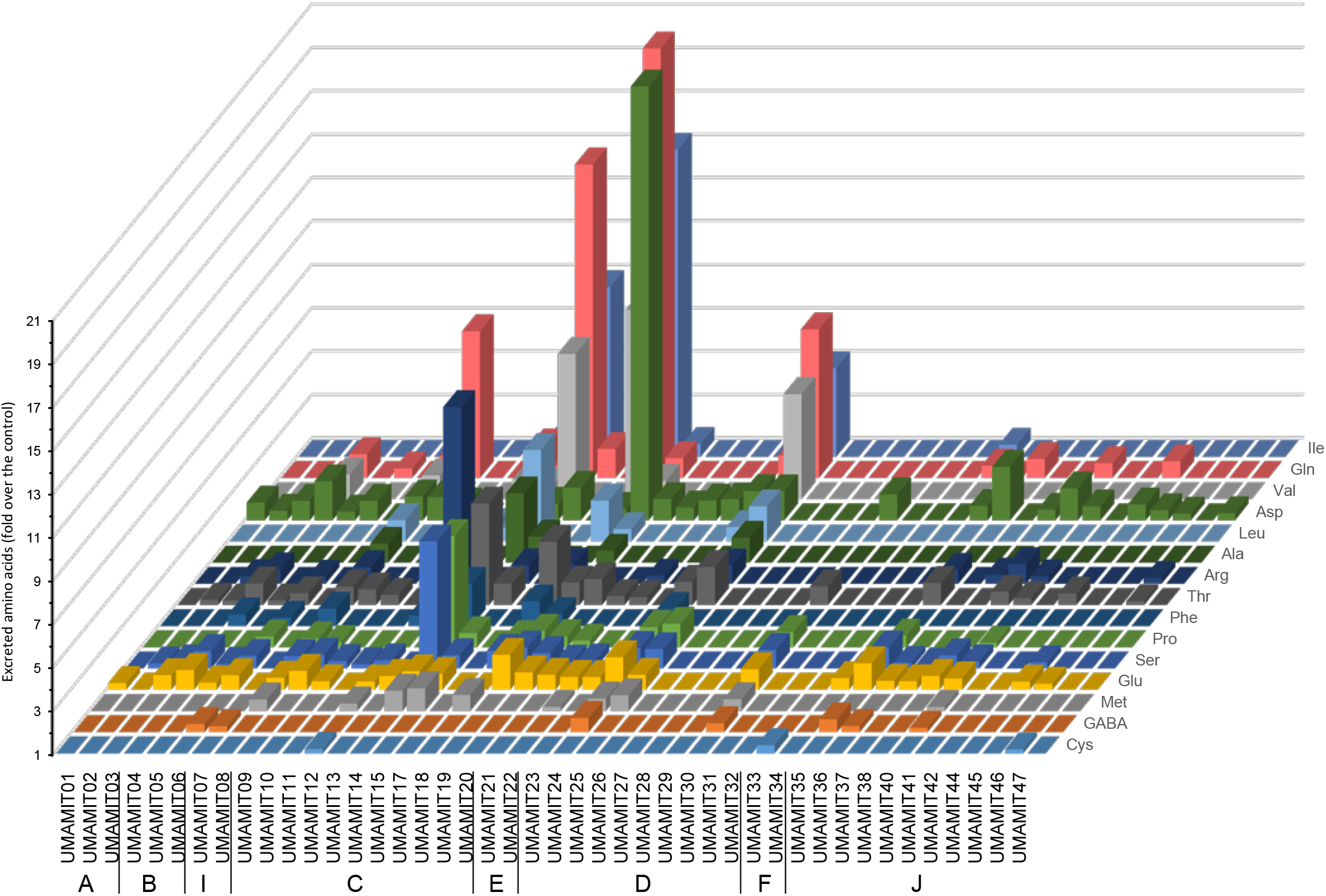
Amino acid excreted by yeast expressing AtUMAMITs. 22Δ10α yeast cells were transformed with plasmids driving the expression of each of the 44 AtUMAMITs, or the empty vector (control). After 22 hours of growth in minimal medium amino acid content in the medium was measured by UPLC. Amino acid excretion is expressed as fold increase relative to cells expressing the empty vector. Only significant changes compared to the empty vector control are shown (t-test, p<0.05, n=4). A to J: clades to which each of the AtUMAMIT belong. See Supplemental File 4 for the data used to create this graphic.

AtUMAMIT14 and 18 were shown to import amino acids at low rates when expressed in *Xenopus* oocytes or yeast (Ladwig *et al.*, 2012; Müller *et al.*, 2015). To test whether other AtUMAMITs would display uptake activity, uptake was measured in 22Δ10α by the classical yeast complementation assay, in which amino acids (3 mM Asn, Gln, Ile, Leu, Met, Phe, Thr, Val, and 6 mM Glu) were provided as the sole nitrogen source. Contrary to the previous yeast strains mentioned above, 22Δ10α is unable to import γ-aminobutyric acid, citrulline, ornithine and 19 of the 20 proteogenic amino acids (Besnard *et al.*, 2016) making it the tool of choice to characterize amino acid transporters. Unfortunately, no uptake activity of any of the nine amino acids tested was detected in this assay (see an example for Met in Supplemental Figure 17), even for AtUMAMIT14 and 18, proven to mediate import in some conditions (Ladwig *et al.*, 2012; Müller *et al.*, 2015). We then used a more sensitive method than the yeast complementation assay to test for import activity. Ten AtUMAMITs were chosen from different clades (AtUMAMIT01, 05, 14, 18, 19, 21, 24, 28, 29 and 35), not paying attention to the putative subcellular localization in plants, and uptake of radiolabeled Al and Gln by cells expressing these AtUMAMITs was measured over time. Uptake of Ala by AtUMAMIT01, and of Ala and Gln by AtUMAMIT18 was observed, but not for any of the other tested proteins (Figure 6). Our results expand the set of AtUMAMITs demonstrated to export amino acids, and confirms that at least some can also import amino acids. Whether these properties can be generalized to the whole UMAMIT family remains to be determined.

**Figure 6:**
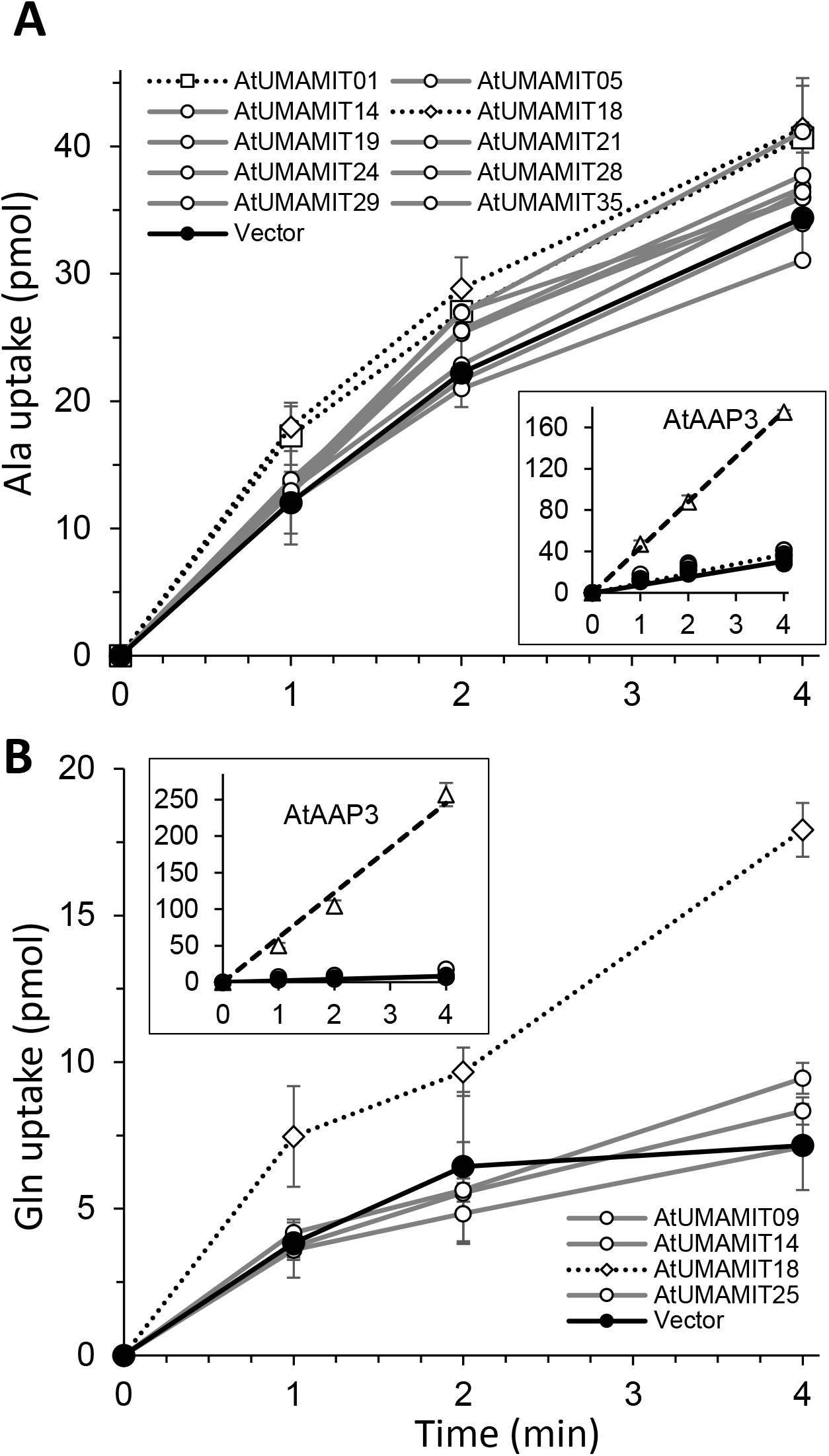
Amino acid uptake activity of selected AtUMAMIT proteins. A. 1 mM Ala uptake. B. 1 mM Gln uptake. The empty vector control is indicated by a solid black line (Vector). The dotted lines correspond to proteins for which the uptake was significantly different from the control at all time points (t-test, p<0.05, n=3 biological replicates). Insets: uptake curve of the importer AtAAP3, used as a positive control (dashed lines), was added to the graph, to emphasize the difference in uptake rate between a proton-coupled importer and the AtUMAMITs.

### Effect of point mutations on the transport activity of AtUMAMIT14

The multiple sequence alignments of the 180 UMAMITs and of the structural domains (Supplemental Figures 1 and 5) identified conserved motifs in structural domains A and/or B. To test the importance of these residues for the function of the UMAMITs, the following mutations were performed on AtUMAMIT14, the most active AtUMAMIT in yeast export assays: G33V, K134E, conserved in structural domain A only, and E122R, G315V and G144V conserved in both domains; the double mutation G144V + G315V was also performed. The corresponding mutagenized DNA sequences were expressed in 22Δ10α, and the effect of the substitutions on amino acid secretion by AtUMAMIT14 into the medium was measured by UPLC. Substitution G33V and K134E respectively led to ~35% and ~50% reductions in the total amount of amino acids specifically secreted by AtUMAMIT14 (*i.e.* after subtraction of the amounts secreted by vector-expressing cells); similar reductions were observed in the amounts of Gln and other amino acid secreted (Figure 7). The effect of the other mutations, alone or in combination, was more drastic: Gln secretion was completely abolished, and the secretion of other amino acids was reduced by ~90%. It is interesting to note that the mutations with the stronger effect correspond to amino acids found in both structural repeats, near or in TMs A5 or B5 (Supplemental Figure 6). To ensure that the absence of export activity was not caused by loss of expression of the mutated proteins, wild type and variant AtUMAMIT14 proteins were expressed in yeast fused to the GFP in their C-termini. GFP fluorescence was detected for all proteins, with the non-functional variants leading to larger aggregates in the cells and brighter fluorescence (Supplemental Figure 18). This result suggests that the non-functional variants are expressed in yeast, but that the mutations lead to defects in stability or trafficking.

**Figure 7:**
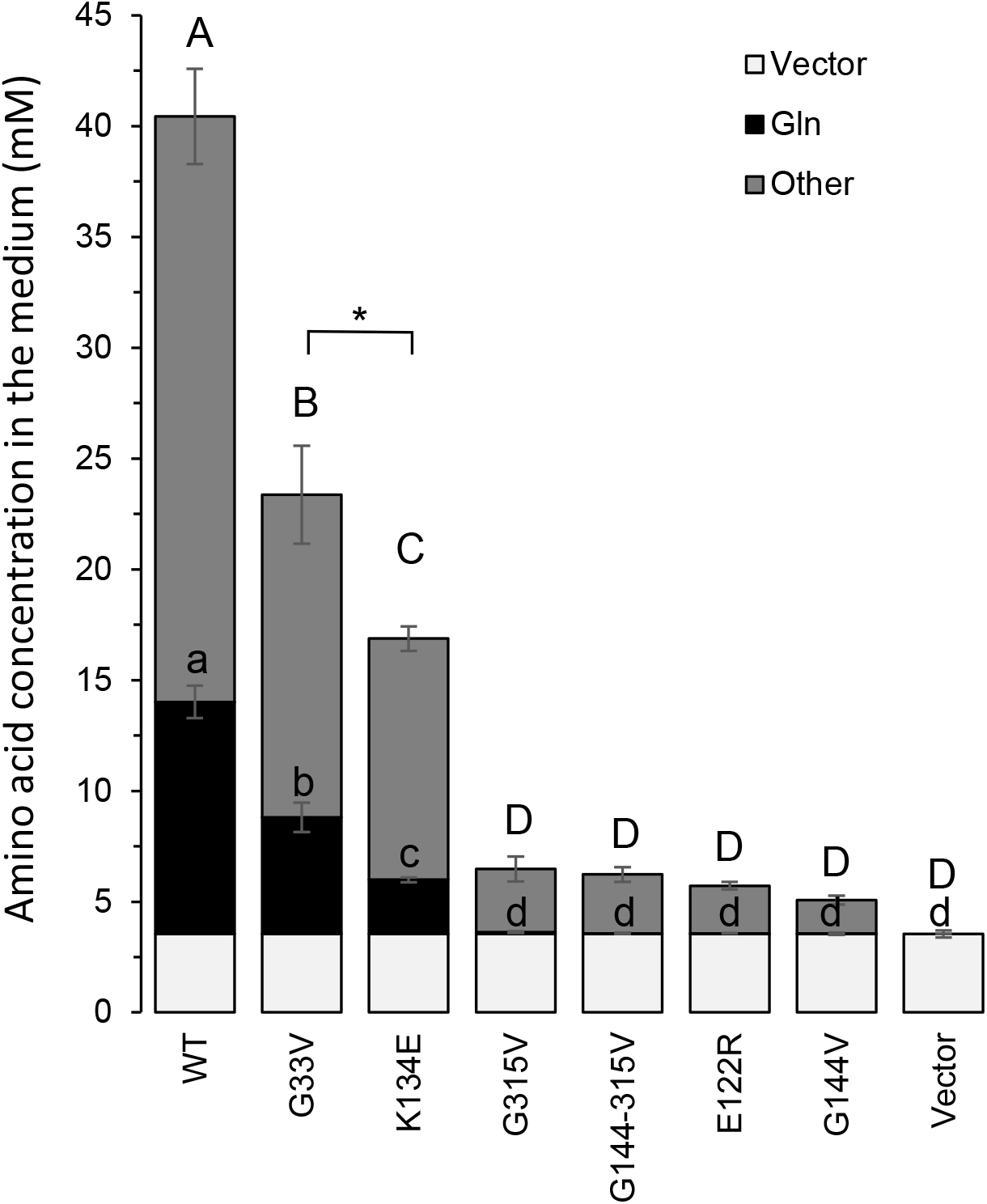
Amino acid excreted by yeast expressing the AtUMAMIT14 mutants. Similar experiment as Figure 5. Amino acid contents excreted by the vector-containing cells are indicated in light grey. The total amount of amino acids secreted by the vector-containing cells is shown by gray bars (Vector). The amounts of Gln and the other amino acids secreted by cells expressing the AtUMAMIT14 proteins in addition to the amounts of that of the vector are represented by black and dark grey bars respectively. ANOVA Tuckey HSD (n=3 biological replicates): lowercase letters correspond to the differences between the samples for Gln content (p<0.0001); uppercase letters correspond to the differences for the sum of the other amino acids (p<0.0001, except *=0.02).

## Discussion

In this study, we wanted to characterize the members of the UMAMIT family using an unbiased, systematic approach involving phylogenic reconstruction, homology modeling, mining expression data, determination of the expression in the plant and the cell, and functional characterization in yeast. The bulk of data can be used as a reference for researchers interested in choosing specific UMAMITs to study, or to characterize UMAMIT genes identified from forward genetic screenings, and quantitative trait locus, genome wide association mapping or transcriptomics analyses, from model organisms or crops. In addition this study enabled to draw hypotheses and conclusion that would open new research avenues on this family and membrane transporters, *e.g.* on their evolution, structure, trafficking or functional properties.

### Most UMAMITs display amino acid exporter properties

Currently, eight AtUMAMITs have been shown to export amino acids (AtUMAMIT11, 14, 18, 23, 24, 25, 28 and 29) (Ladwig *et al.*, 2012; Müller *et al.*, 2015; Besnard *et al.*, 2016; Besnard *et al.*, 2018), and it remained unclear if this characteristic was common to all members of the family. Secretion of amino acids by yeast expressing most AtUMAMITs was detected, except for AtUMAMIT17, 26, 27, 28, 29, 31, 32 and 41 (genes mostly from Clade D). Genes with the strongest export activities in yeast are from Clade C (Chi-SQ p-value=0.032), which is also enriched in proteins that are located at the plasma membrane (Chi-SQ p-value=0.013). Overall, proteins addressed to the plasma membrane display more detectable export activity (Chi-SQ p-value=0.004), suggesting that the yeast assay introduces a bias towards plasma-membrane localized protein, keeping in mind that this bias may originate from the assumption that UMAMITs preferentially address to the same membrane in yeast as in plants. Based on our results, it seems reasonable to predict that most, if not all, AtUMAMITs mediate amino acid export when expressed in a system enabling the measurement of their transport activity. The fact that yeast expressing AtUMAMIT26, 27, 28 and 29 exported less Cys, GABA and Met than the empty vector-expressing yeasts was puzzling, suggesting that these proteins import those amino acids or might be inserted in the membrane in the inverse orientation. Alternatively, their expression in internal yeast membranes could disrupt yeast amino acid metabolism, affecting the cytosolic content and the concentration gradient of these compounds across the membrane, leading to enhanced secretion of amino acid by endogenous amino acid exporter(s) (Velasco *et al.*, 2004) to a higher level than the empty vector-expressing cells.

AtUMAMIT14 and 18 have been shown to display import in addition to export activities, suggesting they facilitate transport of amino acids (Gln, Glu and Citrulline, and His, Asp and Gln, respectively) across membranes (Ladwig *et al.*, 2012; Müller *et al.*, 2015). Our attempt to detect import relied mainly on yeast complementation but using 22Δ10α, a yeast strain different from the ones used in those reports (JT16, 22Δ8AA, 23344c, 30.537a, 22574d). While our yeast complementation assay was most likely not sensitive enough to measure uptake, direct uptake of Ala or Gln could not be detected for most tested AtUMAMITs. To grow, 22Δ10α requires a substantial uptake of amino acids to sustain nitrogen supply to yeast metabolism, while JT16 could grow with only a minute uptake of His to complement the His auxotrophy, since its nitrogen source comes from ammonium in the medium. In addition, the amino acids (Citrulline, Asp, His or Glu) and concentrations (10 μM or 10 mM Gln) employed for characterizing AtUMAMIT14 and 18 (Ladwig *et al.*, 2012; Müller *et al.*, 2015) were different than here, which could explain the apparent discrepancies between the observations. We also noticed that the growth conditions of the yeast cells (*e.g.* rich vs. minimal medium) affect measured uptake rates, possibly by modifying the intracellular amino acid pools: the transport rate by a facilitator typically depends on the electrochemical gradient, which would greatly be influenced by a modified cytosolic amino acid content. Whether the amino acid import detected for AtUMAMIT01, 14 and 18 can be generalized to the whole UMAMIT family remains to be demonstrated using standardized procedures across laboratories. On the contrary, the amino acid electrochemical gradient is in favor of export towards the medium if the latter is chosen appropriately, which enables more sensitive measurement of export but prevents any determination of substrate affinity, since the intracellular concentration of amino acid cannot be set experimentally. This could explain why we detected an export activity for almost all AtUMAMITs, contrary to an import activity. Finally, we have only tested amino acids as substrate. Since transporters from the AAAP and UMAMIT families can transport non-amino acid substrates (Yang *et al.*, 2006; Ranocha *et al.*, 2013; Shin *et al.*, 2015; Jiang *et al.*, 2018; Choi *et al.*, 2019), one should keep in mind that the physiological substrate of some UMAMITs might be hormones or amino acid-containing metabolites, leading to false negative transport results.

Careful comparison of the protein sequences of the closely related AtUMAMITs that display different amino acid export patterns, *e.g.* AtUMAMIT14/ 15 (Leu, Ile, Val), and 18/19 (Asp, Gln), could be used to identify substrate-binding residues positioned inside the homology model. Followed by site-directed mutagenesis and transport assays, these *in silico* results would help determine the structure-function relationships of the UMAMITs and test the validity of the homology model.

### Hypothetical roles of AtUMAMIT using the data produced by this analysis

We propose that UMAMITs would be involved in various functions in amino acid transport. Behaving as facilitators, they enable flow of amino acids downstream from their electrochemical gradient. At the cell level, this would translate into passive export of amino acid from the cytosol to various compartments: to the vacuole lumen, possibly for storing excess amino acids since the concentration of amino acids in the vacuole is typically 10-100 times lower than in the cytosol (Riens *et al.*, 1991; Weiner and Heldt, 1992; Winter *et al.*, 1993, 1994); to the apoplasm where amino acids are about 10 times less concentrated than in the cytosol (Lohaus *et al.*, 2001) for cell-to-tell translocation.

The data provided from this study could be useful for research in membrane protein trafficking processes; for instance, the subcellular localization data could help identify tonoplast targeting signals. Mining those data would also help identify UMAMITs with redundant functions that cannot be easily predicted by sequence similarity, which could help choosing double or triple mutants to study in order to identify their role in the plant. This extensive dataset would also be useful to pick AtUMAMITs putatively involved in a specific function. One might be interested in characterizing the actors of the transport of amino acids to the vacuole in seeds, or roots, or export from cell layers to the phloem. Here, we provide some simple hypotheses generated from our results concerning the role of a few AtUMAMITs (Tables 1 and 2, Figure 4).

Case 1. AtUMAMIT01 and 02 are from the ancient Clade A, the proteins are addressed to the tonoplast, they show low change in mRNA accumulation in response to various treatments, and are expressed in seeds. Hypothesis: these two genes fulfill a role common to all land plants, related to reproduction, maybe storage of amino acids in the vacuole of the embryo. A double mutant might show defects in embryo development, germination or seed protein content.

Case 2: AtUMAMIT05 and 41 are addressed to the tonoplast, expressed at high levels in most plant organs, but their expression is insensitive to growth conditions. Hypothesis: these two genes are involved in central amino acid homeostasis common to all cells, possibly involved in equilibration of amino acid between cytosol and vacuole, and can be considered housekeeping transporters. The double mutant might display decreased fitness, maybe exacerbated in conditions where amino acid remobilization from/transfer to the vacuole is critical, *e.g.* low or high nitrogen availability.

Case 3: AtUMAMIT19 and 24 are expressed at low levels in the plant (in the root stele for AtUMAMIT19), but respond extensively to various stresses. Hypothesis: they are involved in acclimation to new growth conditions, a transition that possibly involves adjustment of central metabolic activity. Note that AtUMAMIT24 has also been shown to have a role in seed nutrition (Besnard *et al.*, 2018), suggesting that some genes can be involved in stress response and have a different role during non-stress conditions.

### Study of the structure of the UMAMITs gives a glimpse into the evolution of family

In addition to bringing some clarity in the diversity of UMAMIT sequences by grouping the genes in clades, the phylogenic analysis enabled drawing some hypotheses on the evolution of the family in plants. The presence of UMAMITs from the lower plants *Selaginella* and *Physcomitrella* only in clade A suggests that the genes from this clade have not diverged much from their common ancestor, and may have retained some of their original functional properties. Our systematic analysis did not reveal any specific characteristic for the Clade A members AtUMAMIT01, 02 and 03 compared to the other AtUMAMITs, but the differences may lie in other parameters that we have not tested.

As expected, monocots, dicots and conifers made their own subclades within the major clades, which was one of the criteria used to define the clades A through J. All dicot families used to build the tree were present in each clade, with the notable exception of clade G: *Brassicaceae* and conifers were absent from this clade. *Brassicaceae* are known for being a non-host of symbiosis with fungi through arbuscular mycorrhiza (Delaux *et al.*, 2014), while *Picea* and *Pinus* associate with fungi through another symbiosis, the ectomycorrhization (Garcia *et al.*, 2015). Looking more closely, the *Brassicales* species *Carica papaya*, which can develop arbuscular mycorrhiza, contained one UMAMIT in this clade (Delaux *et al.*, 2014). This observation highlights a correlation between the ability to develop arbuscular mycorrhiza and the presence of UMAMIT from clade G. It would be interesting to test the effect of loss-of-function of the corresponding *Medicago* gene MtrunA17_Chr1g0171301 on its ability to develop, sustain or benefit from arbuscular mycorrhization.

The sequence similarity of AtUMAMITs from Clade J, their position in clusters in the Arabidopsis genome and the fact that this clade is constituted exclusively from dicot sequences, all together suggest that this clade diversified recently. Species from the *Rosids*, *Malvids*, and *Favids* showed a higher proportion of genes in this clade than was expected if the distribution was homogenous across species (x1.3 to x1.6 fold enrichment). The limited availability of expression data of the genes from this clade (most of the corresponding Arabidopsis genes were not present on the Affymetrix chip) makes formulating any hypothesis difficult.

### Functional UMAMITs might be dimers

The study of the structure of UMAMIT proteins suggested that they are constituted of two repeated, inverted 5 TMs regions, in good agreement with the evolution model proposed for the DMT proteins (Jack *et al.*, 2001). This result also agrees with current views on the evolution of molecular transporters which are often hypothesized to arise from the duplication and fusion of genes encoding monomers (Duran and Meiler, 2013; Forrest, 2015; Drew and Boudker, 2016). Similar to the proteins used for the homology modeling and the EmrE protein from the DMT superfamily (Fleishman *et al.*, 2006), UMAMITs were predicted to display a symmetry around an axis running lengthwise, parallel to the plane of the membrane. Yet, the helices of each of the two inverted structural domains are not grouped together: A1 and A2 helices are grouped with the B3, B4 and B5 helices (and vice-versa for the other helices), with the “cross-over” happening at the level of the A2 and B2 domains, located on the other side of the A5 and B5 domain (Figure 4).

The SwissModel server used, among other templates, the crystal structure of the dimer of GDP-mannose transporter to build the structure of AtUMAMIT14, leading to a model composed of a UMAMIT dimer in which the A5 and B5 helices are located at the interface between the two monomers. Results from homology modeling always need to be confirmed experimentally. Our mutagenesis approach showed that the conserved Gly144 and 315, but not of K134, from these helices suppressed the activity of the transporter. On the model, the two Gly are facing the helix of the same monomer, while the Lys is facing the inside of the monomer. Gly in transmembrane helices is often involved in helix-helix interaction necessary for helix packing by alternating with Val, Leu or Ile residues, similar to a zipper (Walters and DeGrado, 2006; Li *et al.*, 2012). The G144V and G315V mutations may thus disrupt the structure of AtUMAMIT14, rendering the monomers non-functional, or disrupt the interaction between two monomers. More work is needed to resolve the question of the structure of the AtUMAMITs and whether they can form heterodimers which could potentially modify they localization and transport properties.

## Material and methods

### Plant transformation, growth and staining

Arabidopsis (Col-0) was stably transformed by the floral dip method (Clough and Bent, 1998), using *Agrobacterium tumefaciens* GV3101 (pMP90). Transgenic AtUMAMITpromoter-GUS lines were grown on soil (Sungrow mix, composed of peat and perlite), in an incubator (120 μE.m^−2^.s^−1^, 22°C 16 h light / 8 h dark) for 5 weeks, before the organs were harvested for immediate histochemical staining performed as described (Lagarde *et al.*, 1996). For ensuring penetration of the fixing and staining solutions within thick tissues, leaves, stems and siliques were cut in 5-7 mm pieces before reaction. Surface-sterilized seeds were grown on J medium (Besnard *et al.*, 2016) or half strength Murashige and Skoog medium, both supplemented with 1% sucrose and 0.9% agar. Seeds were stratified for three days at 4°C, and moved to a growth chamber with constant light (125 μE.m^−2^.s^−1^). Histochemical reactions were performed in 12 well plates, at 37°C for times varying between genes and organs, to ensure sufficient but not excessive staining. After staining, about 5 mm of stem segments were embedded in 5% agarose, cut in 60-70 μm sections on a Leica VT1200 vibrating microtome. All stained organs were then equilibrated in 60% glycerol, and examined and photographed with an Axio Imager A1 microscope (Zeiss).

### Subcellular localization

AtUMAMIT-GFP constructs were introduced and expressed in leaves of 5-week-old *Nicotiana benthamiana* plants as described (Batoko *et al.*, 2000). GFP fluorescence in the epidermal cells was localized three days after infiltration as previously described (Pratelli *et al.*, 2012).

### Phylogenetic analysis

To construct the 180 taxa tree, sequences of the UMAMITs were retrieved in 2014 from various databases, including TAIR (arabidopsis.org), Phytozome (phytozome.jgi.doe.gov), and the Rice Genome Annotation Project (rice.plantbiology.msu.edu), using several Arabidopsis proteins as initial query for BLAST searches. The sequences were aligned with MUSCLE in the MEGA6 package with default parameters (Tamura *et al.*, 2011), the N and C termini, as well as columns that contained more than 70% gaps, and incomplete protein (pseudogenes, truncated or proteins missing internal sequences) were manually removed from the alignment. The sequences of the 1468 taxa tree were retrieved from Phytozome by BLAST searches using proteins from various species and clades out of the 180 sequence set. The optimum substitution model was searched using modeltest in MEGA6; the Jones-Taylor-Thornton model and the Gamma distribution for modeling the non-uniformity of evolutionary rates among sites were chosen. Maximum likelihood phylogenic analysis was computed on a supercomputer using RaXML (Stamatakis, 2014). Bootstrap analysis was performed using the rapid bootstrap option, and 10,000 or 1,000 repetitions for the 180 and 1468 taxa trees, respectively.

### Clonings

The coding sequences of most of the 44 non-pseudogene AtUMAMIT were amplified by PCR (oligonucleotide sequence is provided in Supplemental Table 1) by using the KOD Hot Start DNA Polymerase kit (Novagen) from a mix of reverse-transcribed RNA extracted from seedlings and inflorescences. The cDNA of AtUMAMIT04, 09, 13, 24, 26, 34 and 37 were amplified either by PCR from full-length clones, using nested PCR or by splicing by overlap extension (SOE) PCR (details available upon request). Genomic sequences (ATG-stop) of AtUMAMIT04, 11, 12, 14, 18, 19, 29, 33 and 34 were cloned similarly using the same primers from Col-0 genomic DNA. PCR products were gel-purified and cloned into pDONRZeof1 by Gateway recombination (Invitrogen). All sequences were confirmed before further cloning. About 1.5 kbp of the region upstream from the ATG of chosen AtUMAMITs were cloned similarly by Gateway cloning. cDNAs (with a stop codon) were Gateway cloned into the yeast expression vector pDR196-W (Loque *et al.*, 2007). cDNA and genomic sequences (without stop codon) were Gateway-cloned into the plant expression vector pPWGTkan (Pratelli *et al.*, 2012) or pC4H-W-YFP (Liang *et al.*, 2017). Promoter sequences were Gateway-cloned into the gateway derivative of pUTkan (Pratelli *et al.*, 2010). Site-directed mutagenesis of the AtUMAMIT14 cDNA cloned into pDONRZeof1 was performed using the Kunkel method as described (Loque *et al.*, 2007). The exchange of the N-terminal region of AtUMAMIT01, 08, 10, 33, 34, 40 and 44 with that of AtUMAMIT14 was performed by SOE PCR using oligonucleotides from Supplemental Table 1.

### Yeast complementation, uptake, and secretion assay

Yeast strain 22Δ10α (MATα *gap1-1 put4-1 uga4-1 can1::HisG lyp1-alp1::HisG hip1::HisG dip5::HisG gnp1Δ agp1Δ ura3-1*) (Besnard *et al.*, 2016) was transformed with the AtUMAMIT cDNAs cloned into pDR196-W using the PEG/LiAc method (Gietz and Schiestl, 2007). Because of the large number of proteins to be assayed for transport properties, >50 colonies obtained from a transformation event were pooled and treated as one replicate, rather than using single colonies. For the plate complementation assay, transformed yeast cells were grown at 30°C for 3 days; cells from one replicate were resuspended in 500 μl sterile water and 10-fold serial dilutions were made in water. Five μl of each dilution were then spotted on solid, freshly made minimal medium (Jacobs *et al.*, 1980) without (NH_4_)_2_SO_4_, supplemented with 3 mM of a single amino acid as sole nitrogen source. For radioactive amino acid uptake assays, one replicate of 22Δ10α cells from pooled colonies were grown at 30°C in 4 ml SC medium lacking uracil for about 16 h and grown in YPD medium containing adenine at a starting OD_600_ of 0.1. Cells were harvested by centrifugation at 2,500 g for 5 minutes when OD_600_ reached 0.4, washed twice with sterile water, and resuspended in Buffer A (50 mM KH2PO4, 600 mM sorbitol) to an OD_600_ of 5. Yeast cells were then aliquoted in 100 μl and kept on ice. Ten μl of 1 M glucose was added to the aliquots, and cells were incubated 30°C for 5 minutes in an Eppendorf thermomixer. Uptake was started by mixing three aliquots of cells with 110 μl of uptake solution (2 mM cold amino acid and 1 μl of ^3^H-labeled amino acid in Buffer A). At each time point, 50 μl of cells were taken, dropped on glass filters, and washed twice with 5 ml of cold Buffer A by vacuum filtration. Filters were put into scintillation vials, covered with 5 ml of Ultima Gold XR scintillation cocktail (Perkin Elmer), and the radioactivity was counted by using a LS6500 multi-purpose scintillation counter (Beckman). For the secretion assay, four replicates of pooled yeast colonies were grown in 2 ml liquid minimum medium containing 2.5 mM (NH_4_)_2_SO_4_ at 30°C at a starting OD_600_ of 0.05 for 22 h. OD_600_ of yeast cultures were measured and supernatants collected by running the cultures through a Millipore filter using MultiScreen HTS Vacuum Manifold (Millipore) first, and then through a 10 kDa exclusion membrane (Ultracell 10, Millipore) by centrifugation at 3000 g for 90 min at room temperature. Amino acids in the medium were detected by UPLC and quantified as described (Collakova *et al.*, 2013).

### Data analysis and visualization

Public expression datasets were retrieved from http://bar.utoronto.ca/ (Toufighi *et al.*, 2005). Statistical and principal component analyses and violin graphs were performed in JMP Pro 15 or Microsoft Excel. Phylogenic trees were visualized and edited using FigTree 1.4.2 (tree.bio.ed.ac.uk/software/figtree/), and exported to Adobe Illustrator for shading. Pictures of the molecular models were created using UCSF Chimera 1.14.

## Supplemental data

Fig. S1: Multiple sequence alignment of 180 UMAMIT sequences

Fig. S2: Distribution of the mRNA contents of the AtUMAMIT genes in Arabidopsis in various conditions and organs.

Fig. S3: Position of the AtUMAMITs on the Arabidopsis chromosomes

Fig. S4: Phylogenic tree of 1469 UMAMIT protein sequences

Fig. S5: Alignment of the two structural repeats of AtUMAMIT and OsUMAMIT protein sequences

Fig. S6: Model of the AtUMAMIT14 predicted protein dimer

Fig. S7: PCA analysis of the localization of the expression of the AtUMAMIT in the plant

Fig. S8: Expression pattern of 15 AtUMAMIT promoters

Fig. S9: Localization in the plant of 13 AtUMAMIT genes, obtained from translatome data

Fig. S10: PCA analysis of the change in expression of the AtUMAMIT in the plant.

Fig. S11: Correlation between change in mRNA expression and expression level of the AtUMAMITs

Fig. S12: Localization of GUS activity of plants expressing GUS under the control of AtUMAMIT promoters

Fig. S13: Subcellular localization of the AtUMAMITs proteins expressed in *N. benthamiana* cells

Fig. S14: Alignment of the N- and C-termini of the AtUMAMITs

Fig. S15: Subcellular localization of some AtUMAMIT proteins without and with N-terminus exchange

Fig. S16: PCA analysis of amino acid export profiles of the AtUMAMIT expressed in yeast

Fig. S17: Methionine uptake complementation assay of yeast expressing the AtUMAMITs

Fig. S18: Expression and localization of wild type and variants of AtUMAMIT14-GFP expressed in yeast cells

Supplementary File 1: Sequence information of the UMAMITs used in the article.

Supplementary File 2: Position of the introns in the *UMAMIT* genes used in the phylogenic tree from Figure 1.

Supplementary File 3: AtGenExpress data for the expression of the UMAMT genes present on the ATH1 Affymetrix microarray chip.

Supplementary File 4: Amounts of secreted amino acids from 22Δ10α yeast cells expressing each of the AtUMAMITs

## Acknowledgements

This work was supported by the School of Plant and Environmental Sciences (Virginia Tech), NSF-IOS (1353366), NSF-MCB (1052048), and the Hatch Program of the National Institute of Food and Agriculture (VA-135908) and the Virginia Agricultural Experiment Station.

## Author contribution

Chengsong Zhao: Investigation, Writing - original draft, Methodology; Réjane Pratelli: Investigation; Shi Yu: Investigation; Brett Shelley: Investigation; Eva Collakova: Investigation; Guillaume Pilot: Investigation, Conceptualization, Writing – review & editing, Supervision, Funding acquisition.

## Data Availability Statement

All data supporting the findings of this study are available within the paper and within its supplementary materials published online, and/or available upon request from the corresponding author (Guillaume Pilot).

## References

Batoko H, Zheng HQ, Hawes C, Moore I. 2000. A rab1 GTPase is required for transport between the endoplasmic reticulum and golgi apparatus and for normal golgi movement in plants. The Plant Cell 12, 2201–2218.

Besnard J, Pratelli R, Zhao C, Sonawala U, Collakova E, Pilot G, Okumoto S. 2016. UMAMIT14 is an amino acid exporter involved in phloem unloading in Arabidopsis roots. Journal of Experimental Botany 67, 6385–6397.

Besnard J, Zhao C, Avice JC, Vitha S, Hyodo A, Pilot G, Okumoto S. 2018. Arabidopsis UMAMIT24 and 25 are amino acid exporters involved in seed loading. Journal of Experimental Botany 69, 5221–5232.

Brady SM, Orlando DA, Lee JY, Wang JY, Koch J, Dinneny JR, Mace D, Ohler U, Benfey PN. 2007. A high-resolution root spatiotemporal map reveals dominant expression patterns. Science 318, 801–806.

Busov VB, Johannes E, Whetten RW, Sederoff RR, Spiker SL, Lanz-Garcia C, Goldfarb B. 2004. An auxin-inducible gene from loblolly pine (Pinus taeda L.) is differentially expressed in mature and juvenile-phase shoots and encodes a putative transmembrane protein. Planta 218, 916–927.

Cartwright DA, Brady SM, Orlando DA, Sturmfels B, Benfey PN. 2009. Reconstructing spatiotemporal gene expression data from partial observations. Bioinformatics 25, 2581–2587.

Choi J, Eom S, Shin K, Lee R-A, Choi S, Lee J-H, Lee S, Soh M-S. 2019. Identification of Lysine Histidine Transporter 2 as an 1-Aminocyclopropane Carboxylic Acid Transporter in Arabidopsis thaliana by Transgenic Complementation Approach. Frontiers in Plant Science 10.

Clough SJ, Bent AF. 1998. Floral dip: a simplified method for *Agrobacterium*-mediated transformation of *Arabidopsis thaliana*. The Plant Journal 16, 735–743.

Collakova E, Aghamirzaie D, Fang Y, Klumas C, Tabataba F, Kakumanu A, Myers E, Heath LS, Grene R. 2013. Metabolic and Transcriptional Reprogramming in Developing Soybean (Glycine max) Embryos. Metabolites 3, 347–372.

Delaux PM, Varala K, Edger PP, Coruzzi GM, Pires JC, Ane JM. 2014. Comparative phylogenomics uncovers the impact of symbiotic associations on host genome evolution. PloS Genetics 10, e1004487.

Denancé N, Ranocha P, Martinez Y, Sundberg B, Goffner D. 2014a. Light-regulated compensation of wat1(walls are thin1) growth and secondary cell wall phenotypes is auxin-independent. Plant Signaling and Behavior 5, 1302–1304.

Denancé N, Ranocha P, Oria N, Barlet X, Riviere MP, Yadeta KA, Hoffmann L, Perreau F, Clement G, Maia-Grondard A, van den Berg GC, Savelli B, Fournier S, Aubert Y, Pelletier S, Thomma BP, Molina A, Jouanin L, Marco Y, Goffner D. 2013. Arabidopsis wat1 (walls are thin1)-mediated resistance to the bacterial vascular pathogen, Ralstonia solanacearum, is accompanied by cross-regulation of salicylic acid and tryptophan metabolism. The Plant Journal 73, 225–239.

Denancé N, Szurek B, Noel LD. 2014b. Emerging functions of nodulin-like proteins in non-nodulating plant species. Plant and Cell Physiology 55, 469–474.

Drew D, Boudker O. 2016. Shared Molecular Mechanisms of Membrane Transporters. Annual Review of Biochemistry 85, 543–572.

Duran AM, Meiler J. 2013. Inverted topologies in membrane proteins: a mini-review. Computational & Structural Biotechnology Journal 8, e201308004.

Fleishman SJ, Harrington SE, Enosh A, Halperin D, Tate CG, Ben-Tal N. 2006. Quasi-symmetry in the cryo-EM structure of EmrE provides the key to modeling its transmembrane domain. Journal of Molecular Biology 364, 54–67.

Forrest LR. 2015. Structural Symmetry in Membrane Proteins. Annual Reviews of Biophysics 44, 311–337.

Gamas P, Niebel Fde C, Lescure N, Cullimore J. 1996. Use of a subtractive hybridization approach to identify new Medicago truncatula genes induced during root nodule development. Molecular Plant-Microbe Interactions 9, 233–242.

Garcia K, Delaux PM, Cope KR, Ane JM. 2015. Molecular signals required for the establishment and maintenance of ectomycorrhizal symbioses. New Phytologist 208, 79–87.

Gietz RD, Schiestl RH. 2007. High-efficiency yeast transformation using the LiAc/SS carrier DNA/PEG method. Nature Protocols 2, 31–34.

Heller G, Adomas A, Li G, Osborne J, van Zyl L, Sederoff R, Finlay RD, Stenlid J, Asiegbu FO. 2008. Transcriptional analysis of Pinus sylvestris roots challenged with the ectomycorrhizal fungus Laccaria bicolor. BMC Plant Biology 8, 19.

Heller G, Lunden K, Finlay RD, Asiegbu FO, Elfstrand M. 2012. Expression analysis of Clavata1-like and Nodulin21-like genes from Pinus sylvestris during ectomycorrhiza formation. Mycorrhiza 22, 271–277.

Jack DL, Yang NM, Saier MHJr., 2001. The drug/metabolite transporter superfamily. European Journal of Biochemistry 268, 3620–3639.

Jacobs P, Jauniaux JC, Grenson M. 1980. A cis-dominant regulatory mutation linked to the argB-argC gene cluster in Saccharomyces cerevisiae. Journal of Molecular Biology 139, 691–704.

Jiang X, Xie Y, Ren Z, Ganeteg U, Lin F, Zhao C, Xu H. 2018. Design of a New Glutamine-Fipronil Conjugate with alpha-Amino Acid Function and Its Uptake by A. thaliana Lysine Histidine Transporter 1 ( AtLHT1). Jounral of Agricultural and Food Chemistry 66, 7597–7605.

Kall L, Krogh A, Sonnhammer EL. 2007. Advantages of combined transmembrane topology and signal peptide prediction--the Phobius web server. Nucleic Acids Research 35, W429–432.

Kelley LA, Mezulis S, Yates CM, Wass MN, Sternberg MJ. 2015. The Phyre2 web portal for protein modeling, prediction and analysis. Nature Protocols 10, 845–858.

Komarova NY, Meier S, Meier A, Grotemeyer MS, Rentsch D. 2012. Determinants for Arabidopsis peptide transporter targeting to the tonoplast or plasma membrane. Traffic 13, 1090–1105.

Krogh A, Larsson B, von Heijne G, Sonnhammer ELL. 2001. Predicting transmembrane protein topology with a hidden Markov model: application to complete genomes. Journal of Molecular Biology 305, 567–580.

Ladwig F, Stahl M, Ludewig U, Hirner AA, Hammes UZ, Stadler R, Harter K, Koch W. 2012. Siliques are Red1 from Arabidopsis acts as a bidirectional amino acid transporter that is crucial for the amino acid homeostasis of siliques. Plant Physiology 158, 1643–1655.

Lagarde D, Basset M, Lepetit M, Conejero G, Gaymard F, Astruc S, Grignon C. 1996. Tissue-specific expression of *Arabidopsis AKT1* gene is consistent with a role in K^+^ nutrition. The Plant Journal 9, 195–203.

Li E, Wimley WC, Hristova K. 2012. Transmembrane helix dimerization: beyond the search for sequence motifs. Biochimica et Biophysica Acta 1818, 183–193.

Liang Y, Richardson S, Yan J, Benites VT, Cheng-Yue C, Tran T, Mortimer J, Mukhopadhyay A, Keasling JD, Scheller HV, Loque D. 2017. Endoribonuclease-Based Two-Component Repressor Systems for Tight Gene Expression Control in Plants. ACS Synthetic Biology 6, 806–816.

Lohaus G, Pennewiss K, Sattelmacher B, Hussmann M, Hermann Muehling K. 2001. Is the infiltration-centrifugation technique appropriate for the isolation of apoplastic fluid? A critical evaluation with different plant species. Physiologia Plantarum 111, 457–465.

Lomize MA, Pogozheva ID, Joo H, Mosberg HI, Lomize AL. 2012. OPM database and PPM web server: resources for positioning of proteins in membranes. Nucleic Acids Research 40, D370–376.

Loque D, Lalonde S, Looger LL, von Wiren N, Frommer WB. 2007. A cytosolic trans-activation domain essential for ammonium uptake. Nature 446, 195–198.

McGuffin LJ, Adiyaman R, Maghrabi AHA, Shuid AN, Brackenridge DA, Nealon JO, Philomina LS. 2019. IntFOLD: an integrated web resource for high performance protein structure and function prediction. Nucleic Acids Research 47, W408–W413.

Müller B. 2016. Characterization of UmamiTs in Arabidopsis: amino acid transporters involved in amino acid cycling, phloem unloading and the supply of symplasmically isolated sink tissues, University of Regensburg Regensburg, Germany.

Müller B, Fastner A, Karmann J, Mansch V, Hoffmann T, Schwab W, Suter-Grotemeyer M, Rentsch D, Truernit E, Ladwig F, Bleckmann A, Dresselhaus T, Hammes UZ. 2015. Amino Acid Export in Developing Arabidopsis Seeds Depends on UmamiT Facilitators. Current Biology 25, 3126–3131.

Okumoto S, Pilot G. 2011. Amino acid export in plants: a missing link in nitrogen cycling. Molecular Plant 4, 453–463.

Omasits U, Ahrens CH, Muller S, Wollscheid B. 2014. Protter: interactive protein feature visualization and integration with experimental proteomic data. Bioinformatics 30, 884–886.

Pan Q, Cui B, Deng F, Quan J, Loake GJ, Shan W. 2016. RTP1 encodes a novel endoplasmic reticulum (ER)-localized protein in Arabidopsis and negatively regulates resistance against biotrophic pathogens. New Phytologist 209, 1641–1654.

Pedrazzini E, Komarova NY, Rentsch D, Vitale A. 2013. Traffic routes and signals for the tonoplast. Traffic 14, 622–628.

Pratelli R, Guerra DD, Yu S, Wogulis M, Kraft E, Frommer WB, Callis J, Pilot G. 2012. The ubiquitin E3 ligase LOSS OF GDU2 is required for GLUTAMINE DUMPER1-induced amino acid secretion in Arabidopsis. Plant Physiology 158, 1628–1642.

Pratelli R, Pilot G. 2014. Regulation of amino acid metabolic enzymes and transporters in plants. Journal of Experimental Botany 65, 5535–5556.

Pratelli R, Voll LM, Horst RJ, Frommer WB, Pilot G. 2010. Stimulation of nonselective amino acid export by glutamine dumper proteins. Plant Physiology 152, 762–773.

Ranocha P, Denance N, Vanholme R, Freydier A, Martinez Y, Hoffmann L, Kohler L, Pouzet C, Renou JP, Sundberg B, Boerjan W, Goffner D. 2010. Walls are thin 1 (WAT1), an Arabidopsis homolog of Medicago truncatula NODULIN21, is a tonoplast-localized protein required for secondary wall formation in fibers. The Plant Journal 63, 469–483.

Ranocha P, Dima O, Nagy R, Felten J, Corratge-Faillie C, Novak O, Morreel K, Lacombe B, Martinez Y, Pfrunder S, Jin X, Renou JP, Thibaud JB, Ljung K, Fischer U, Martinoia E, Boerjan W, Goffner D. 2013. Arabidopsis WAT1 is a vacuolar auxin transport facilitator required for auxin homoeostasis. Nature Communications 4, 2625.

Rentsch D, Schmidt S, Tegeder M. 2007. Transporters for uptake and allocation of organic nitrogen compounds in plants. FEBS letters 581, 2281–2289.

Riens B, Lohaus G, Heineke D, Heldt HW. 1991. Amino acid and sucrose content determined in the cytosolic, chloroplastic, and vacuolar compartments and in the phloem sap of spinach leaves. Plant Physiology 97, 227–233.

Schmid M, Davison TS, Henz SR, Pape UJ, Demar M, Vingron M, Scholkopf B, Weigel D, Lohmann JU. 2005. A gene expression map of Arabidopsis thaliana development. Nature Genetics 37, 501–506.

Shin K, Lee S, Song WY, Lee RA, Lee I, Ha K, Koo JC, Park SK, Nam HG, Lee Y, Soh MS. 2015. Genetic identification of ACC-RESISTANT2 reveals involvement of LYSINE HISTIDINE TRANSPORTER1 in the uptake of 1-aminocyclopropane-1-carboxylic acid in Arabidopsis thaliana. Plant and Cell Physiology 56, 572–582.

Song Y, DiMaio F, Wang RY, Kim D, Miles C, Brunette T, Thompson J, Baker D. 2013. High-resolution comparative modeling with RosettaCM. Structure 21, 1735–1742.

Stamatakis A. 2014. RAxML version 8: a tool for phylogenetic analysis and post-analysis of large phylogenies. Bioinformatics 30, 1312–1313.

Studer G, Biasini M, Schwede T. 2014. Assessing the local structural quality of transmembrane protein models using statistical potentials (QMEANBrane). Bioinformatics 30, i505–511.

Tamura K, Peterson D, Peterson N, Stecher G, Nei M, Kumar S. 2011. MEGA5: molecular evolutionary genetics analysis using maximum likelihood, evolutionary distance, and maximum parsimony methods. Molecular Biology & Evolution 28, 2731–2739.

Tang Y, Zhang Z, Lei Y, Hu G, Liu J, Hao M, Chen A, Peng Q, Wu J. 2019. Cotton WATs Modulate SA Biosynthesis and Local Lignin Deposition Participating in Plant Resistance Against Verticillium dahliae. Frontiers in Plant Science 10, 526.

Tegeder M, Rentsch D. 2010. Uptake and partitioning of amino acids and peptides. Molecular Plant 3, 997–1011.

Toufighi K, Brady SM, Austin R, Ly E, Provart NJ. 2005. The Botany Array Resource: e-Northerns, Expression Angling, and promoter analyses. The Plant Journal 43, 153–163.

Tsirigos KD, Peters C, Shu N, Kall L, Elofsson A. 2015. The TOPCONS web server for consensus prediction of membrane protein topology and signal peptides. Nucleic Acids Research 43, W401–407.

Vastermark A, Wollwage S, Houle ME, Rio R, Saier MHJr., 2014. Expansion of the APC superfamily of secondary carriers. Proteins 82, 2797–2811.

Velasco I, Tenreiro S, Calderon IL, André B. 2004. *Saccharomyces cerevisiae* Aqr1 is an internal-membrane transporter involved in excretion of amino acids. Eukaryotic Cell 3, 1492–1503.

Walters RF, DeGrado WF. 2006. Helix-packing motifs in membrane proteins. Proceedings of the National Academy of Science Of USA 103, 13658–13663.

Waterhouse A, Bertoni M, Bienert S, Studer G, Tauriello G, Gumienny R, Heer FT, de Beer TAP, Rempfer C, Bordoli L, Lepore R, Schwede T. 2018. SWISS-MODEL: homology modelling of protein structures and complexes. Nucleic Acids Research 46, W296–W303.

Weiner H, Heldt HW. 1992. Inter- and intracellular distribution of amino acids and other metabolites in maize (*Zea mays* L.) leaves. Planta 187, 242–246.

Winter H, Robinson DG, Heldt HW. 1993. Subcellular volumes and metabolite concentrations in barley leaves. Planta 191, 180–190.

Winter H, Robinson DG, Heldt HW. 1994. Subcellular volumes and metabolite concentrations in spinach leaves. Planta 193, 530–535.

Wolfenstetter S, Wirsching P, Dotzauer D, Schneider S, Sauer N. 2012. Routes to the tonoplast: the sorting of tonoplast transporters in Arabidopsis mesophyll protoplasts. The Plant Cell 24, 215–232.

Xu J, Zhang Y. 2010. How significant is a protein structure similarity with TM-score = 0.5? Bioinformatics 26, 889–895.

Yang Y, Hammes UZ, Taylor CG, Schachtman DP, Nielsen E. 2006. High-affinity auxin transport by the AUX1 influx carrier protein. Current Biology 16, 1123–1127.

Zhao H, Ma H, Yu L, Wang X, Zhao J. 2012. Genome-wide survey and expression analysis of amino acid transporter gene family in rice (Oryza sativa L.). PLoS One 7, e49210.

