## Supplemental Figures for "Detailed characterization of the UMAMITs provides insight into their evolution, functional properties as amino acid transporters and role in the plant"

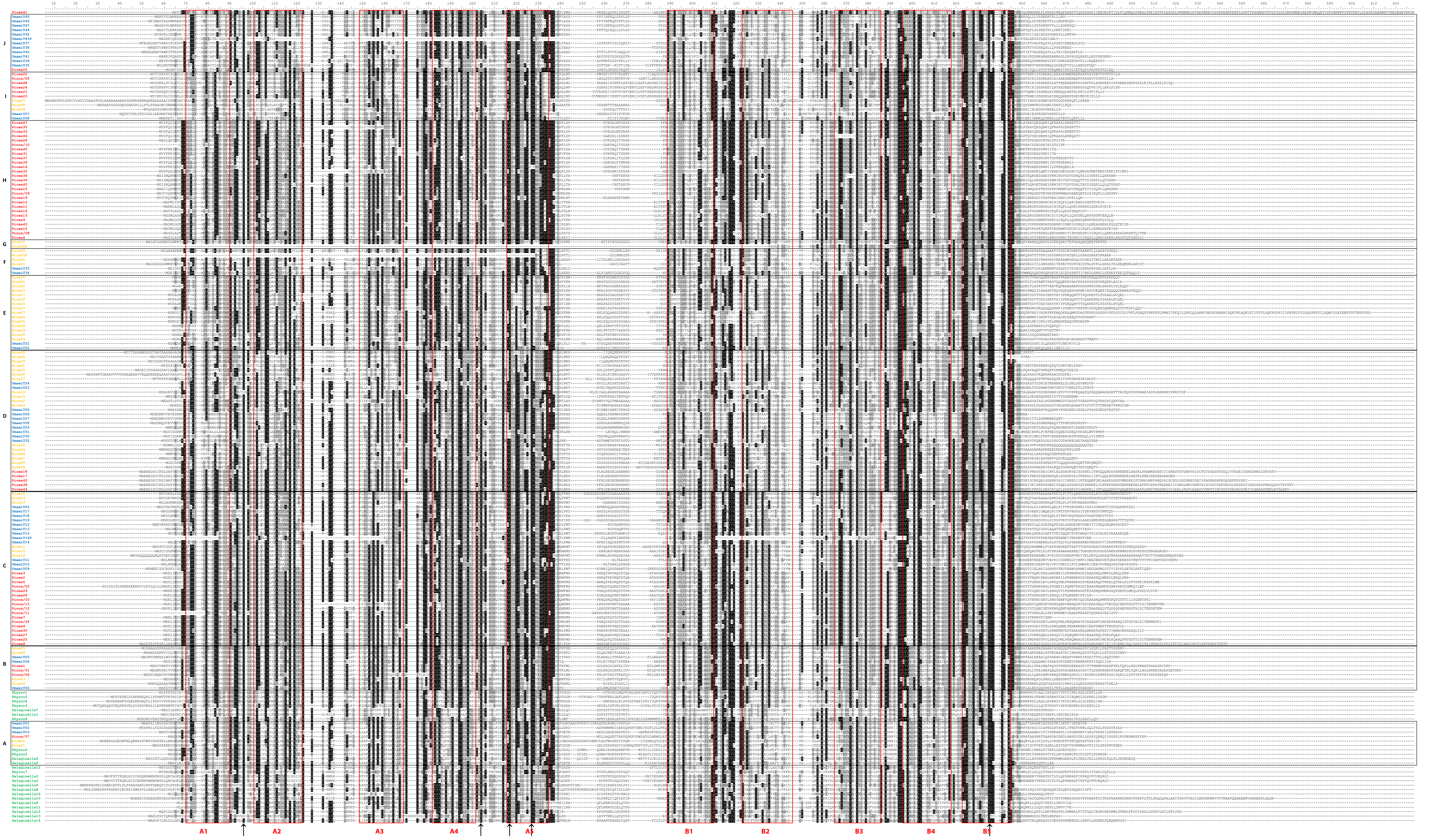


**Supplemental Figure 1: Multiple sequence alignment of 180 UMAMIT sequences.**
Proteins from Arabidopsis, rice, pine, *Selaginella* and *Physcomitrella* were aligned using ClustalW. Residues mutagenized in UMAMIT14 are localized on the alignment at positions 96 (G33), 204 (E122), 212 (K134), 227 (G144) and 435 (G315). Letters A to J on the left correspond to the clades from Figure 1.

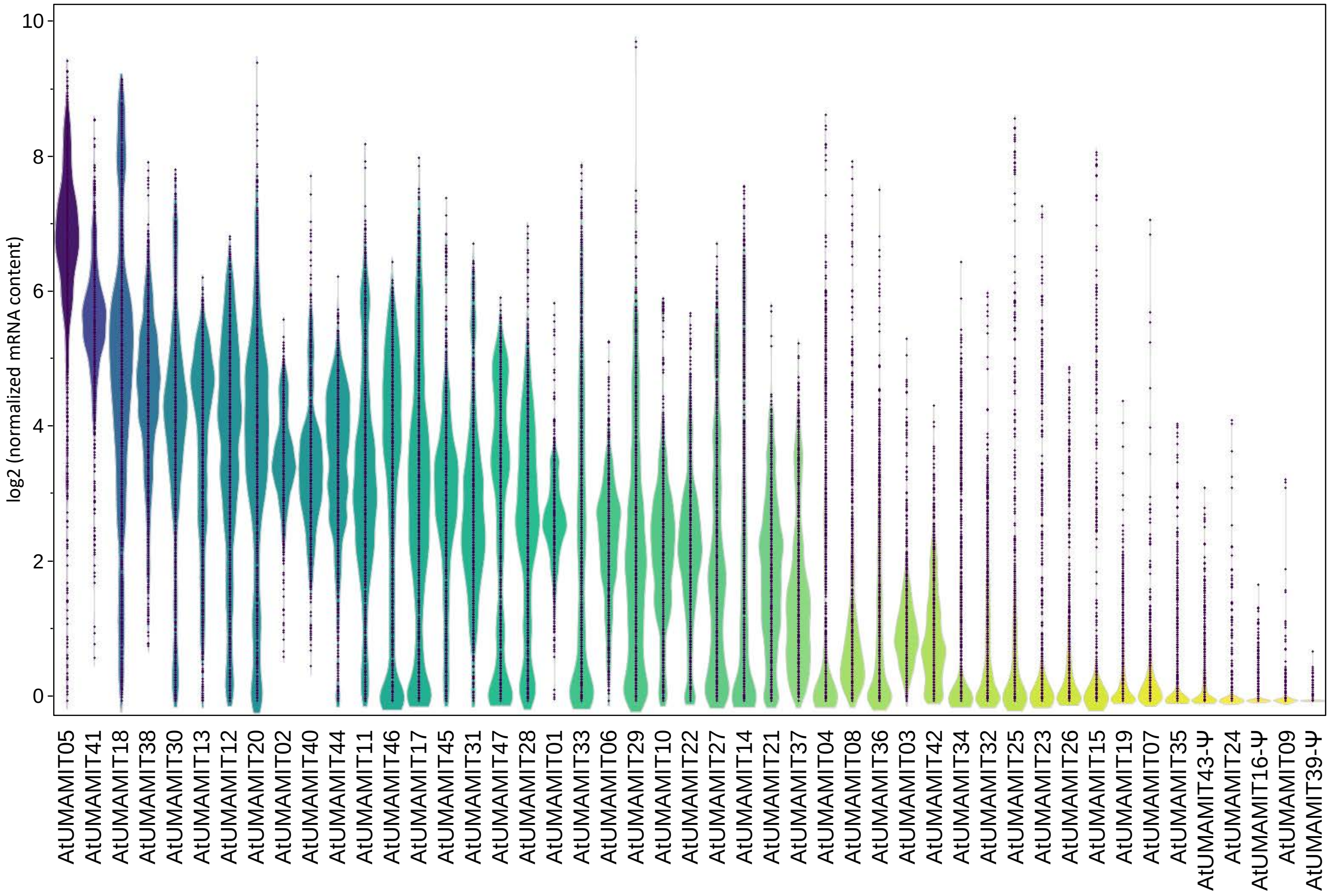


**Supplemental Figure 2: Distribution of the mRNA contents of the AtUMAMIT genes in Arabidopsis in various conditions and organs.**

Violin plot of the mRNA content determined by RNAseq from 1229 publicly available samples, extracted from Genevestigator-normalized data in August 2020 (details about normalization can be obtained from Genevestigator support). Plots are colored based on the average expression of the genes.

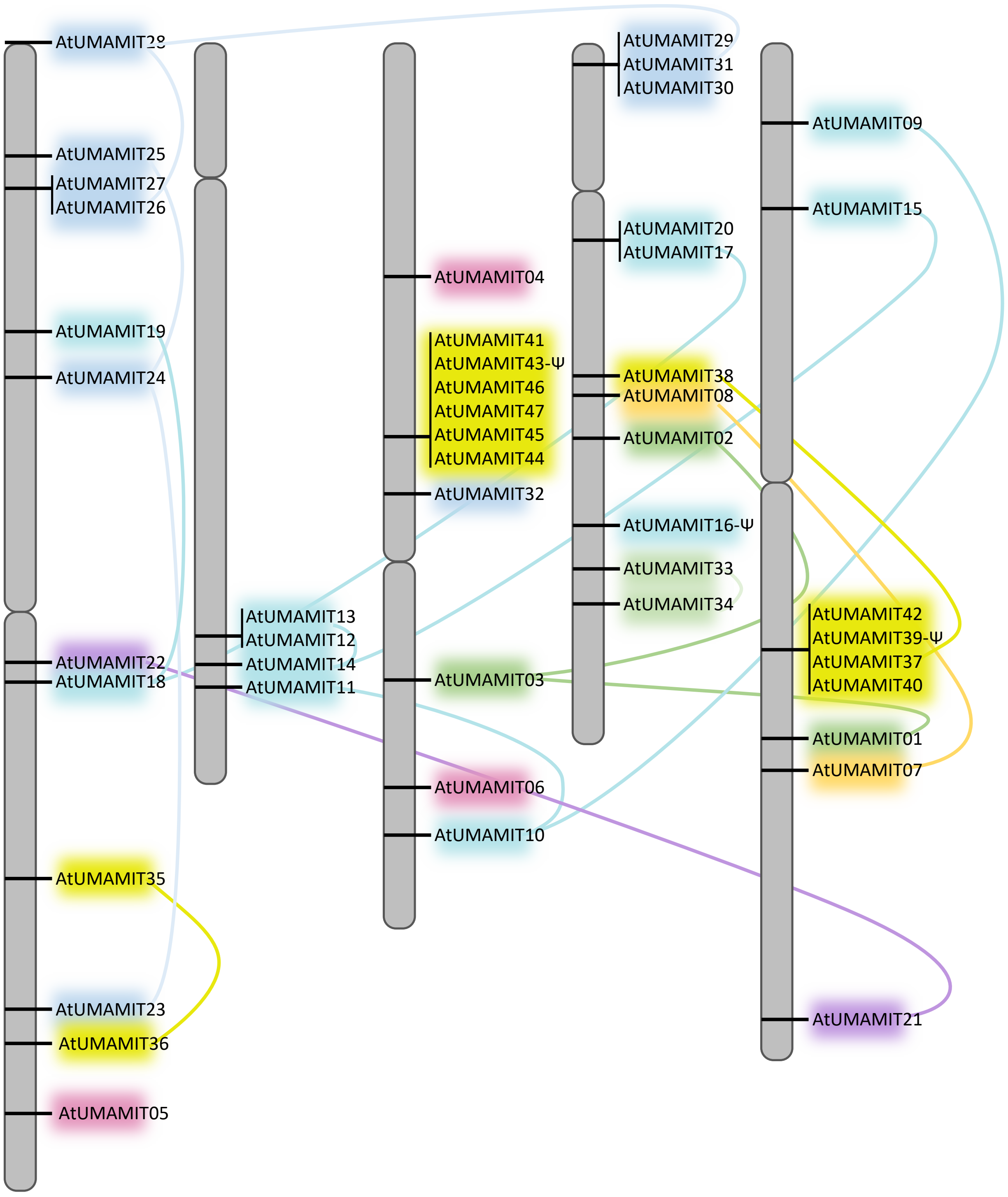


**Supplemental Figure 3: Position of the AtUMAMIT genes on the Arabidopsis chromosomes.**
The position of the genes was determined using the Chromosome Map Tool from TAIR (https://www.arabidopsis.org/). Color schemes correspond to the clades from Figure 1, and the curved lines highlight the closely related genes.

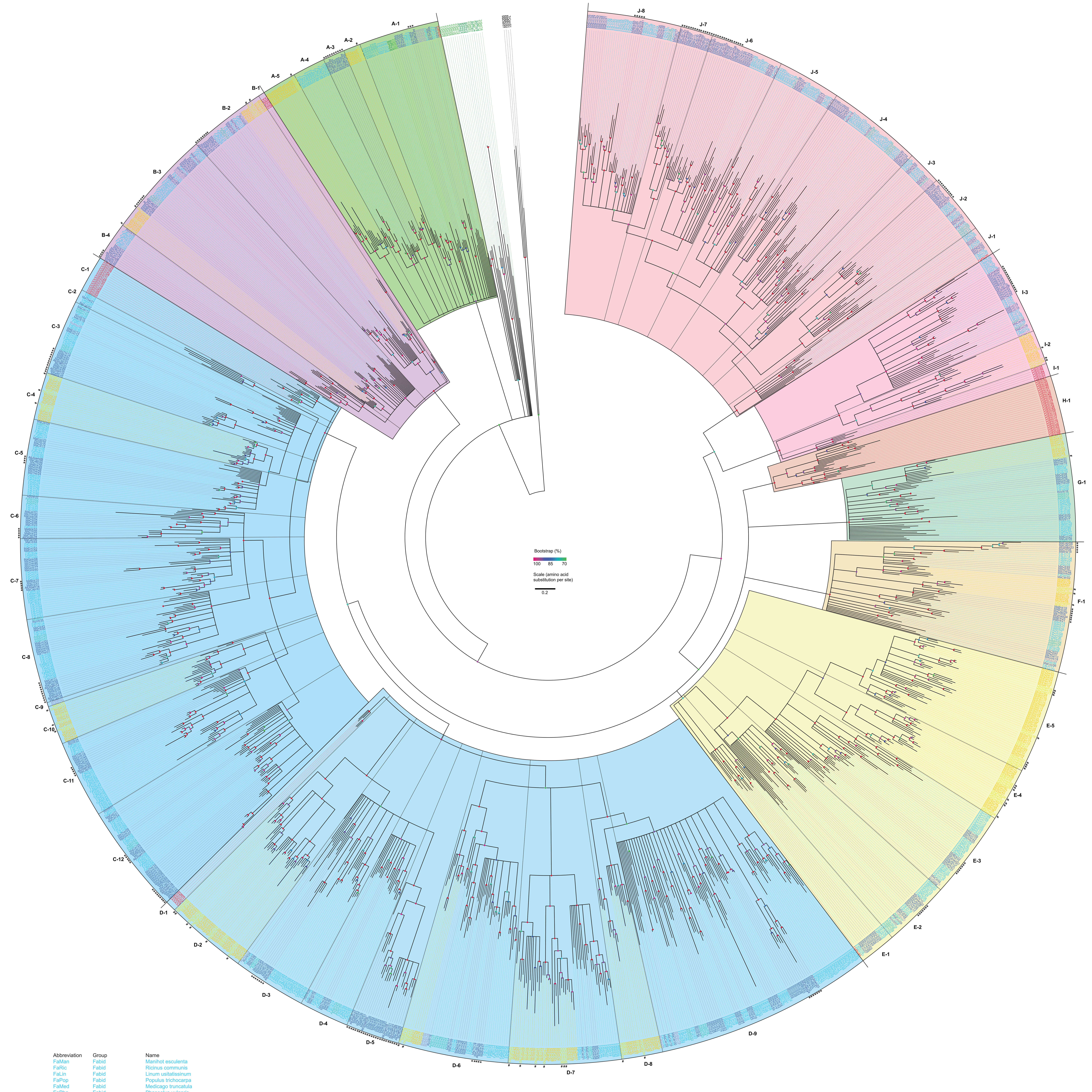


| Abbreviation | Group | Name | |
|---|---|---|---|
| FaMan | Fabid | Manihot esculenta | |
| FaRic | Fabid | Ricinus communis | |
| FaLin | Fabid | Linum usitatissinum | |
| FaPop | Fabid | Populus trichocarpa | |
| FaMed | Fabid | Medicago truncatula | |
| FaPha | Fabid | Phaseolus vulgaris | |
| FaGly | Fabid | Glycine max | |
| FaCuc | Fabid | Cucumis sativus | |
| FaPru | Fabid | Prunus persica | |
| FaMal | Fabid | Malus domestica | |
| FaFra | Fabid | Fragaria vesca | |
| AtUmam | Malvid | Arabidopsis thaliana | * |
| MaCap | Malvid | Capsella rubella | * |
| MaBra | Malvid | Brassica rapa | * |
| MaThl | Malvid | Thellungiella halophyta | * |
| MaCar | Malvid | Carica papaya | |
| MaGos | Malvid | Gossypium raimondii | |
| MaThe | Malvid | Theobroma cacao | |
| MaCit | Malvid | Citrus sinensis | |
| MaCit | Malvid | Citrus clementina | |
| MaEuc | Malvid | Eucalyptus grandis | |
| RoVit | Rosid | Vitis vinifera | |
| AsSol | Asterid | Solanum lycopersicum | |
| AsMim | Asterid | Mimulus guttatus | |
| RuAqu | Ranunculales | Aquilegia coerulea | |
| MoSor | Monocots | Sorghum bicolor | |
| MoZea | Monocots | Zea mays | |
| MoSet | Monocots | Setaria italica | |
| MoPan | Monocots | Panicum virgatum | |
| OsUmam | Monocots | Oryza sativa | # |
| MoBra | Monocots | Brachypodium distachyon | |
| CoPic | Conifers | Picea abies | |
| CoPin | Conifers | Pinus pinaster | |
| TrSel | Tracheophyta | VSelaginella moellendorfii | |
| BrPh | Bryophyta | Physcomitrella patens | |
| MaMic | Mamiellales | Micromonas pusilla | |
| MaMic | Mamiellales | Micromonas pusilla | |
| MaOst | Mamiellales | Ostreococcus lucimarinus | |
| TrCoc | Trebouxiophyceae | Coccomyxa subellipsoida | |

**Supplemental Figure 4. Phylogenic tree of 1466 UMAMIT proteins**.
Proteins names are colored by species, see left pannel; # indicate rice sequences, * indicates Arabidopsis sequences. Colored dots on the branches indicate bootstrap values of 1000 replicates. The tree was constructed using RaxML from sequences of Supplemental File 1. Letters and numbers around the radial tree indicate Clades and Subclades.

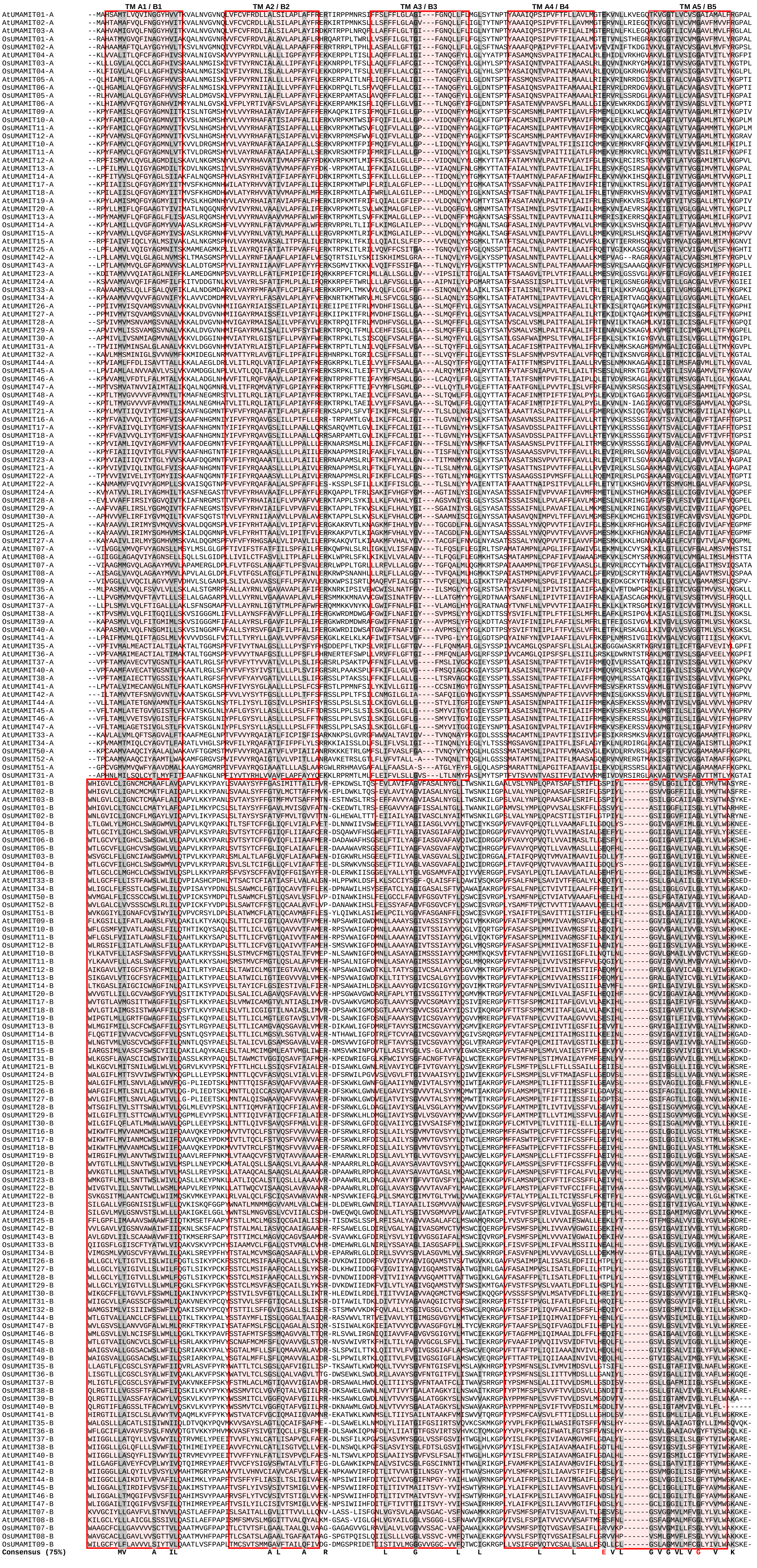


**Supplemental Figure 5: Alignment of the structural repeats of the OsUMAMIT and AtUMAMIT protein sequences.** The sequence of each protein was split between transmembrane domain A5 and B1, and the half-sequences (A1 to A5: A; B1 to B5: B) from all proteins were aligned. Un-aligned regions before A1, between A5 and B1, and after B5 were omitted from the alignment. Residues present in more than 75% of the half-sequences are indicated below the alignment (Consensus). Red residues are the residues mutagenized in AtUMAMIT14. Shading corresponds to 50% conservation (light gray) or 75% conservation (dark gray).

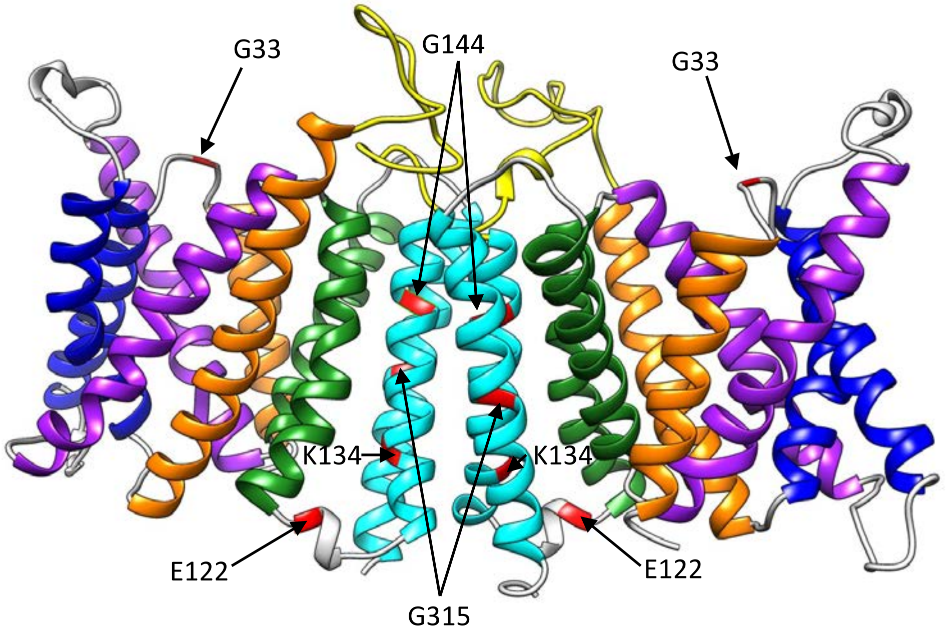


**Supplemental Figure 6: Model of the AtUMAMIT14 predicted protein dimer.**
The model was created by Swissmodel. Colors correspond to the homologous helices in the protein (*e.g.* A1 and B1 in orange) to show the arrangement of the transmembrane domains of the two structural repeats. Loops are colored in gray, except the extra-cytoplasmic loop located between structural repeats A and B. Red: mutagenized residues.

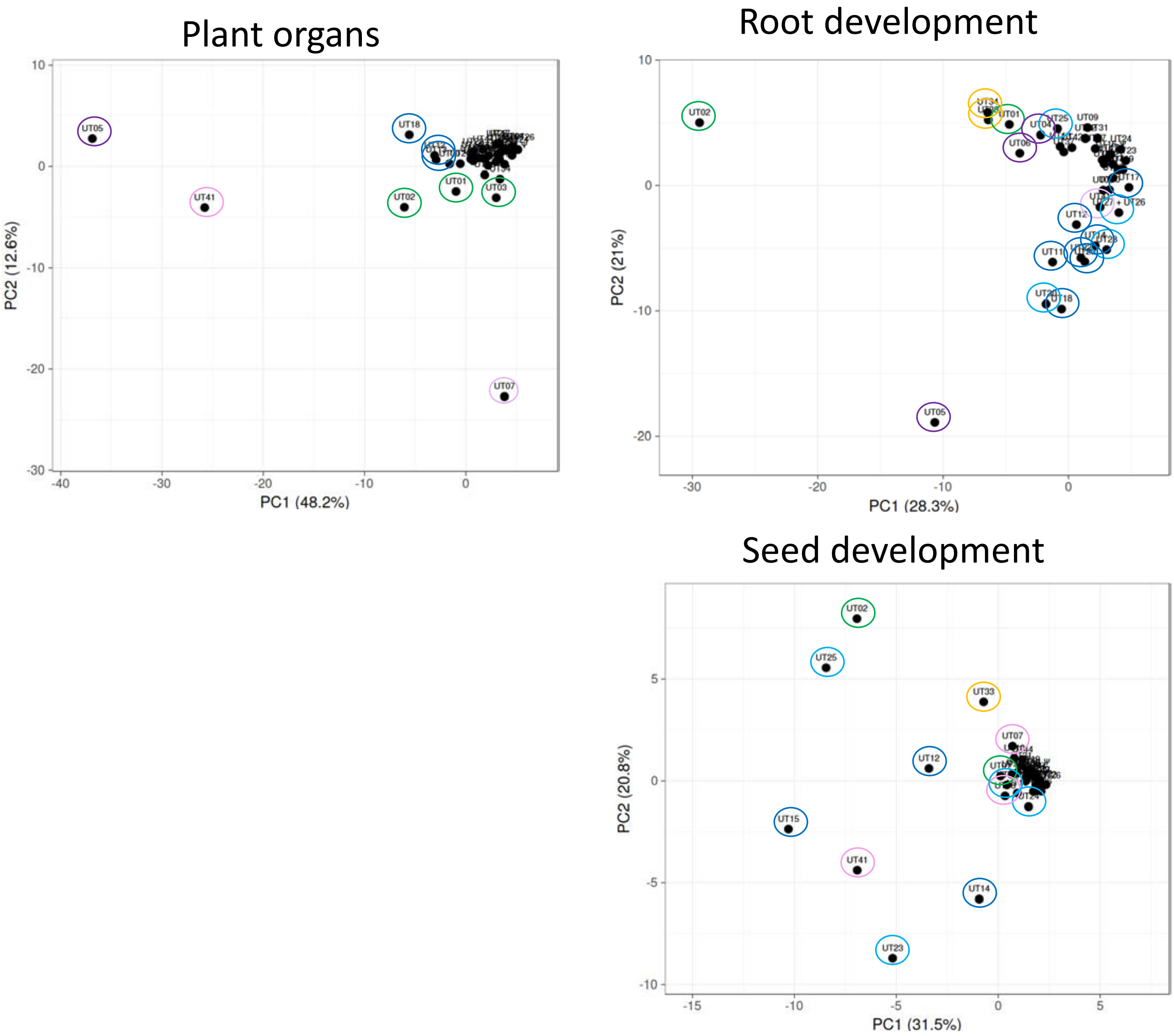


**Supplemental Figure 7: PCA analysis of the localization of the expression of the AtUMAMIT in the plant.**
Data from Supplemental File 3A, B and C were analyzed using JMP. The color of the circles around the best individualized genes corresponds to the Clades from Figure 1.

UMAIMT02

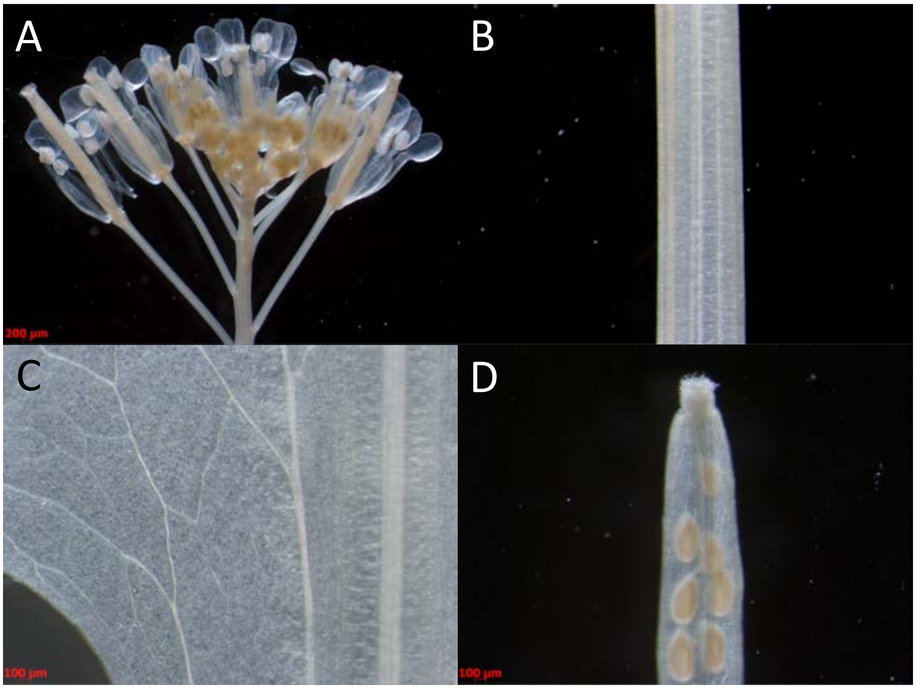


UMAIMT03

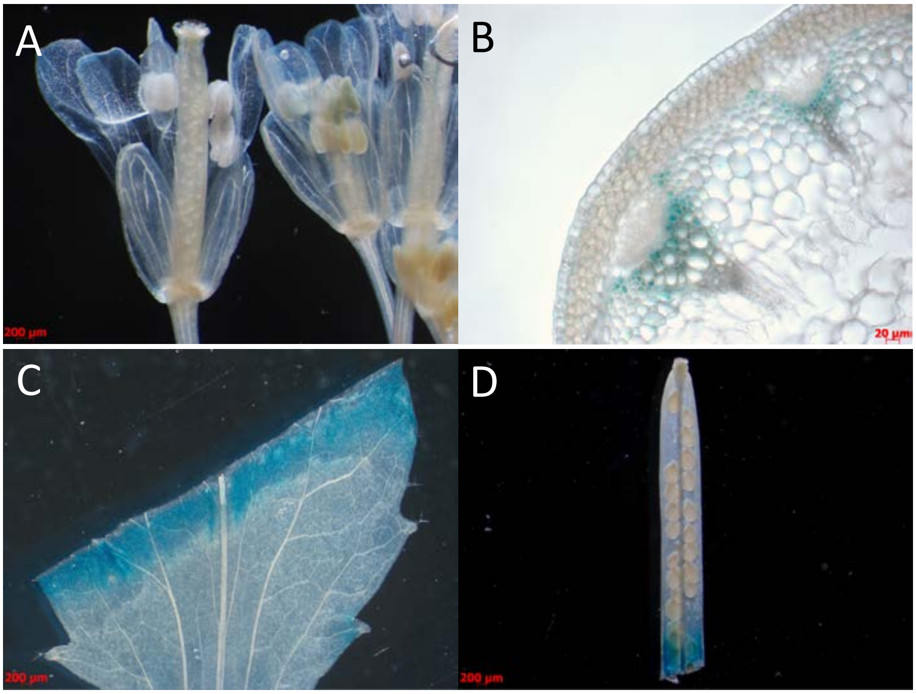

UMAIMT04

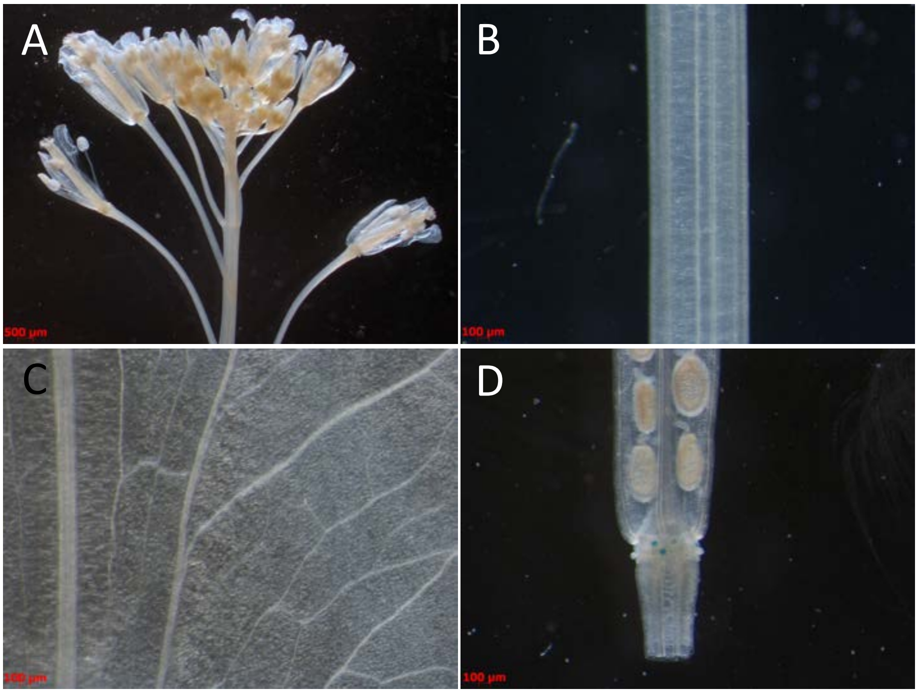


UMAIMT11

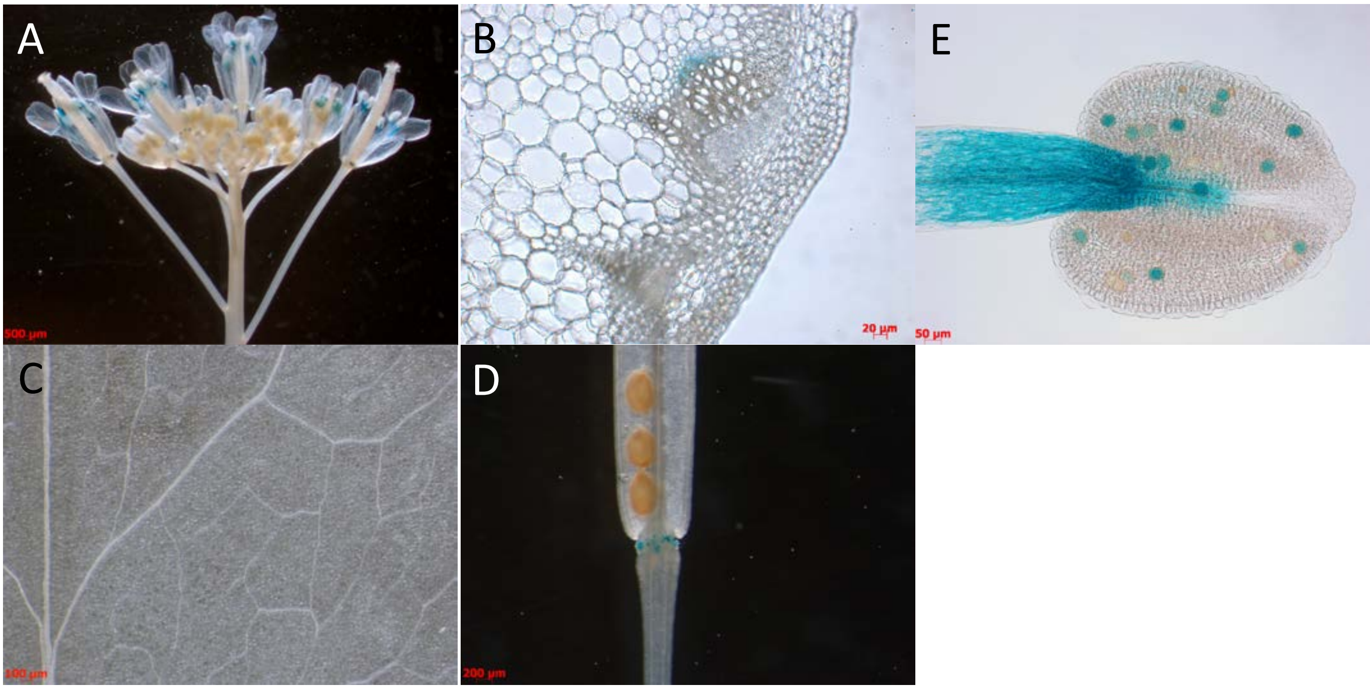

UMAMIT12

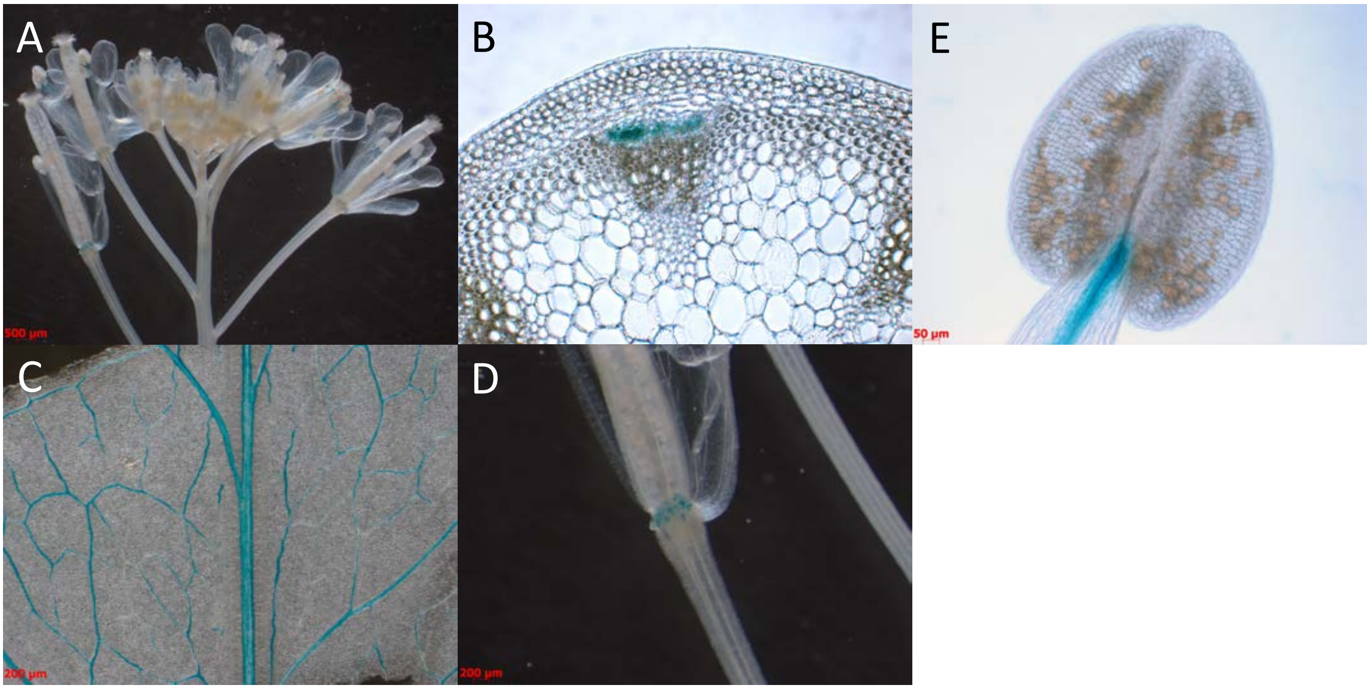


UMAMIT22

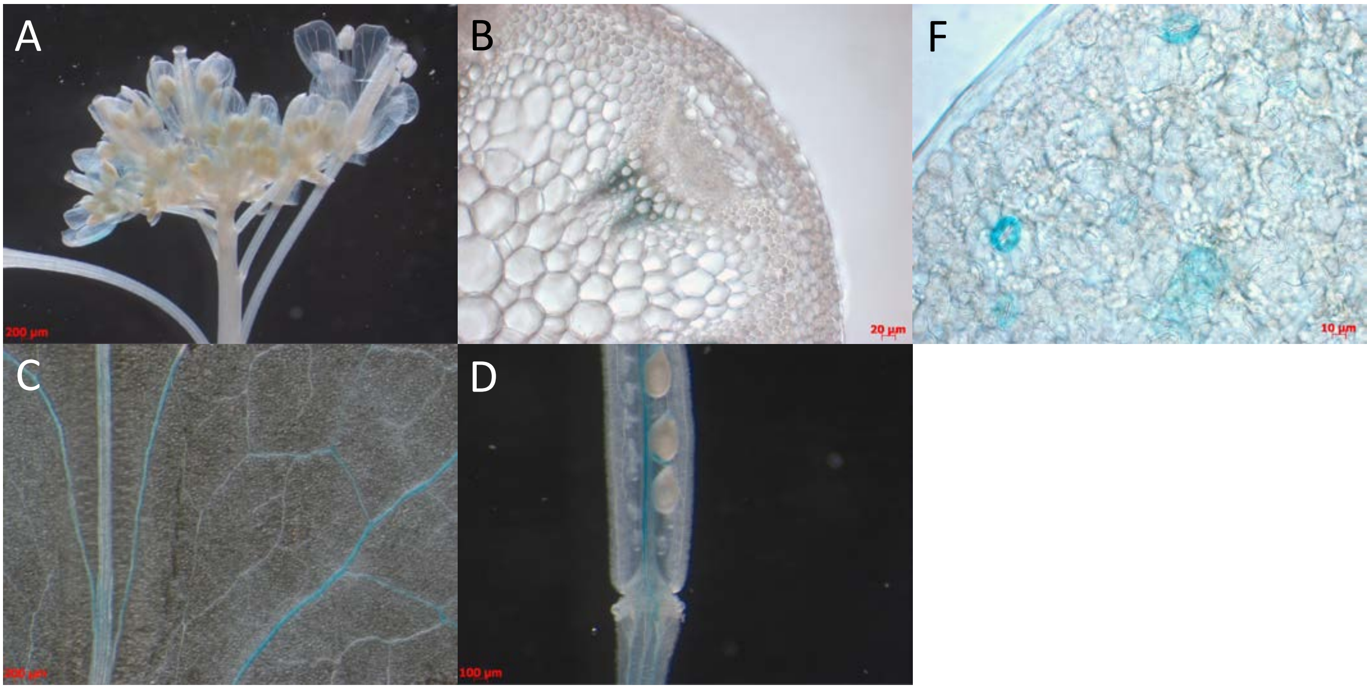

UMAMIT23

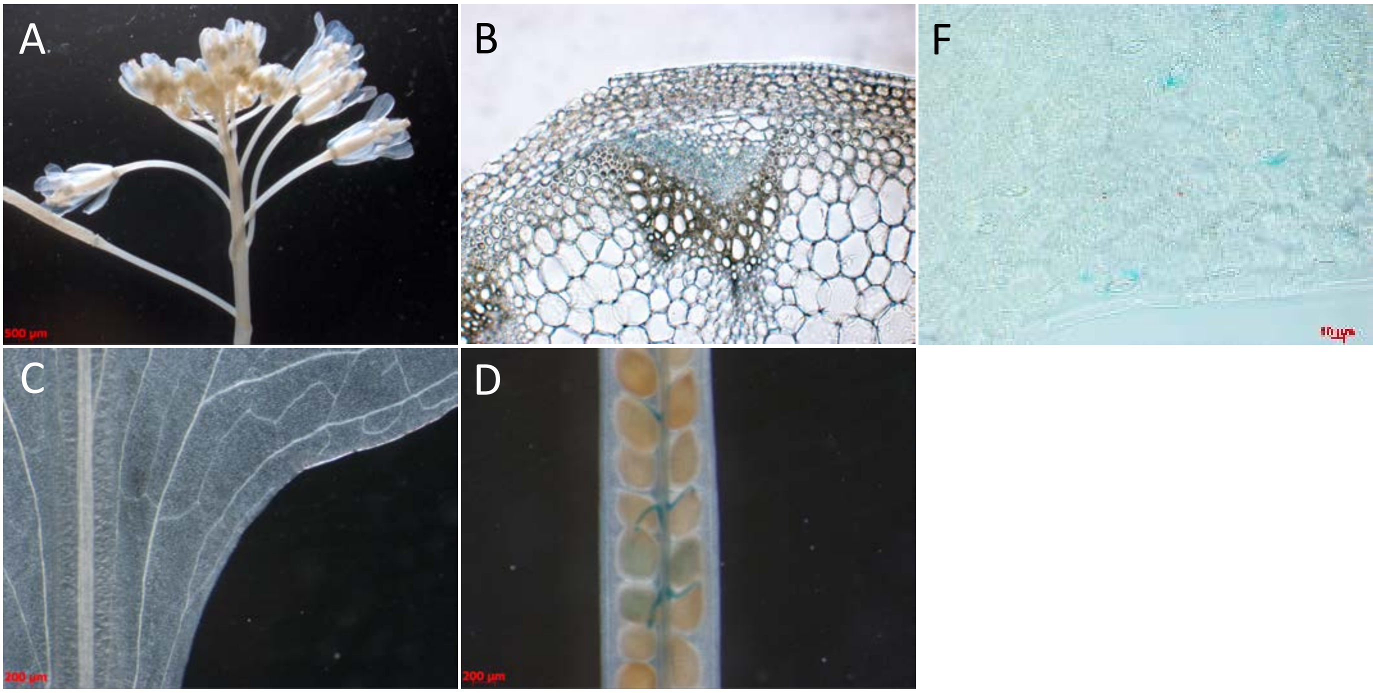


UMAMIT24

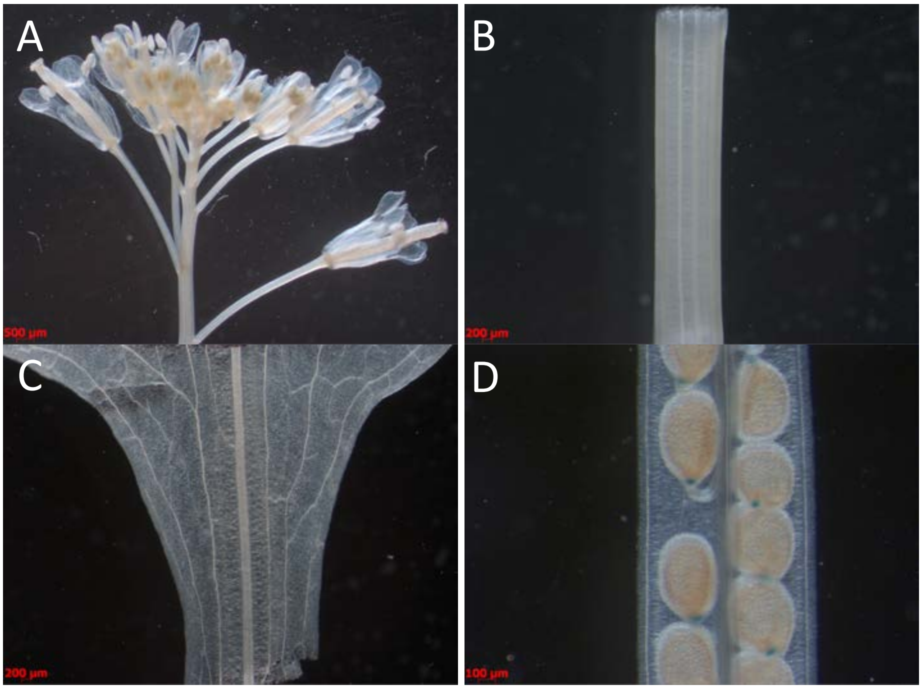

UMAMIT28

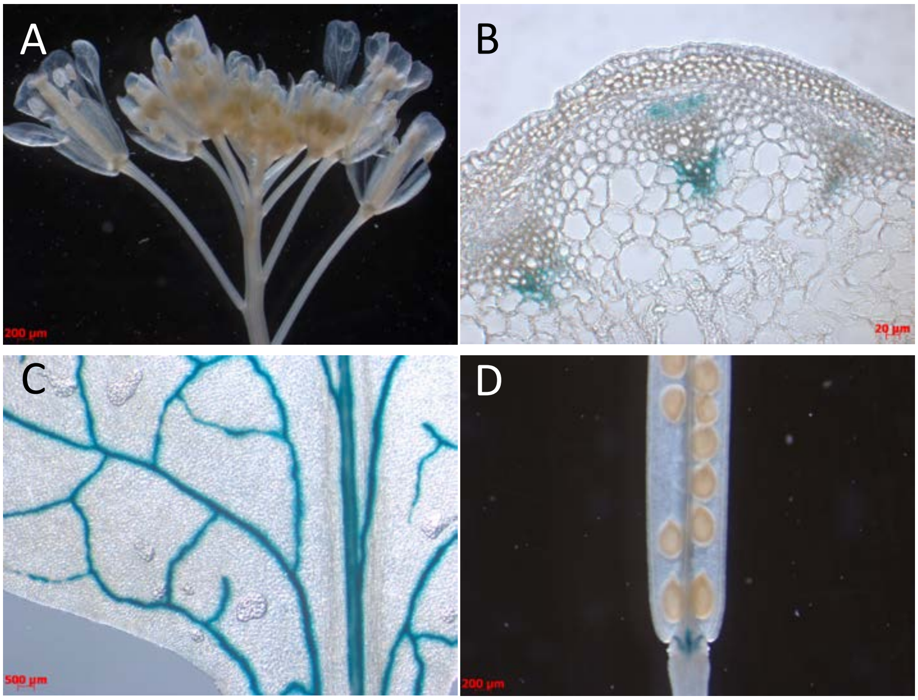


UMAMIT29

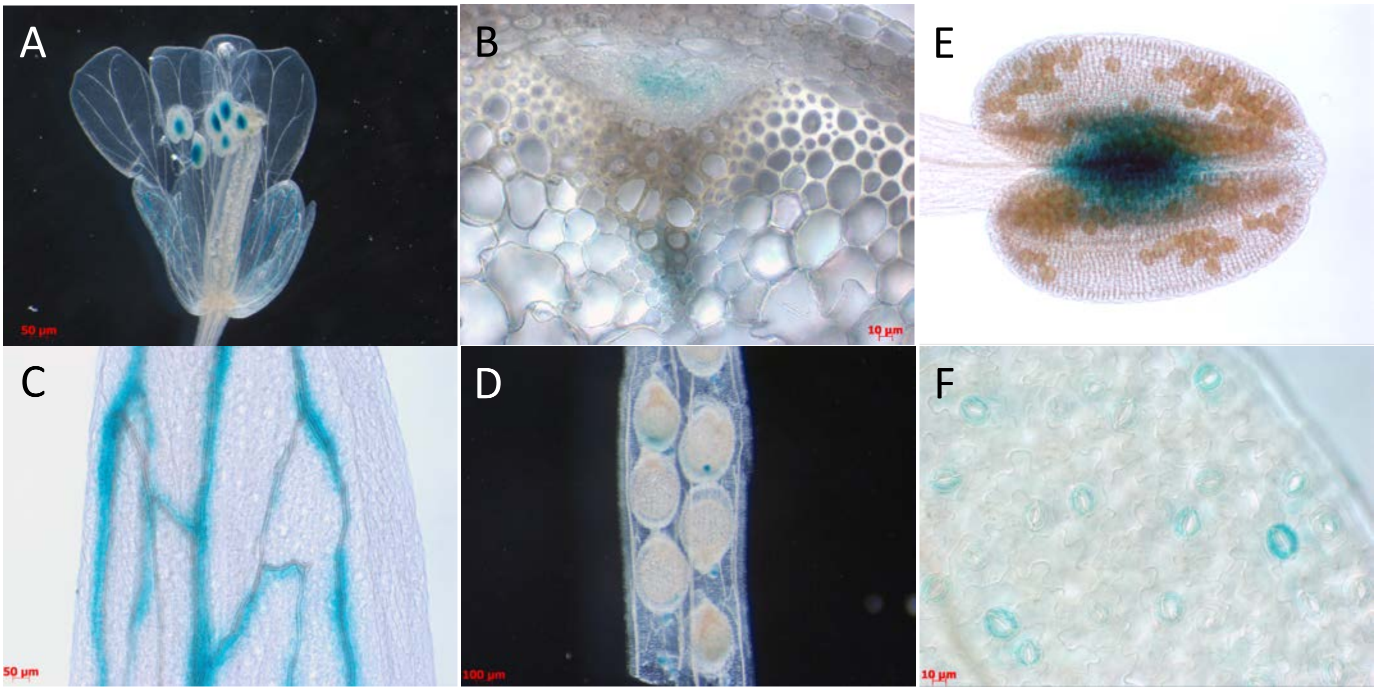

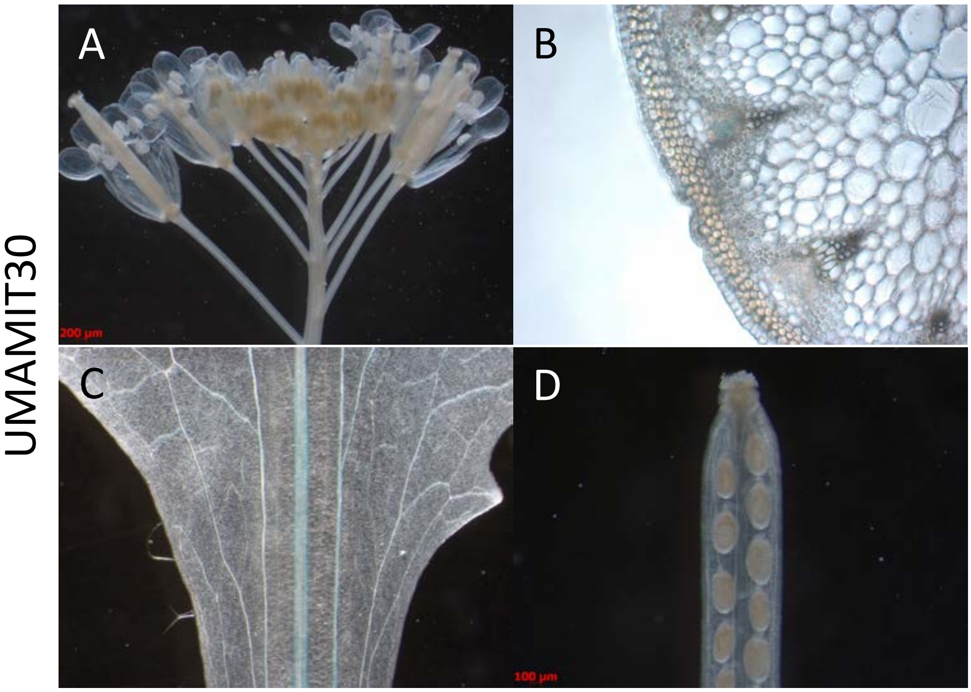
UMAMIT30
A
B
C
D
200 µm
100 µm

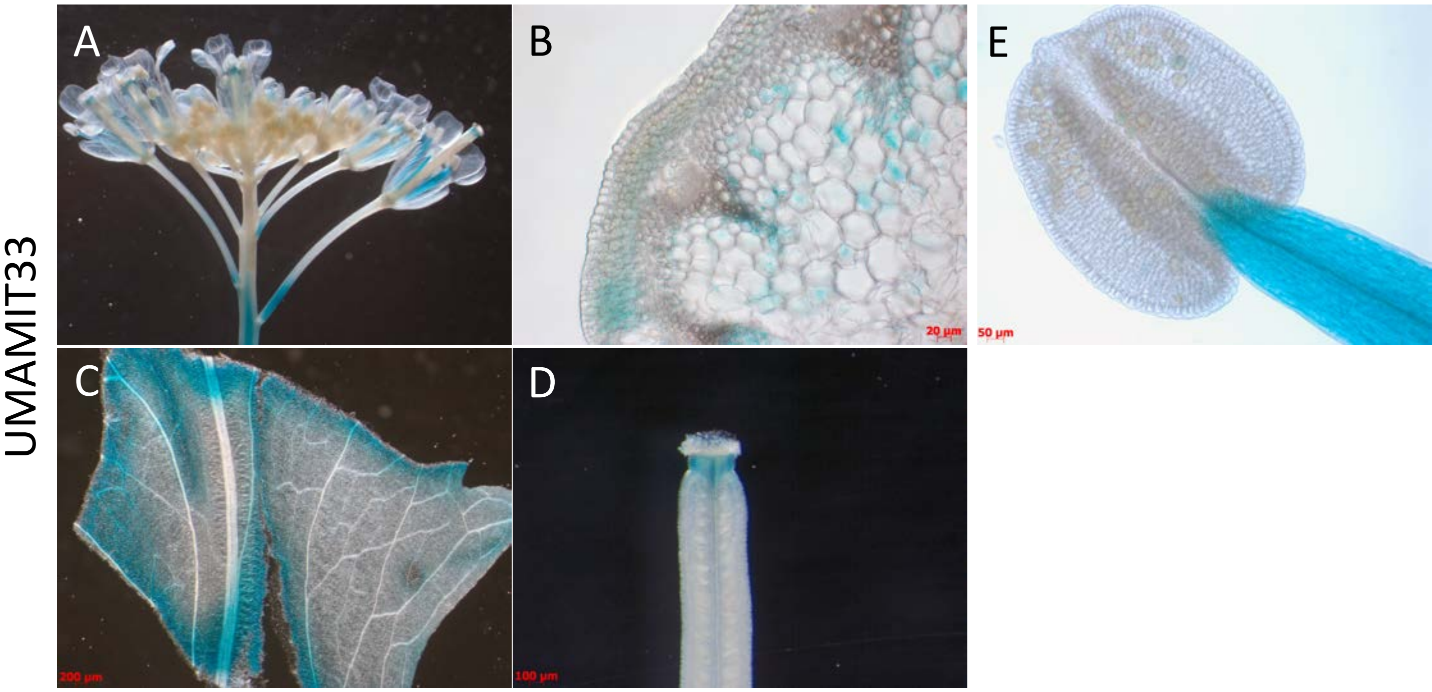
UMAMIT33
A
B
E
C
D
20 µm
50 µm
200 µm
100 µm

UMAMIT34

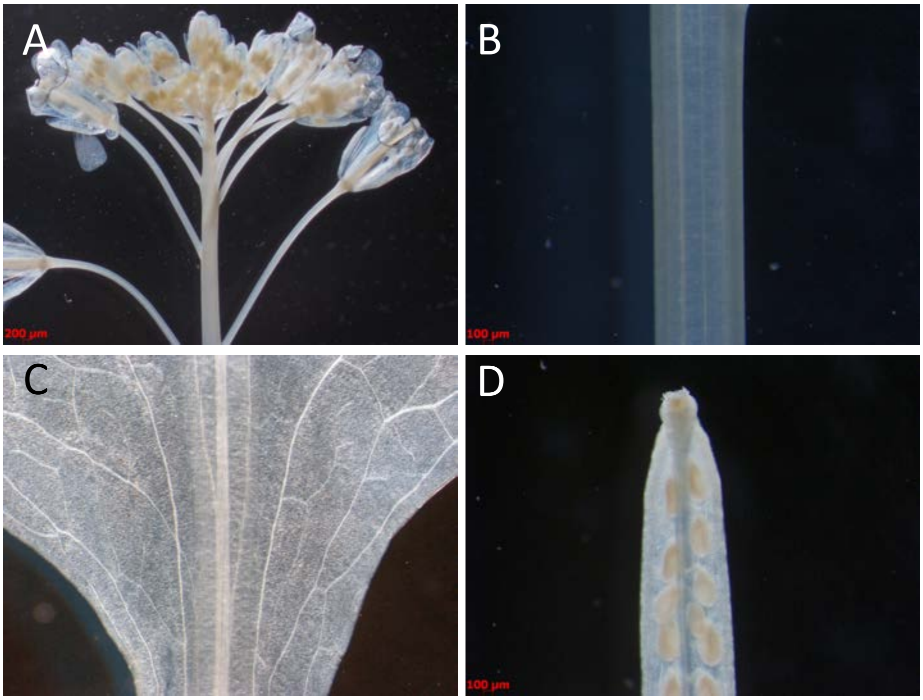


UMAMIT42

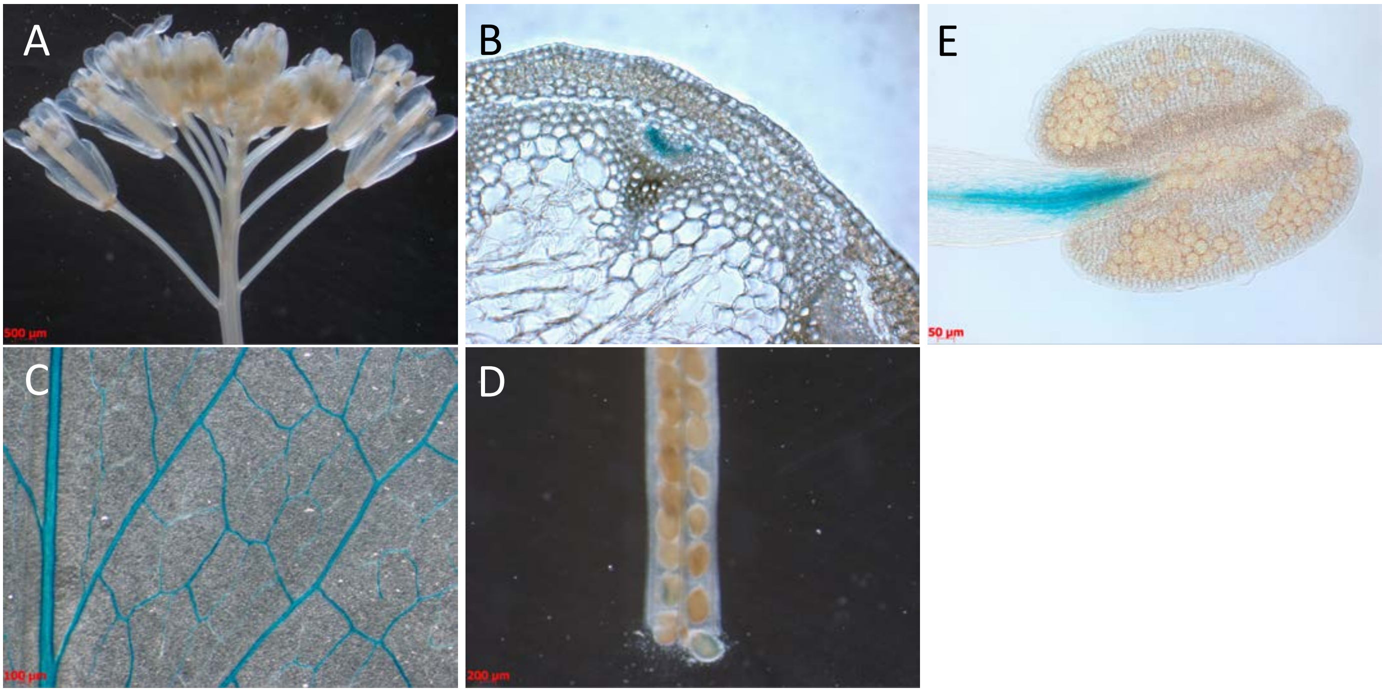

UMAMIT46

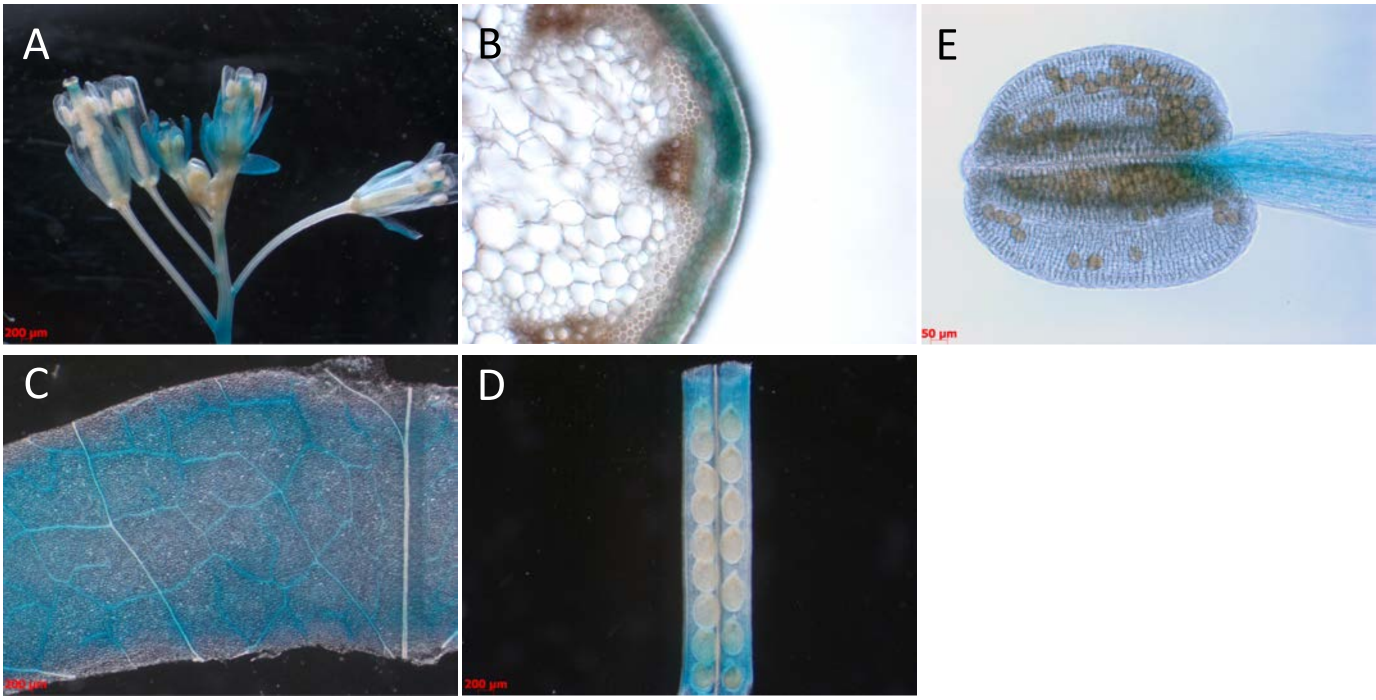


**Supplemental Figure 8: Expression patterns of 15 UMAMIT promoters.**
Plants expressing UMAMITp-GUS plants were grown on soil for 5-6 weeks, and the organs were stained independently.

A. Inflorescence / flower
B. Stem or stem cross-section
C. Leave detail
D. Silique
E. Stamen
F. Leaf epidermis

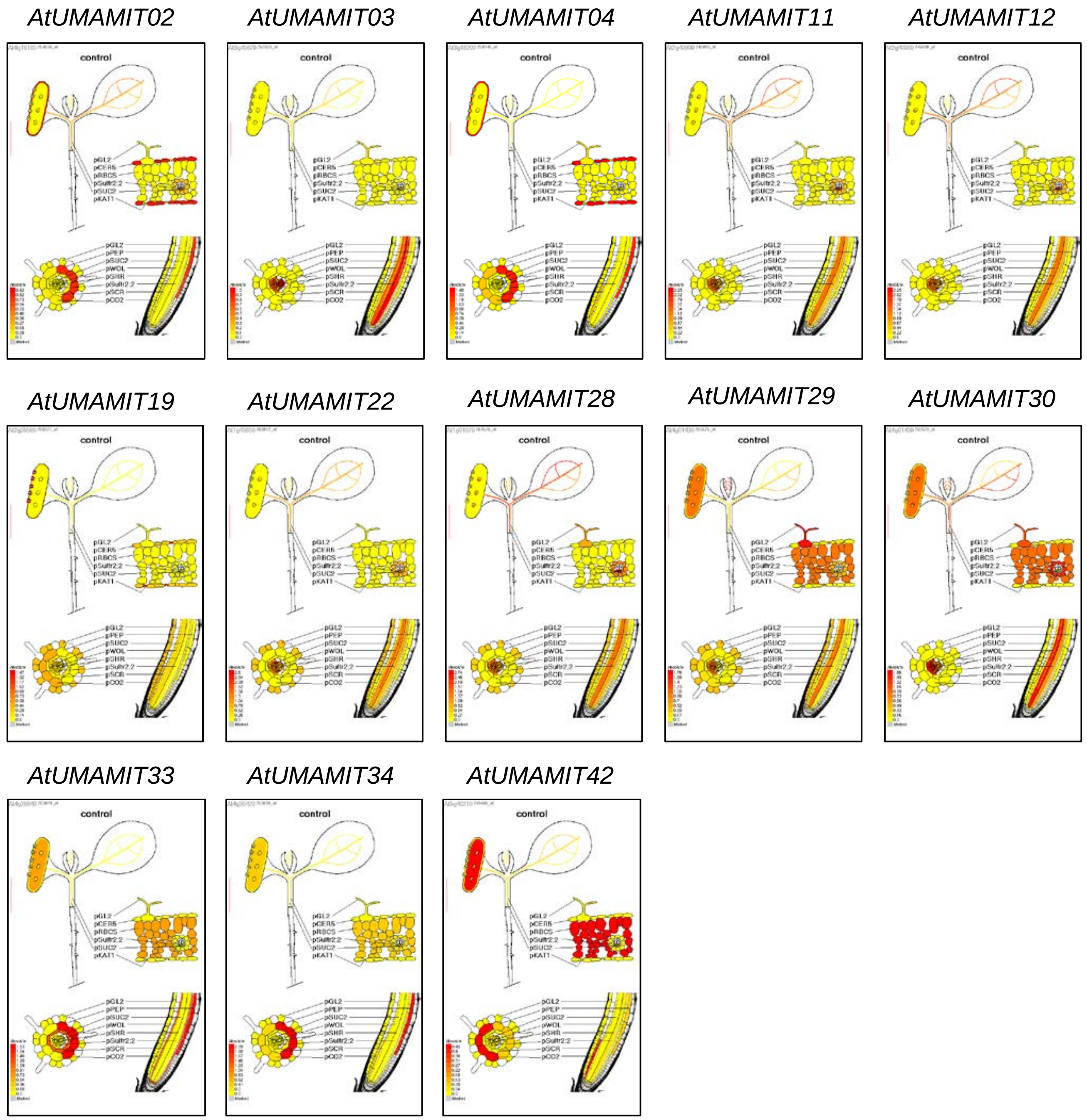


**Supplemental Figure 9: Localization in the plant of 13 *AtUMAMIT* genes, obtained from translatome data.** Data were obtained from http://efp.ucr.edu/cgi-bin/absolute.cgi. Briefly, mRNA localization was obtained by translatome analysis of transgenic plants that express a tagged ribosomal protein in specific cell types (Mustroph et al., 2009). This approach does not use making protoplasts from cells, contrary to the root expression dataset used in Supplemental File 3B. Data for *AtUMAMIT23* and *24* could not be retrieved from the server.

**Mustroph A, Zanetti ME, Jang CJ, Holtan HE, Repetti PP, Galbraith DW, Girke T, Bailey-Serres J**. 2009. Profiling translatomes of discrete cell populations resolves altered cellular priorities during hypoxia in Arabidopsis. *Proceedings of the National Academy of Science Of USA* **106**, 18843-18848.

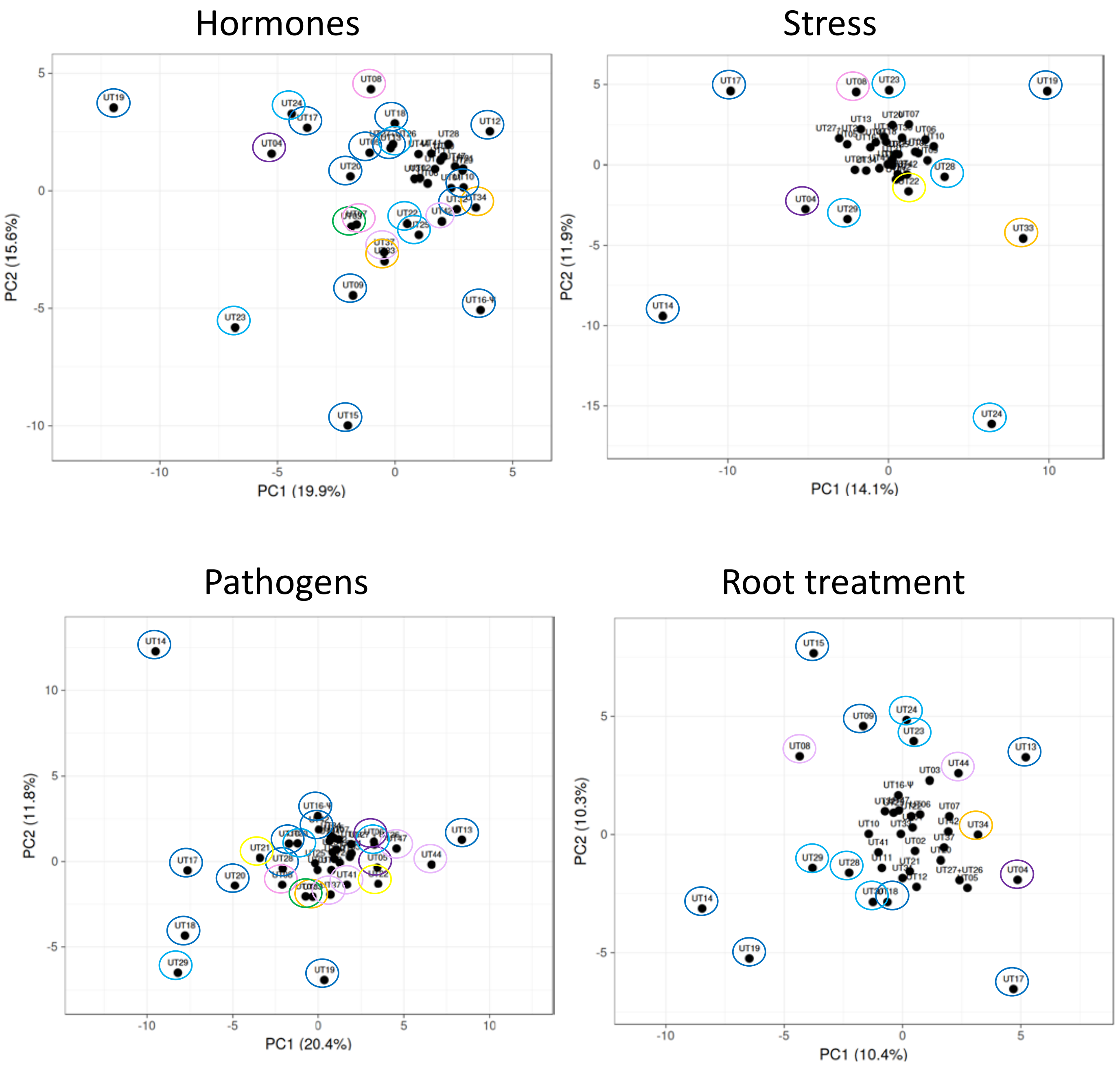


**Supplemental Figure 10: PCA analysis of the change in expression of the AtUMAMIT in the plant.**
Data from Supplemental File 3D, E, F and G were analyzed using JMP. The color of the circles around the best individualized genes corresponds to the Clades from Figure 1.

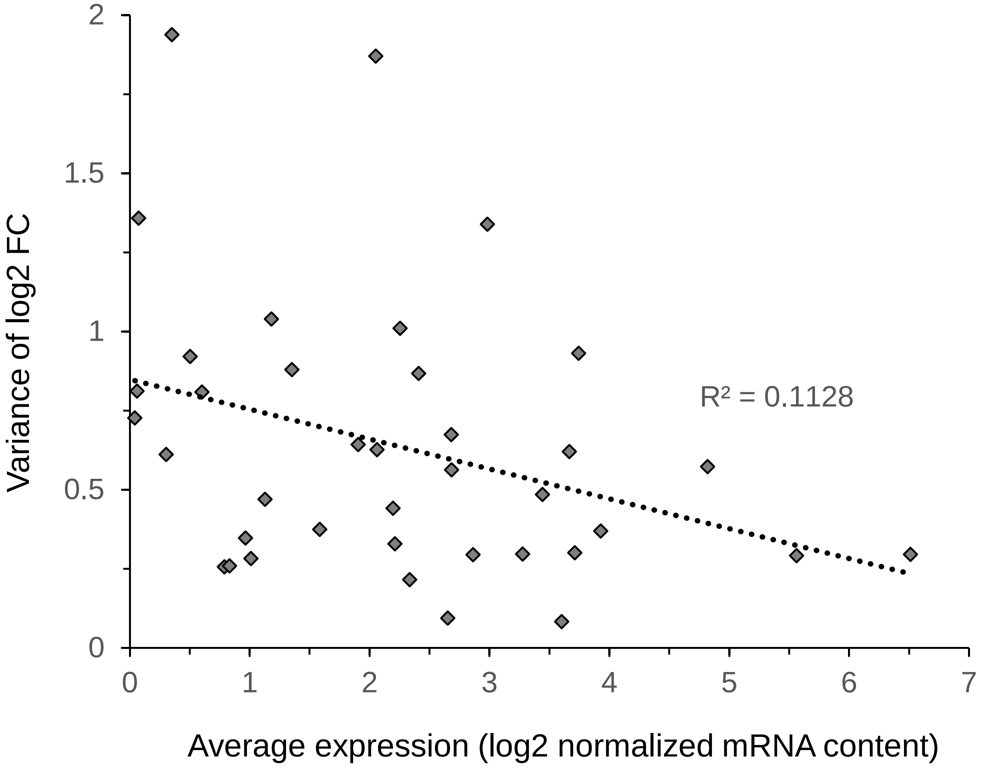


**Supplemental Figure 11: Correlation between change in expression and expression level of the AtUMAMITs.**

The averages of the mRNA levels from RNAseq data retrieved from Genevestigator (see Supplemental Figure 1) were plotted against the variances of the fold change from the AtGenExpresss dataset (hormones, stress, pathogens and root treatment, from Supplemental File 3). Each data point represent one *AtUMAMIT* gene.

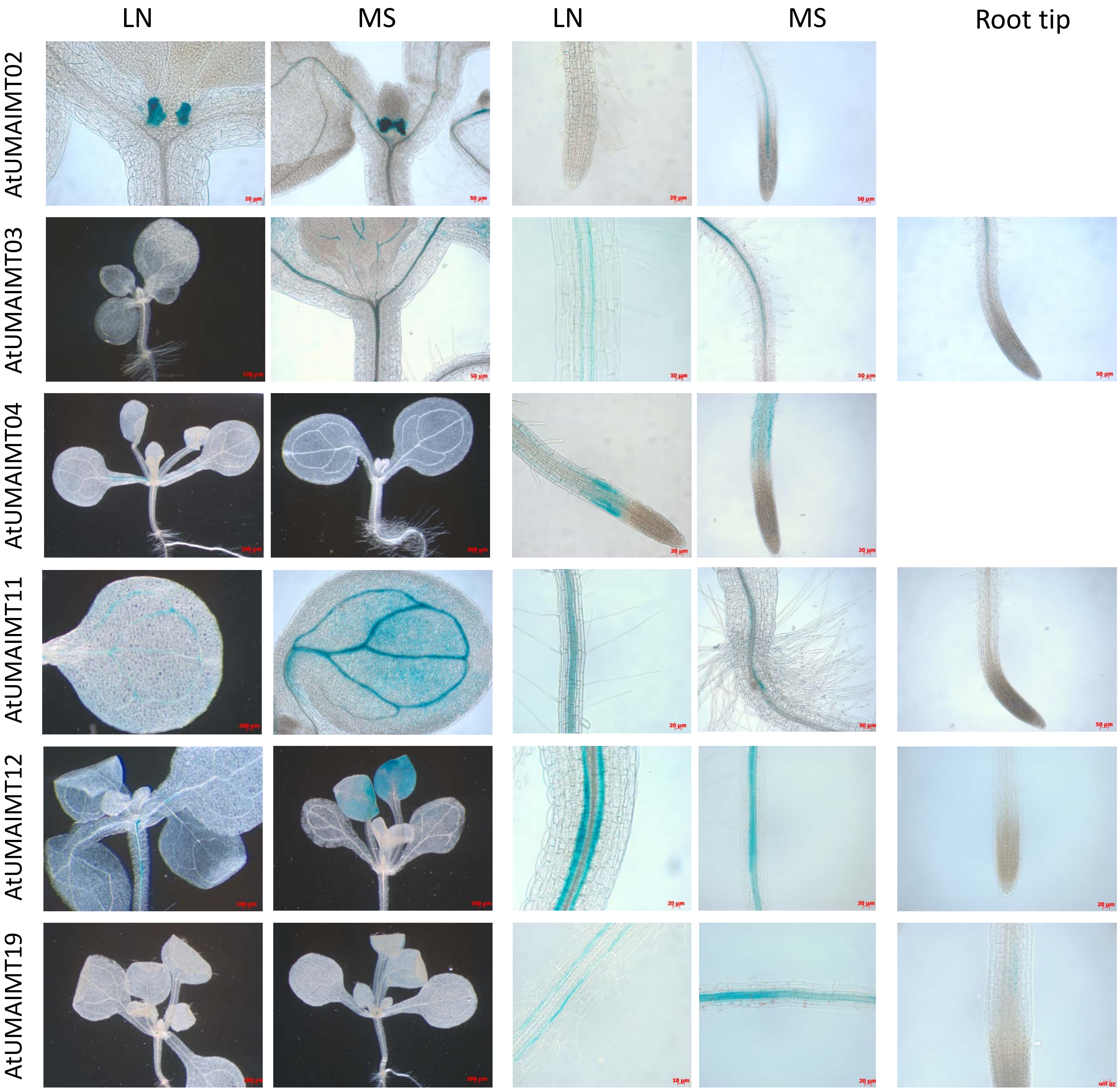
LN
MS
LN
MS
Root tip
AtUMAIMT02
AtUMAIMT03
AtUMAIMT04
AtUMAIMT11
AtUMAIMT12
AtUMAIMT19

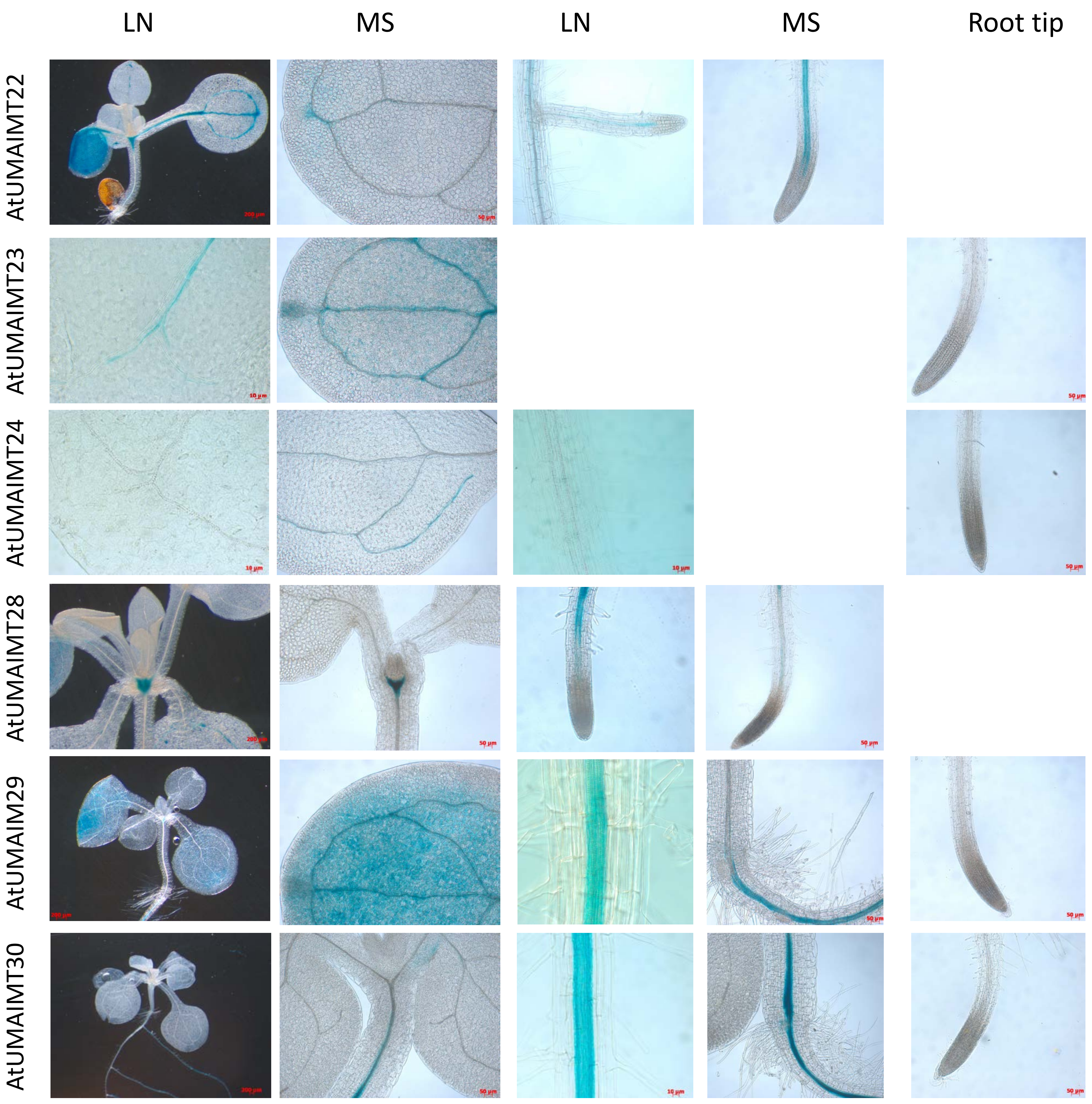
LN
MS
LN
MS
Root tip
AtUMAIMT22
AtUMAIMT23
AtUMAIMT24
AtUMAIMT28
AtUMAIM29
AtUMAIMT30

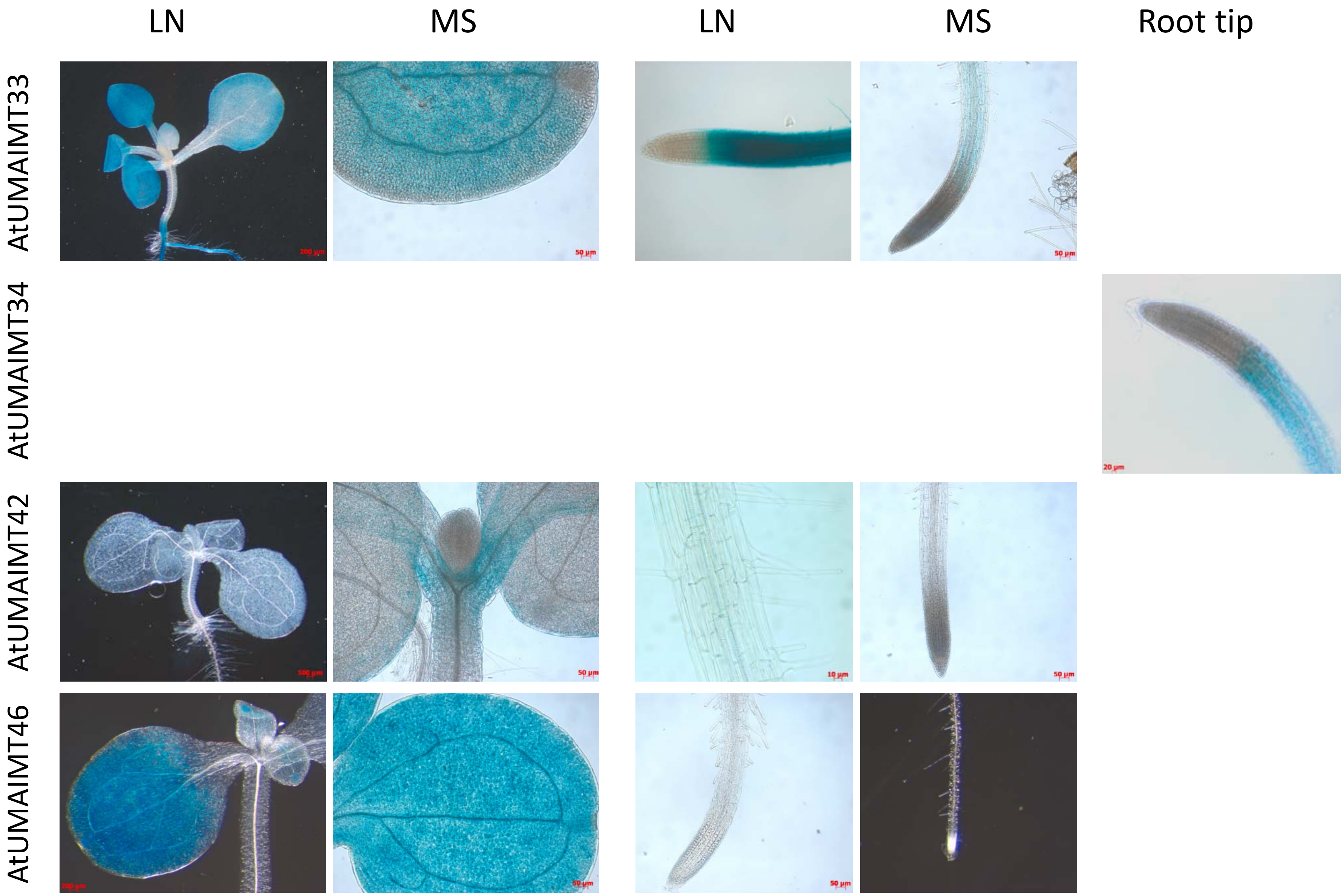


**Supplemental Figure 12. Localization of GUS activity of plants expressing GUS under the control of AtUMAMIT promoters.**

Plants were grown in vitro on media containing low nitrogen (LN) or high nitrogen (1/2 MS medium - MS). For each gene, the staining reaction of plants grown on the LN or MS media was let proceed for the same amount of time, but the reaction time can be different from one gene to another.

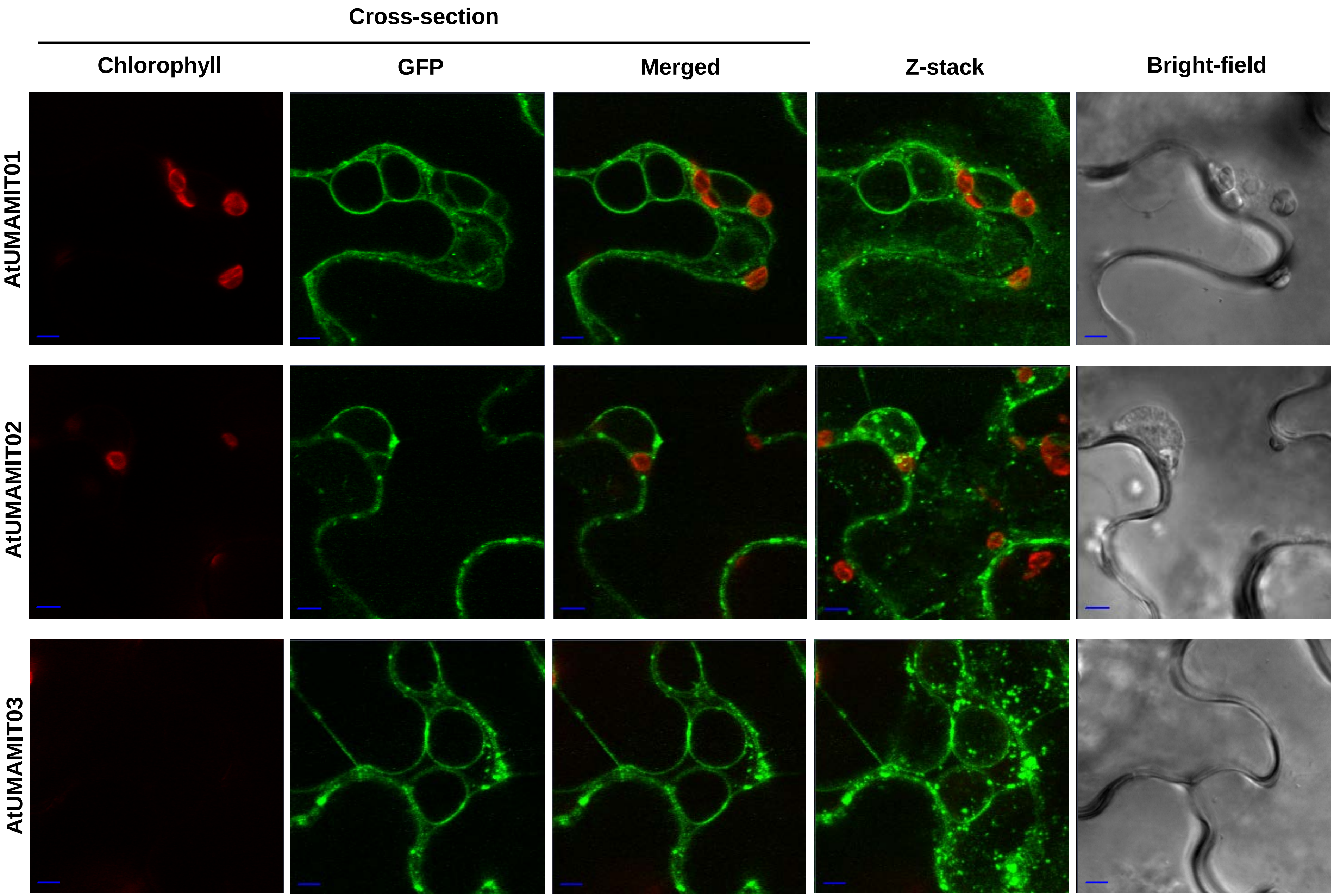
Cross-section
Chlorophyll
GFP
Merged
Z-stack
Bright-field
AtUMAMIT01
AtUMAMIT02
AtUMAMIT03

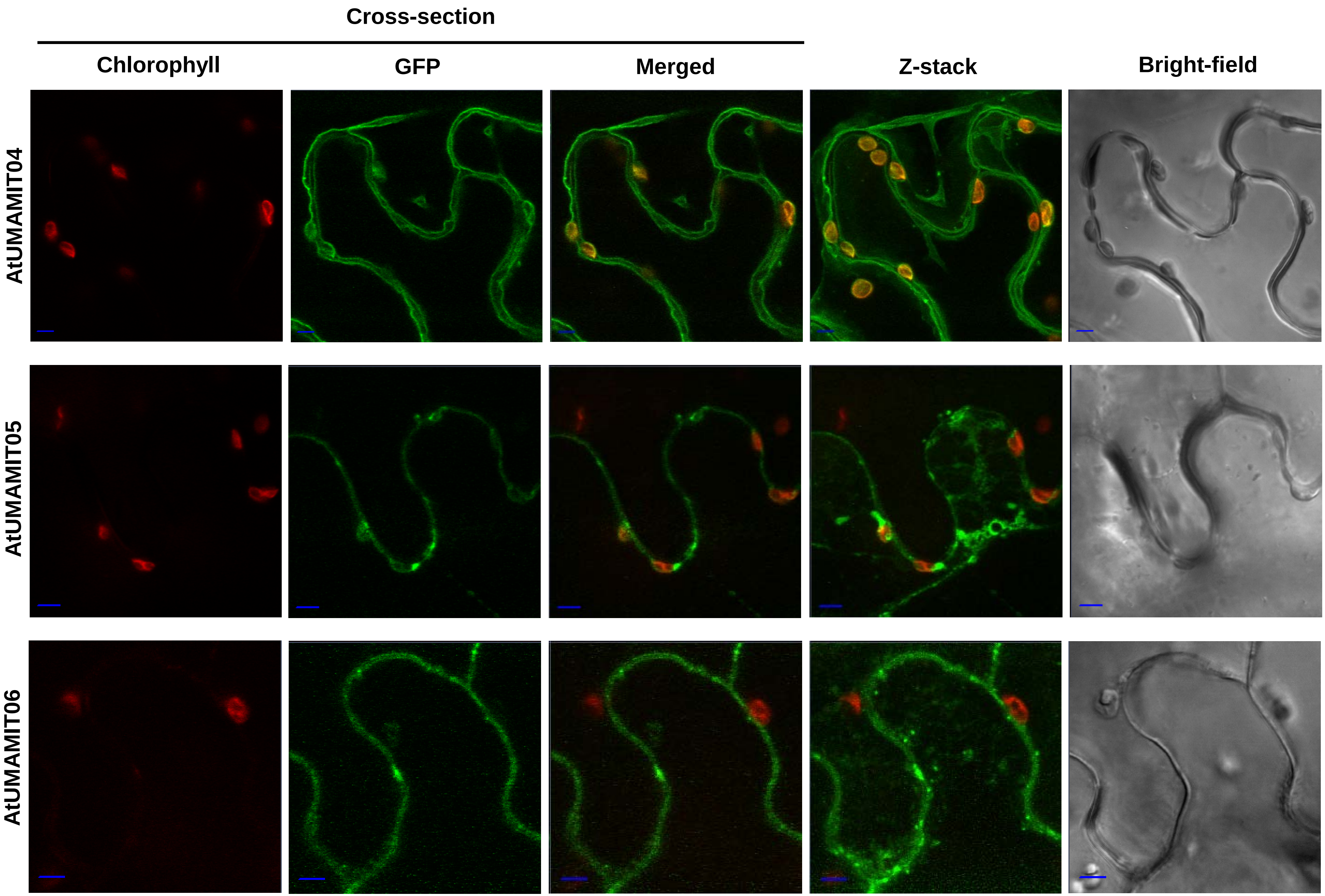

Cross-section
Chlorophyll
GFP
Merged
Z-stack
Bright-field
AtUMAMIT04
AtUMAMIT05
AtUMAMIT06

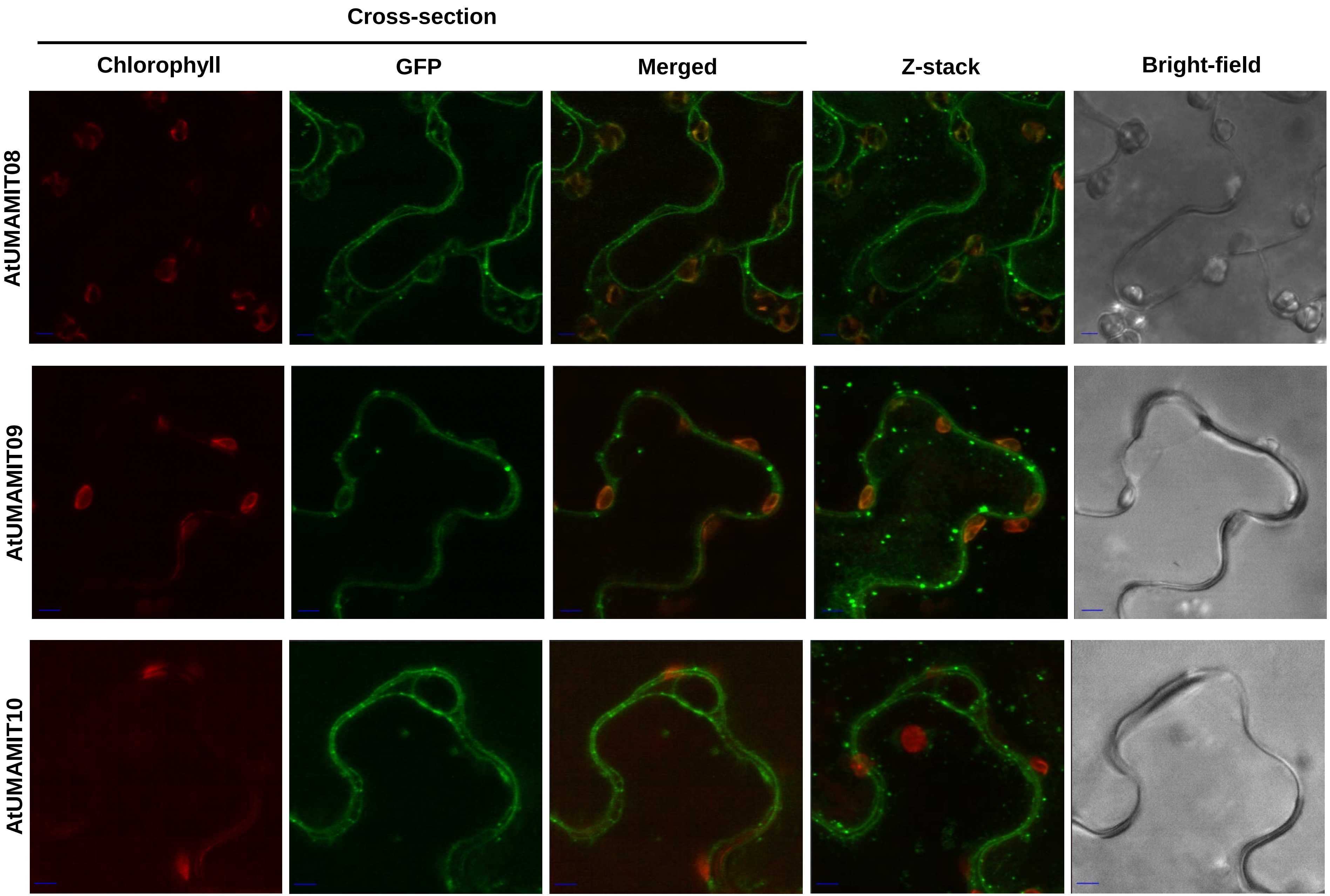

Cross-section
Chlorophyll
GFP
Merged
Z-stack
Bright-field
AtUMAMIT08
AtUMAMIT09
AtUMAMIT10

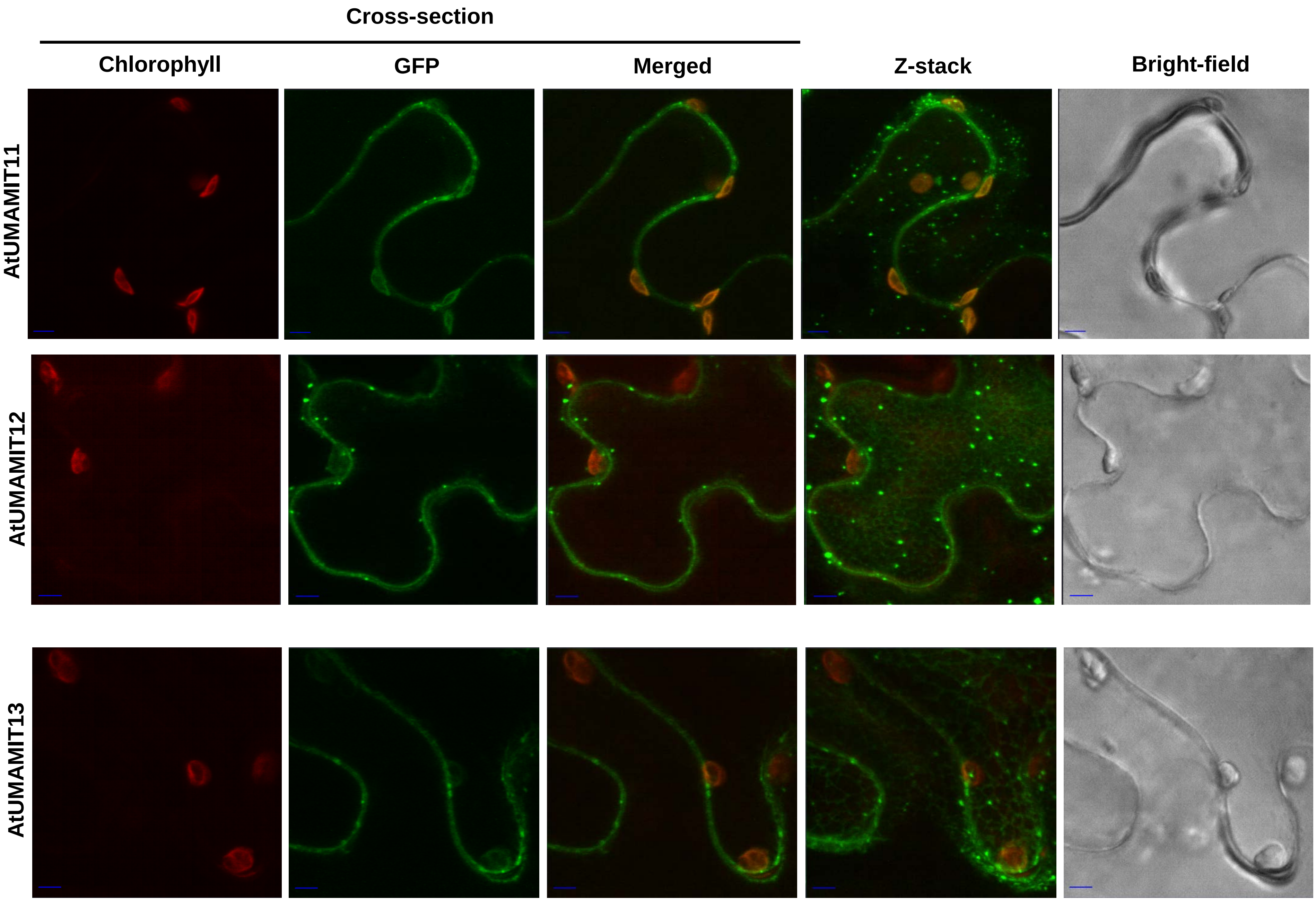

Cross-section
Chlorophyll
GFP
Merged
Z-stack
Bright-field
AtUMAMIT11
AtUMAMIT12
AtUMAMIT13

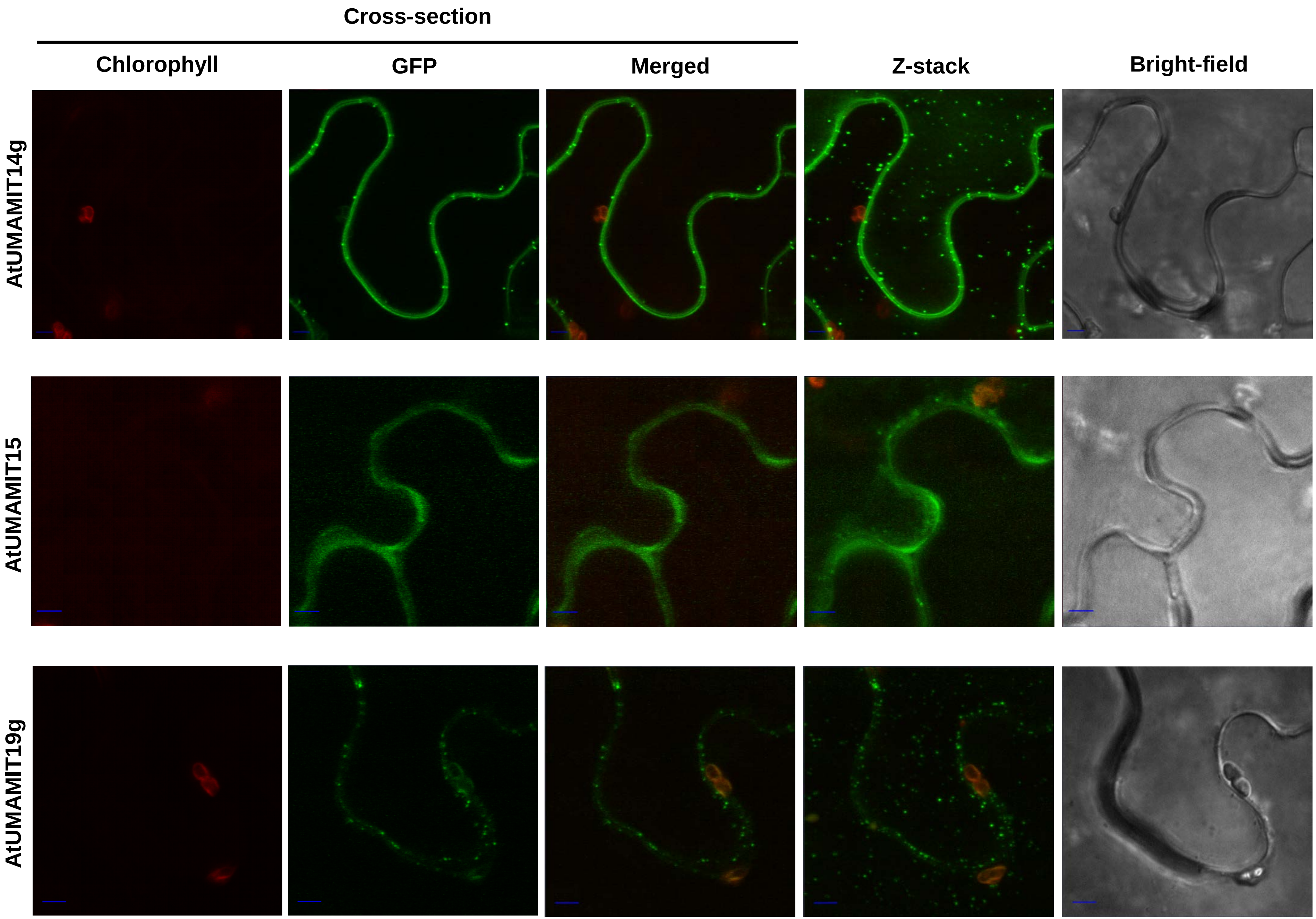
Cross-section
Chlorophyll
GFP
Merged
Z-stack
Bright-field
AtUMAMIT14g
AtUMAMIT15
AtUMAMIT19g

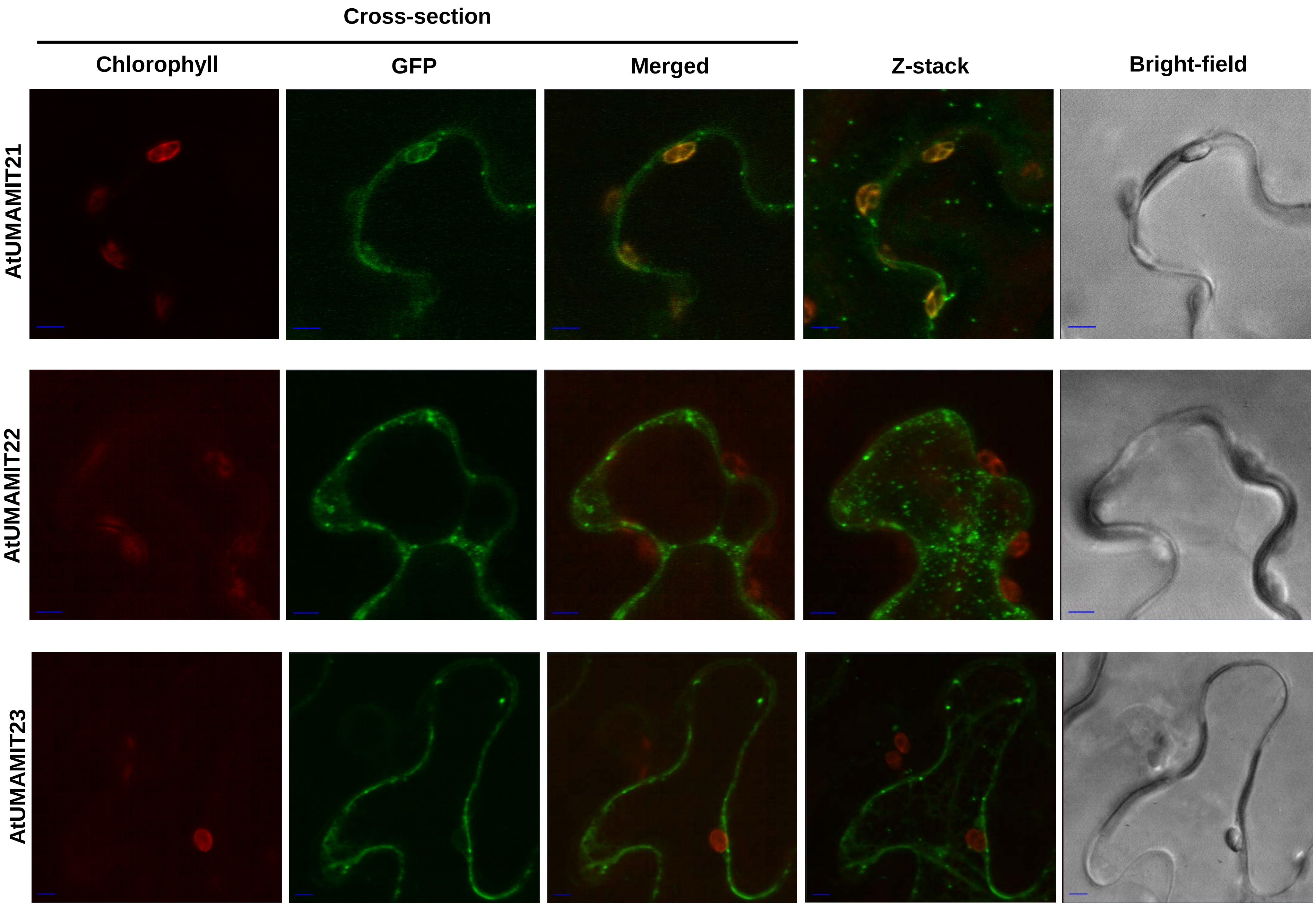

Cross-section
Chlorophyll
GFP
Merged
Z-stack
Bright-field
AtUMAMIT21
AtUMAMIT22
AtUMAMIT23

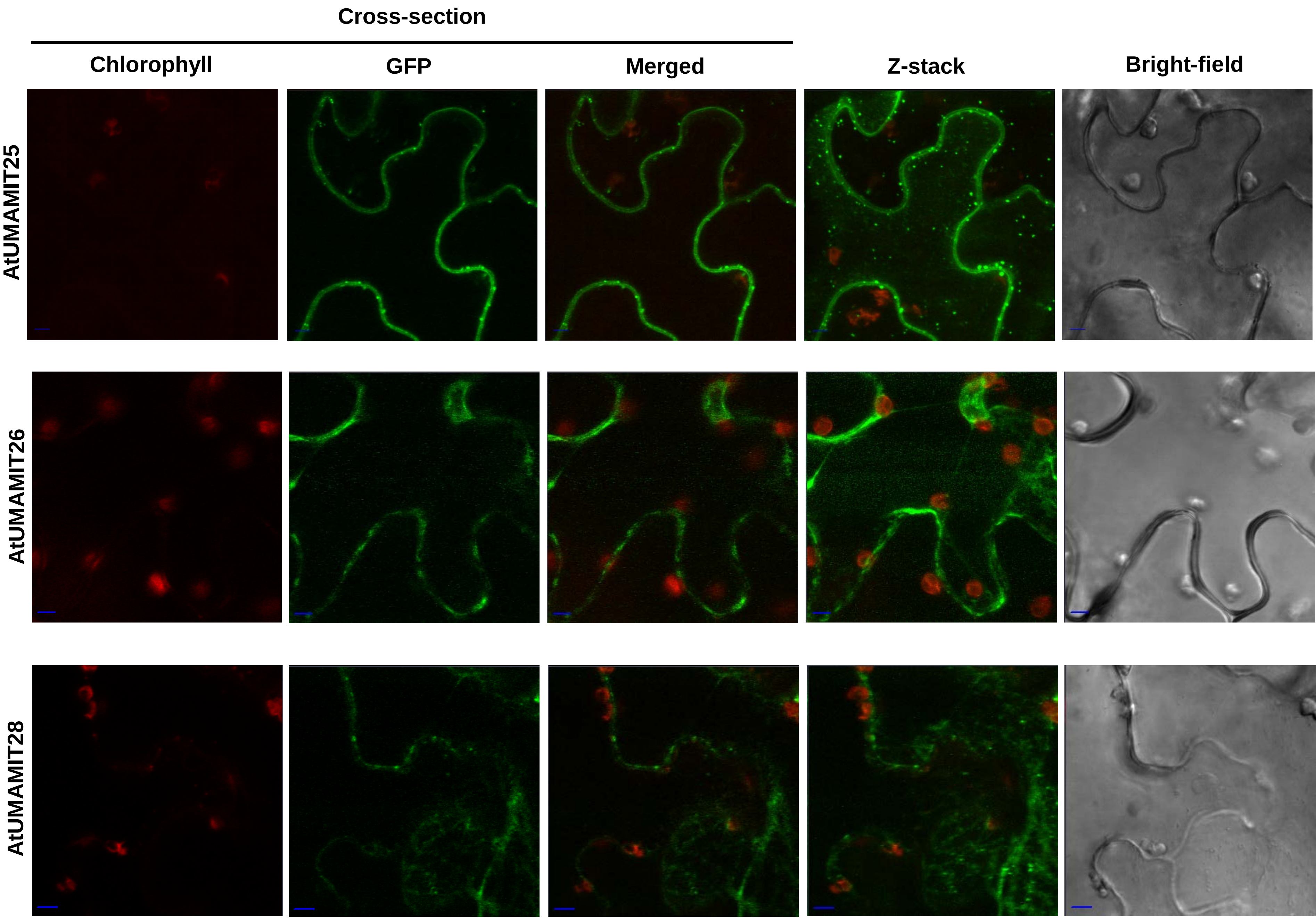
Cross-section
Chlorophyll
GFP
Merged
Z-stack
Bright-field
AtUMAMIT25
AtUMAMIT26
AtUMAMIT28

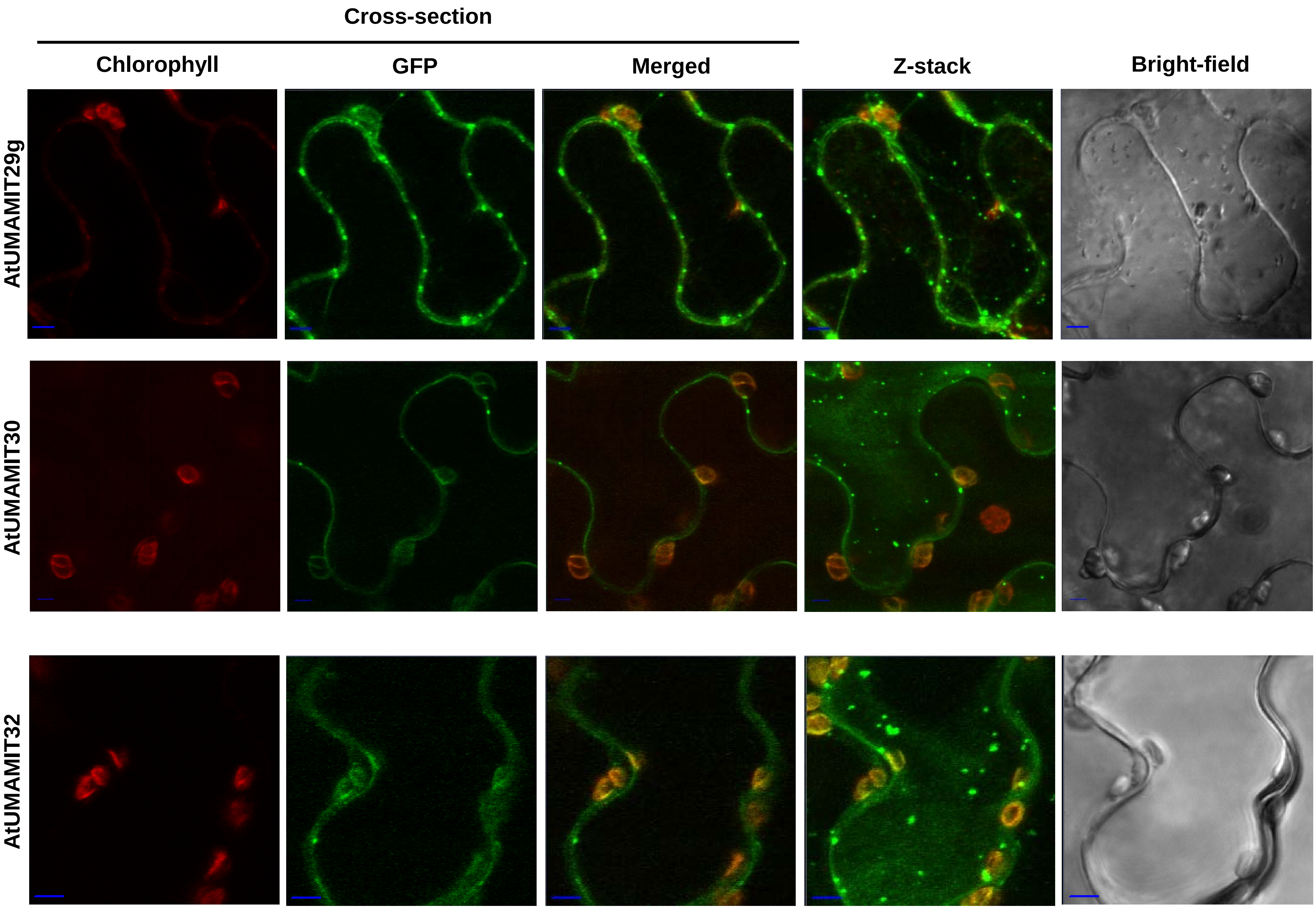
Cross-section
Chlorophyll
GFP
Merged
Z-stack
Bright-field
AtUMAMIT29g
AtUMAMIT30
AtUMAMIT32

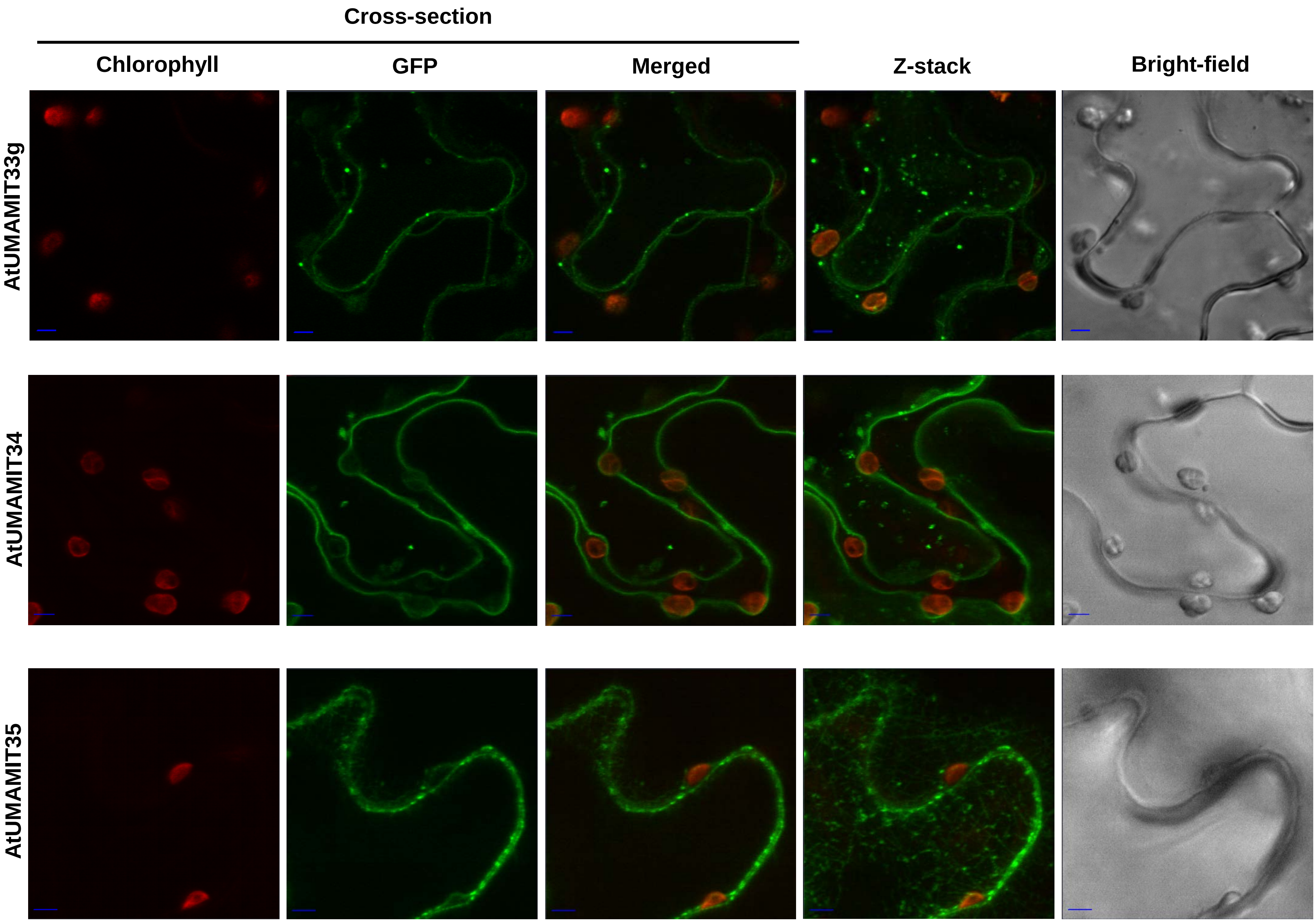
Cross-section
Chlorophyll
GFP
Merged
Z-stack
Bright-field
AtUMAMIT33g
AtUMAMIT34
AtUMAMIT35

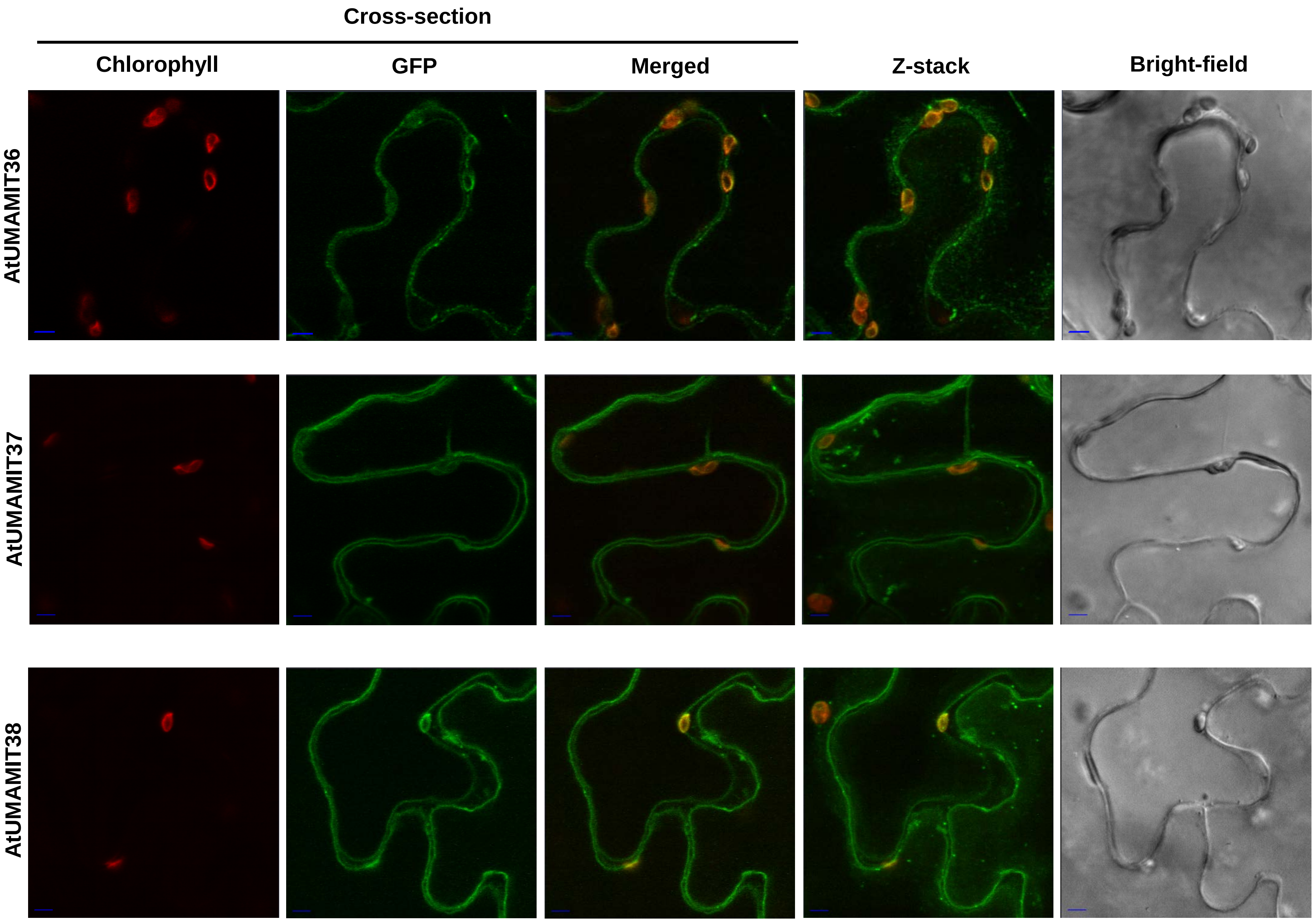

Cross-section
Chlorophyll
GFP
Merged
Z-stack
Bright-field
AtUMAMIT36
AtUMAMIT37
AtUMAMIT38

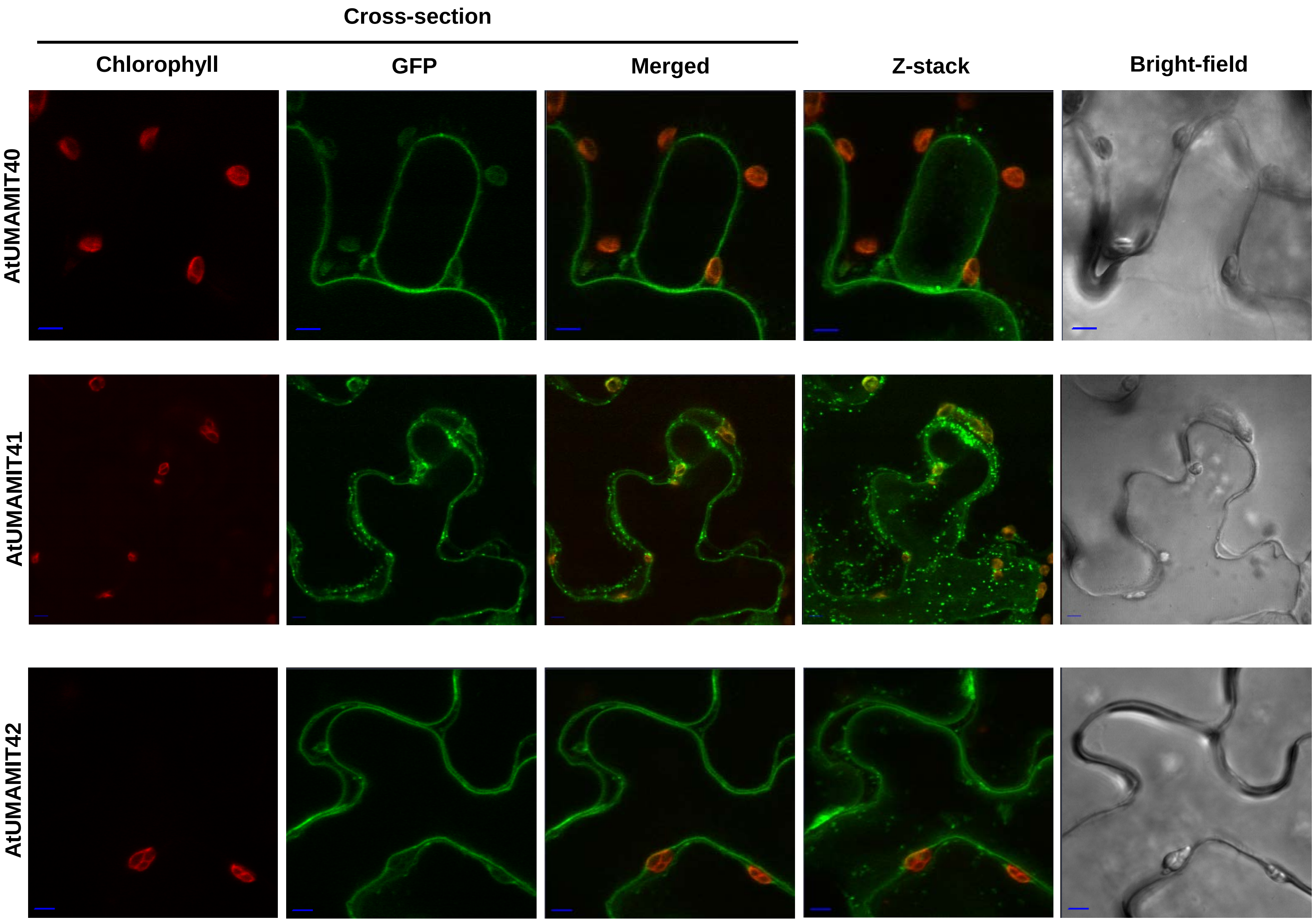
Cross-section
Chlorophyll
GFP
Merged
Z-stack
Bright-field
AtUMAMIT40
AtUMAMIT41
AtUMAMIT42

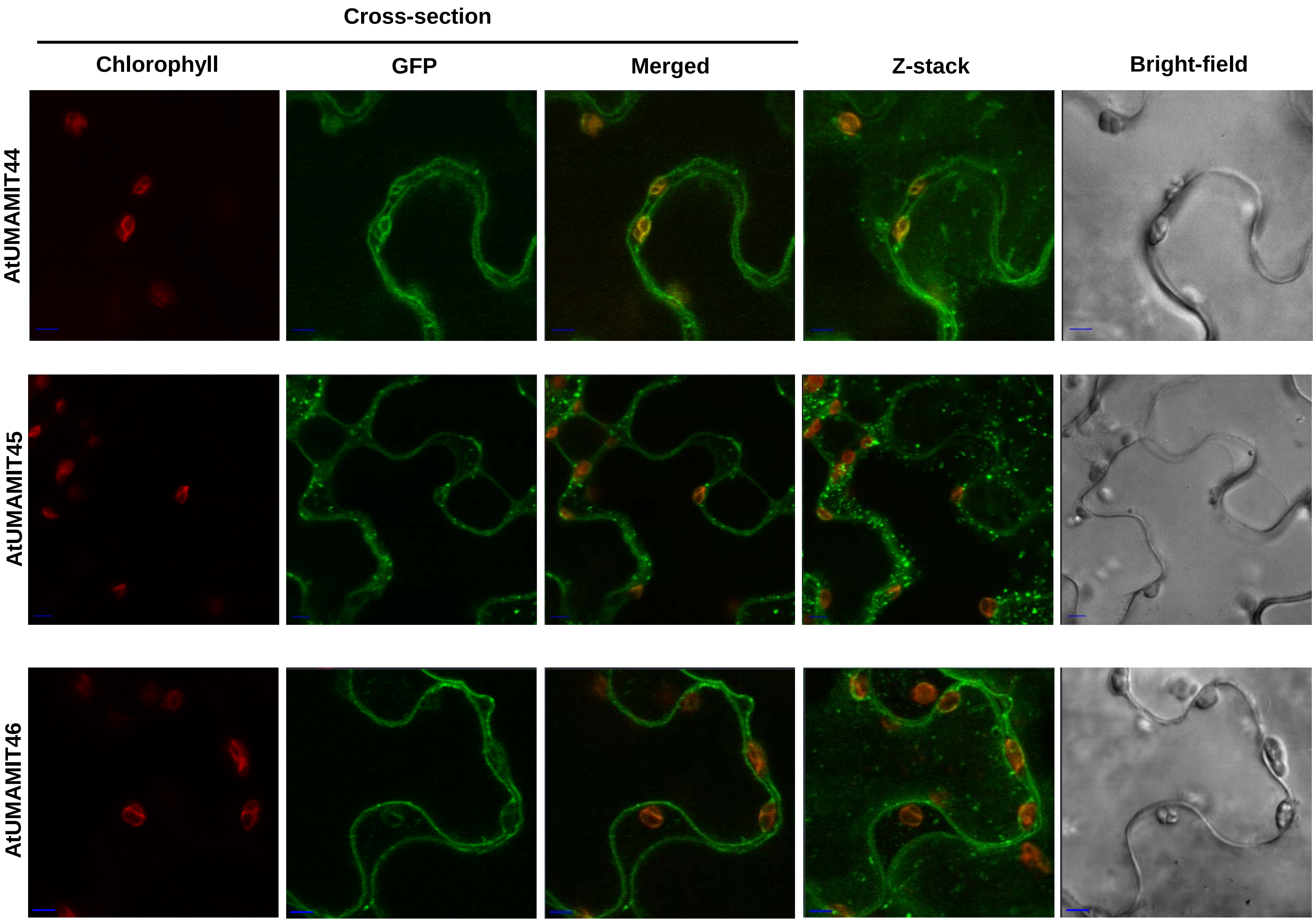
Cross-section
Chlorophyll
GFP
Merged
Z-stack
Bright-field
AtUMAMIT44
AtUMAMIT45
AtUMAMIT46

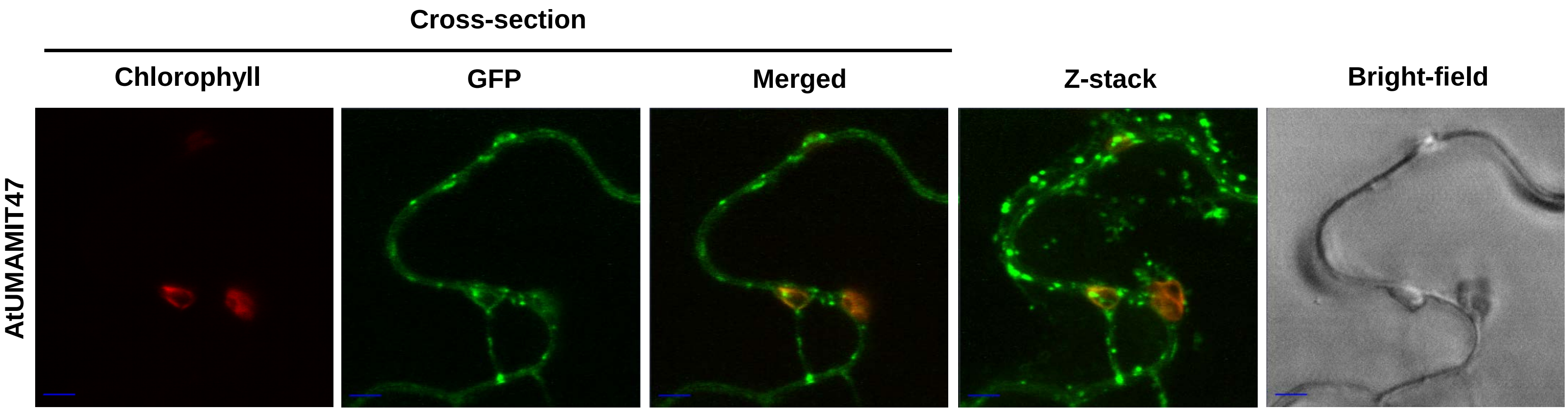


**Supplemental Figure 13: Subcellular localization of the AtUMAMITs proteins expressed in *N. benthamiana cells*.**
AtUMAMITs were expressed with the GFP fused at their C-terminus in *Nicotiana benthamiana* leaf epidermis, and observed by confocal miscroscopy. A z-stack of at least 12 sections was constructed; cross-section images were taken from the z-stack. Scale bar = 5 µM.

**A**

```
ATUMAMIT01----------MAAPAILNG-GDATERETRMAHSA
ATUMAMIT02----------MTAPMILTGSGSAAERDARMAHTA
ATUMAMIT03-------------------MESTVEREAWKAHVA
ATUMAMIT04-----------------MGKGV--VSEKVKLVVA
ATUMAMIT05-----------MADNTDNRRSLWGVPEKLQLHIA
ATUMAMIT06---------------------MSPIPERAKLHIA
ATUMAMIT07-MQFFCVNLYRSVLNLLEERMKTEMIEEMVIVGG
ATUMAMIT08-------------------MMKRETLIEAGIIGG
ATUMAMIT09-------------MSMEEISSCESFLTSSKPYFA
ATUMAMIT10--------------------MGLKMSESAKPYFA
ATUMAMIT11--------------------MGLRMSESAKPYFA
ATUMAMIT12----------------MEEVKKRDCMEKARPFIS
ATUMAMIT13----------------------MACMKKALPFIL
ATUMAMIT14-----------------------MALKTWKPFIT
ATUMAMIT15-----------------------MKFERARPFIA
ATUMAMIT17--------------------MKGGKMDKLKPIIA
ATUMAMIT18--------------------MKGGSMEKIKPILA
ATUMAMIT19--------------------MGRGLMNSLKPYLA
ATUMAMIT20------------------MEGVSATMHKLRPYLL
ATUMAMIT21-----------------------MDMESKKPYLM
ATUMAMIT22-----------------------MMMEHKANMAM
ATUMAMIT23----------------------------MKDITA
ATUMAMIT24----------------------------MKSVVA
ATUMAMIT25-----------------------MAKSDMLPFLA
ATUMAMIT26---------------MGEDMRVVKVESKWPPIIV
ATUMAMIT27---------------MGEDMRGVKVVSKWPPMIV
ATUMAMIT28---------------MAGDMQGVRVVEKYSPVIV
ATUMAMIT29----------------------MMKEEQWAPVIV
ATUMAMIT30---------------------MGFIDGKWAPMIV
ATUMAMIT31---------------------MGYCDGKWTPVII
ATUMAMIT32--------------------MVKFDTKLWKAVLM
ATUMAMIT33-----------------------MEISKYKAVLA
ATUMAMIT34----------------------MGKIEEYKPVMA
ATUMAMIT35------------------MGLMDTRWETIVPFIV
ATUMAMIT36-------------------MEVKVRRDELVPFVA
ATUMAMIT37----------MTRGAGGETVAWSYFCRDVVPFTA
ATUMAMIT38-----------MR---EETVSWKYFKRDVVPFTA
ATUMAMIT40-----------MREAGEEKVAWKYFTRDVVPFAA
ATUMAMIT41-------------------MARKYFQREVLPVTA
ATUMAMIT42------------------MVHGRLCNRDGWILTA
ATUMAMIT44-----------------MA-SITLRRRDAVLLTA
ATUMAMIT45-----------------MARTVSLWRREAVFLTA
ATUMAMIT46--------------MPIMAGTASPWRREAVFLTA
ATUMAMIT47-----------------MAGAVSLWRREAVFLTA
```

**B**

| | |
|---|---|
| **ATUMAMIT01** | WASYREQQTTSAGNEIASSSDVRISEPFIYRDETGK |
| **ATUMAMIT02** | WASFRERKTAVSGIGIAPHGLKTSEPLIFNGTVNRLGQLFSGLPSSSVKSAD |
| **ATUMAMIT03** | WASYKEKKAAAAMAVIPITSKEAEPLIYKDHKNKPIGHLFTKSPISSPKSDD |
| **ATUMAMIT04** | WGKNEERKLALEESQQDPESLTKHLLEAQHKKSNSESEV |
| **ATUMAMIT05** | YGKSEERKFAALEKAAIQSSAEHGIERAPVSRNSIKSSITTPLLHQSTDNV |
| ATUMAMIT06 | MGKSWENQALCQQQQHMISSAASDFGDEEDYHNNKPRSPISQPLISS |
| ATUMAMIT07 | WAKGKEGFSEIESFESEFDSKKPLLS |
| **ATUMAMIT08** | WAKGKEDCEEIDEMKQDDEESLLRTEFDLQKPLLL |
| ATUMAMIT09 | WGKQKENQVTICELAKIDSNSKVTEDVEANGSKMKISEGDNSMLSTIVISVPLSETHLKKTIQEP |
| **ATUMAMIT10** | WGKEGDIDEEENIEEKFVEIVKCCNRCDIKVLSMMPRIDEEVDVEMQSAGTAKVAVGFS |
| ATUMAMIT11 | WGKHVDDDGEETRHEDNVVAVKCCSGNNGLTIMPKIDEADEEDVETGKATSEKESSVPEVVVVVFCSENVHSVSRF |
| ATUMAMIT12 | WGKGKDYKYNSTLQLDDESAQPKLELSGNGKDNVDHEVITISKQGEQRRTAVETV |
| ATUMAMIT13 | WGKAKDYEYPSTPQIDDDLAQATTSKQKEQRRTVIESV |
| ATUMAMIT14 | WGKSKDEPSSSFSDMDKELPLSTPQIVLPSKANAKMDTNDASVVISRPNTNESV |
| ATUMAMIT15 | WGKSKDKGGMLQPNAGCAETVVKIDQQKVPTPDNNQVVSISYHLMIPKAAARSQE |
| ATUMAMIT17 | WGKAKDEVISVEEKIGMQELPITNTSTKVEGGGITSEVNEGVTNNTQV |
| ATUMAMIT18 | WGKSKDEVNPLDEKIVAKSQELPITNVVKQTNGHDVSGAPTNGVVTST |
| ATUMAMIT19 | WGKGKDKRMTDDDEDCKGLPIKSPVKPVDTGKGLAAELEMKSKEGQEAKATTTTQVEI |
| **ATUMAMIT20** | WGKGKDYEVSGLDILEKNSLQELPITTKSEDDNKLVSSISDNSNVTIPGGAHSNTSGI |
| ATUMAMIT21 | WGKSREEKNSGDDKIDLQKENDVVCNEVKVVIS |
| ATUMAMIT22 | WGKTKEEEIQRYGEKQSQKEIIEEVIIV |
| ATUMAMIT23 | WAKNKEMKSMLTTSDHNETNKTSKDITVNNLPTLSTNVP |
| **ATUMAMIT24** | WCKMKEKKSASTTSDHIETNKNNKELDLGNLSSVNRDVP |
| ATUMAMIT25 | WGKDREVSEKEEEREKVKQQNHKVKSESNEDIESRLPVASSGNGSTRSTSP |
| ATUMAMIT26 | WGKNKDMEA |
| ATUMAMIT27 | WGKNKETEADITTLSSRMNNEDQRV |
| ATUMAMIT28 | WGKNKETESSTALSSGMDNEAQYTTPNKDNDSKSPV |
| ATUMAMIT29 | WGRKNETDQSVSKTLNSSQFSQNKDNEDHTIANHKDTNLPV |
| ATUMAMIT30 | WSRSKQIVECKIMKLPTNTVEEEKEEEGRTNVNNGQLLVIPMTP |
| ATUMAMIT31 | LGKVRLMKEECEKKLPCRFNEDDQEEDDDEQYKKGHLMVVPMTP |
| **ATUMAMIT32** | WGKSKDKSASVTKQEPLDLDIEGCGTAPKELNSTAHQVSAK |
| **ATUMAMIT33** | WGKSEDYQEESTDLKLENEHNTSSQSDIVSIMIGDKAFRSSELLEPLLM |
| **ATUMAMIT34** | WGKAKDVMMNQDQRDNDQKSEVKIHIEDSSNTTICNKDLKNPLLSKHKSTEEIQTHQQLY |
| ATUMAMIT35 | WSQVQKDDPNETVEKNDNHQLDSDEQTTPLLLANGDFDQV |
| ATUMAMIT36 | WGQLKESEEKQSSNEERKSIKTIHHRDEDEYKVPLLINQEESPV |
| **ATUMAMIT37** | WGKAREDSIKTVAGTEQSPLLPSHTIEEEASLSCD |
| **ATUMAMIT38** | WGKAREDSTKTVSDSEQSLLLPSHDREED |
| **ATUMAMIT40** | WGKAREDTIKTVAGSEQSPLLLTHIIEDGAFPLS |
| **ATUMAMIT41** | WGKAKEVALVEDDNKANHEEANEADLDSPSGSQKAPLLESYKNDEHV |
| **ATUMAMIT42** | WGKAKEDKVDIIGAIESSPSHNAPLLDNFKS |
| **ATUMAMIT44** | WGKAKEGKTQFLSLSEETPLLDENIDDRI |
| ATUMAMIT45 | WGKANEEKDQLLLVSGKERTPLLLNG |
| **ATUMAMIT46** | WGKANEEKDQLSFSEKEKTPLLLNRKNDQV |
| ATUMAMIT47 | WGKANEEKNKLLSFSGKEKTPLLLSGKNDQI |

**Supplemental Figure 14: Alignment of the N- and C- termini of the AtUMAMITs**

A- Alignment of the sequences from the beginning of the protein to the transmembrane domain A1 (N-terminus)of the AtUMAMITs.

B- Alignment of the sequences from the end of the transmembrane domain B5 to the end of the protein (C-terminus)of the AtUMAMITs

Acidic residues are in red, aromatic residues in green, and Leu and Ile in blue. Proteins detected at the tonoplast are in bold, while proteins detected at the plasma membrane are underlined (see Table 1).

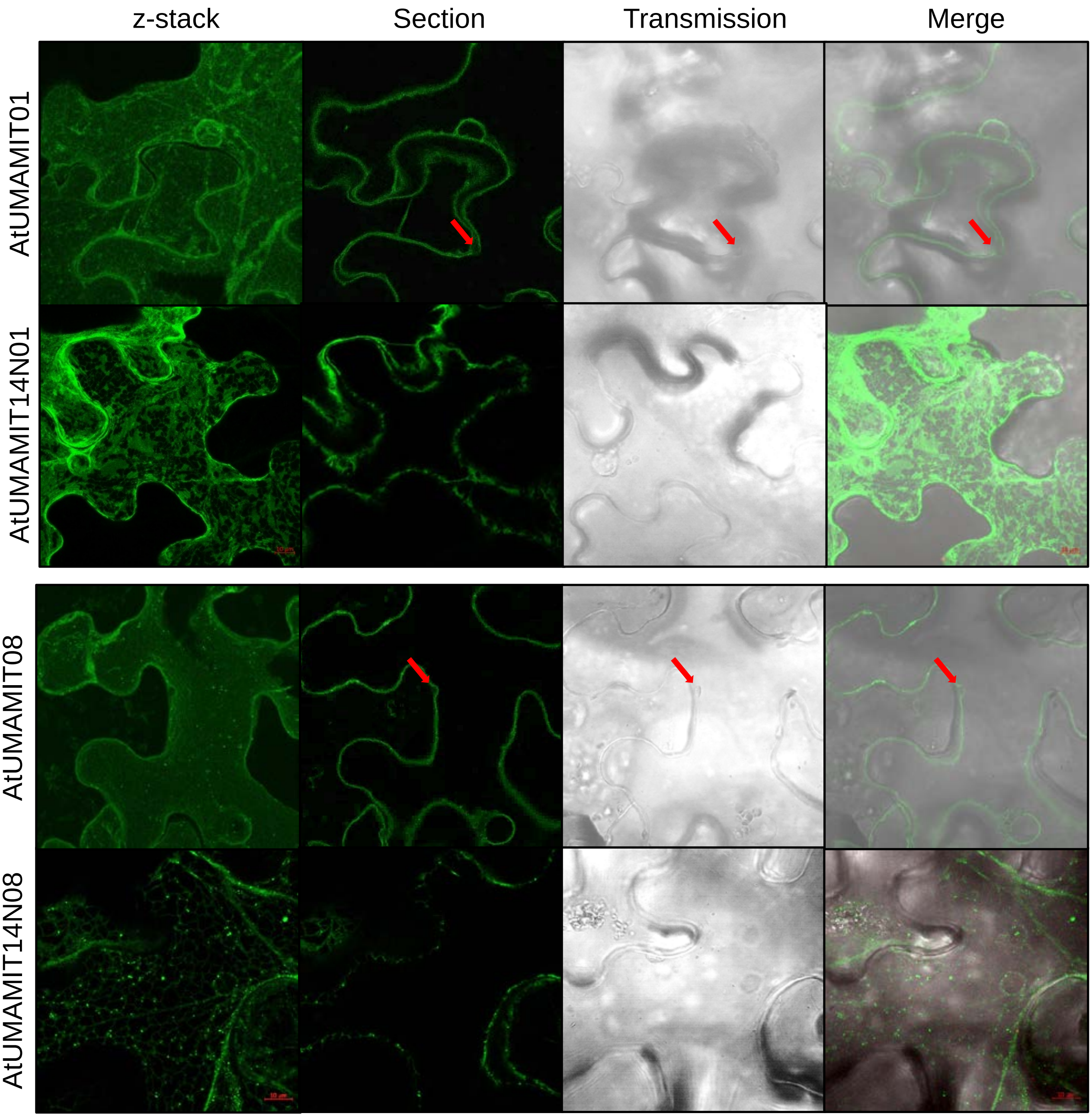
z-stack
Section
Transmission
Merge
AtUMAMIT01
AtUMAMIT14N01
AtUMAMIT08
AtUMAMIT14N08

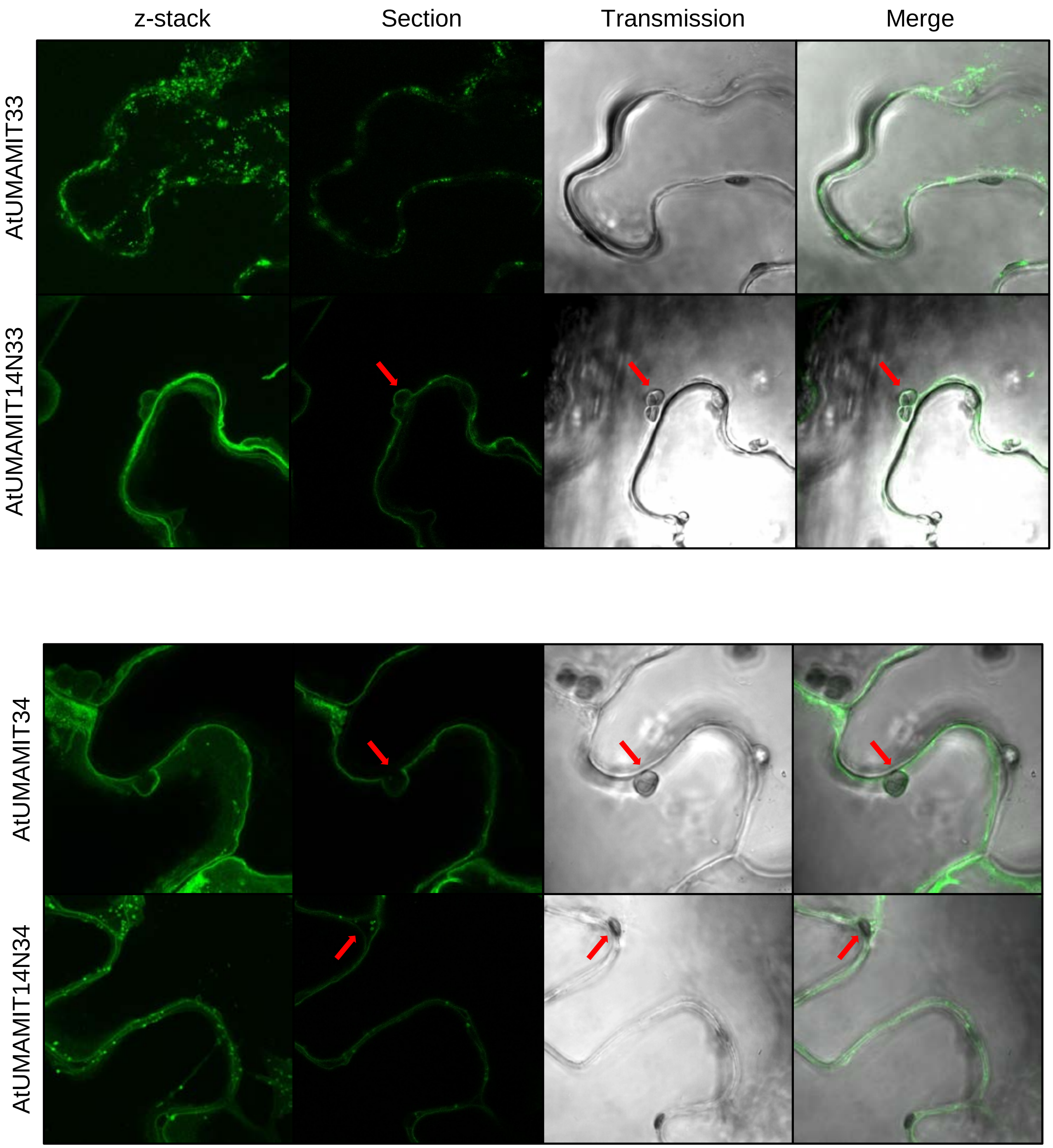
z-stack
Section
Transmission
Merge
AtUMAMIT33
AtUMAMIT14N33
AtUMAMIT34
AtUMAMIT14N34

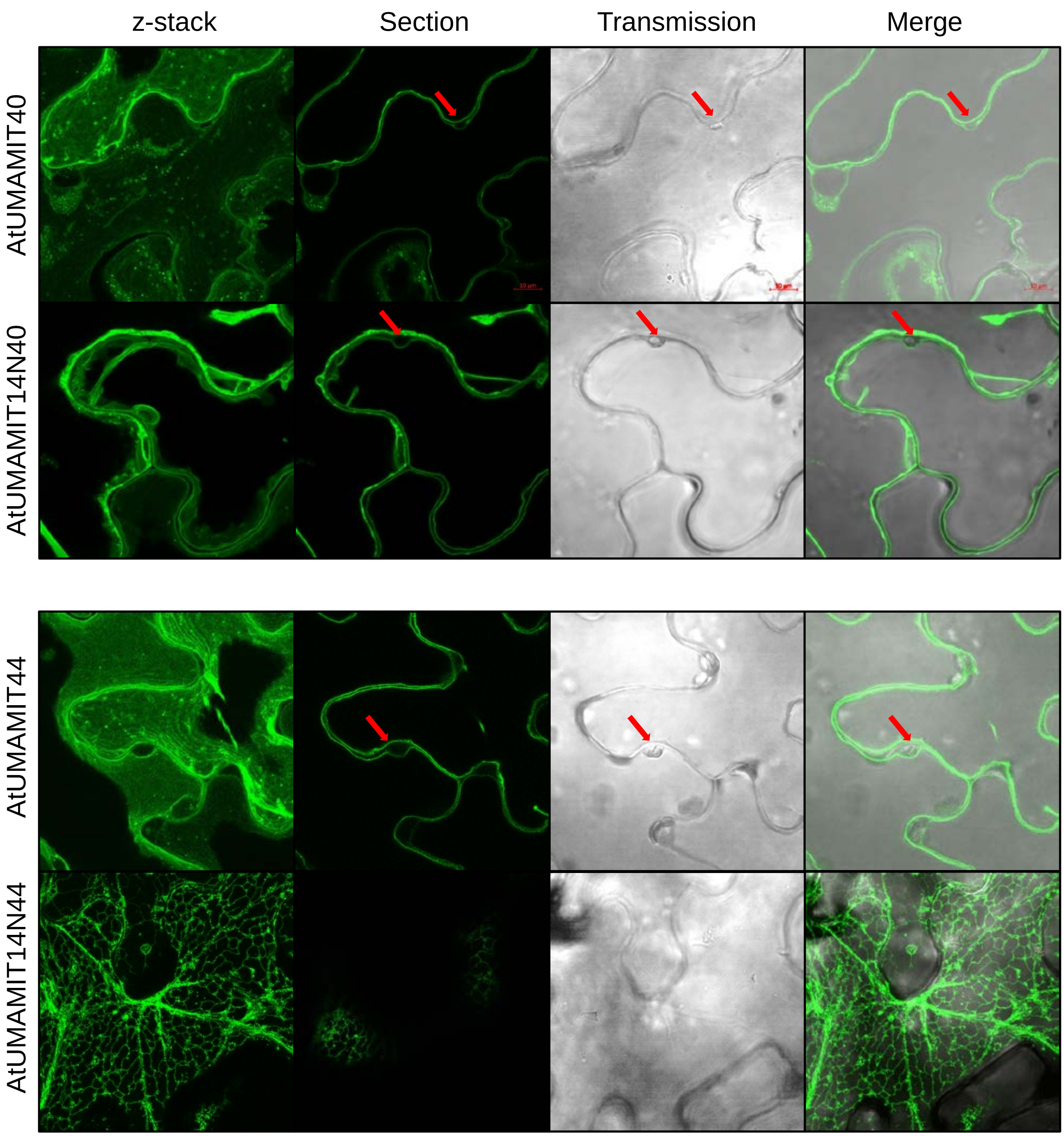


**Supplemental Figure 15: Subcellular localization of AtUMAMIT proteins without and with N-terminus exchange.**

Each protein was expressed in *N. benthamiana* epidermis cells with the YFP fused to their C-terminus. For each studied protein, the unmodified protein localization is presented on top, and the modified protein (for which the N-terminus of AtUMAMIT14 replaced the native N-terminus) at the bottom of each panel. The section and transmission images were obtained from the z-stack, for which a projection was created to show the pattern of the fluorescence in the cell (z-stack). Red arrows point to chloroplasts to emphasize the tonoplast or plasma membrane localization.

A

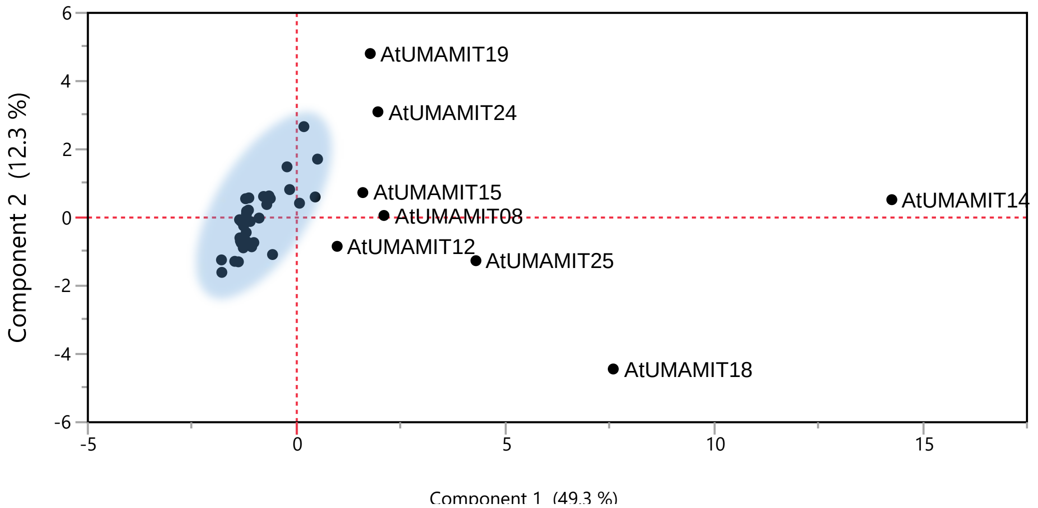


B

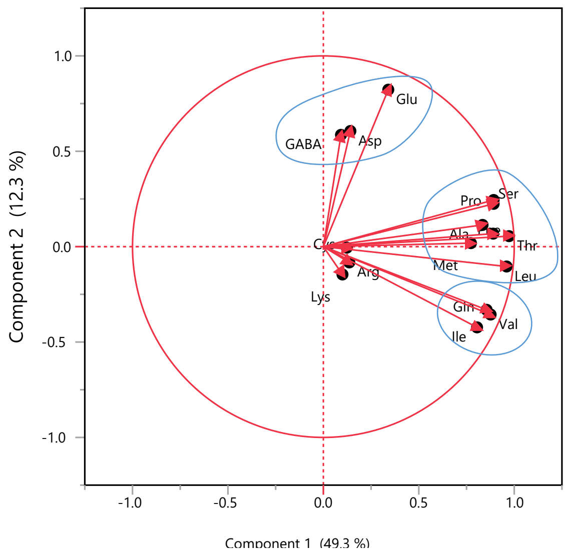


**Supplemental Figure 16. PCA analysis of amino acid export profiles of the AtUMAMIT expressed in yeast.**

Data from Supplemental File 4 were used for analysis in JMP15.

A. PCA plot. The most individualized proteins are labeled.

B. Loading plot with names of the corresponding amino acids. This graph shows three groups of amino acids whose content correlates with one-another inside each group: (1) Glu, Asp and GABA, (2) Gln, Val and Ile, (3) Pro, Ser, Phe, Ala, Thr, Leu and Met.

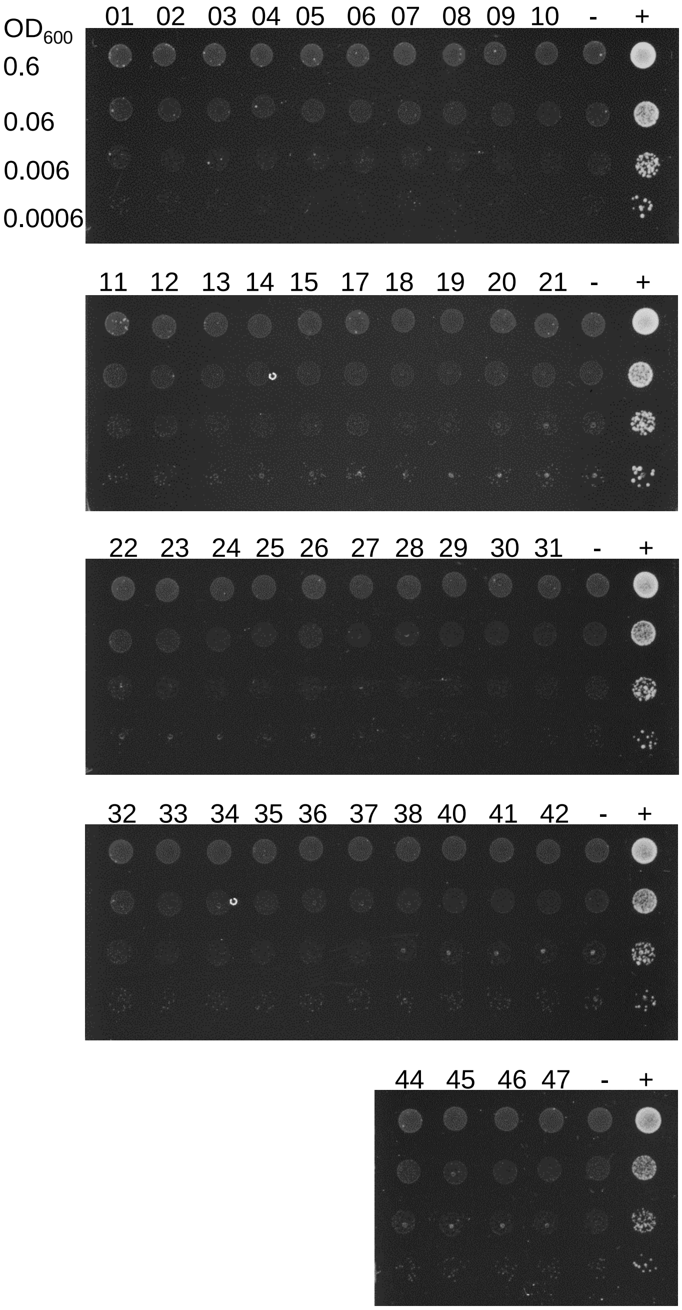


**Supplemental Figure 17. Methionine uptake complementation assay of yeast expressing the AtUMAMITs.**
22Δ10α cells transformed with the AtUMAMITs, or AAP3 (+) or vector only (-) were streaked on plasmid-selective SD medium and grown at 29°C for 3 days, resuspended in $H_2O$, diluted to $OD_{600}$ = 0.6, 0.06, 0.006, and 0.0006, and spotted on assay medium with 3 mM  Met as the sole nitrogen source.

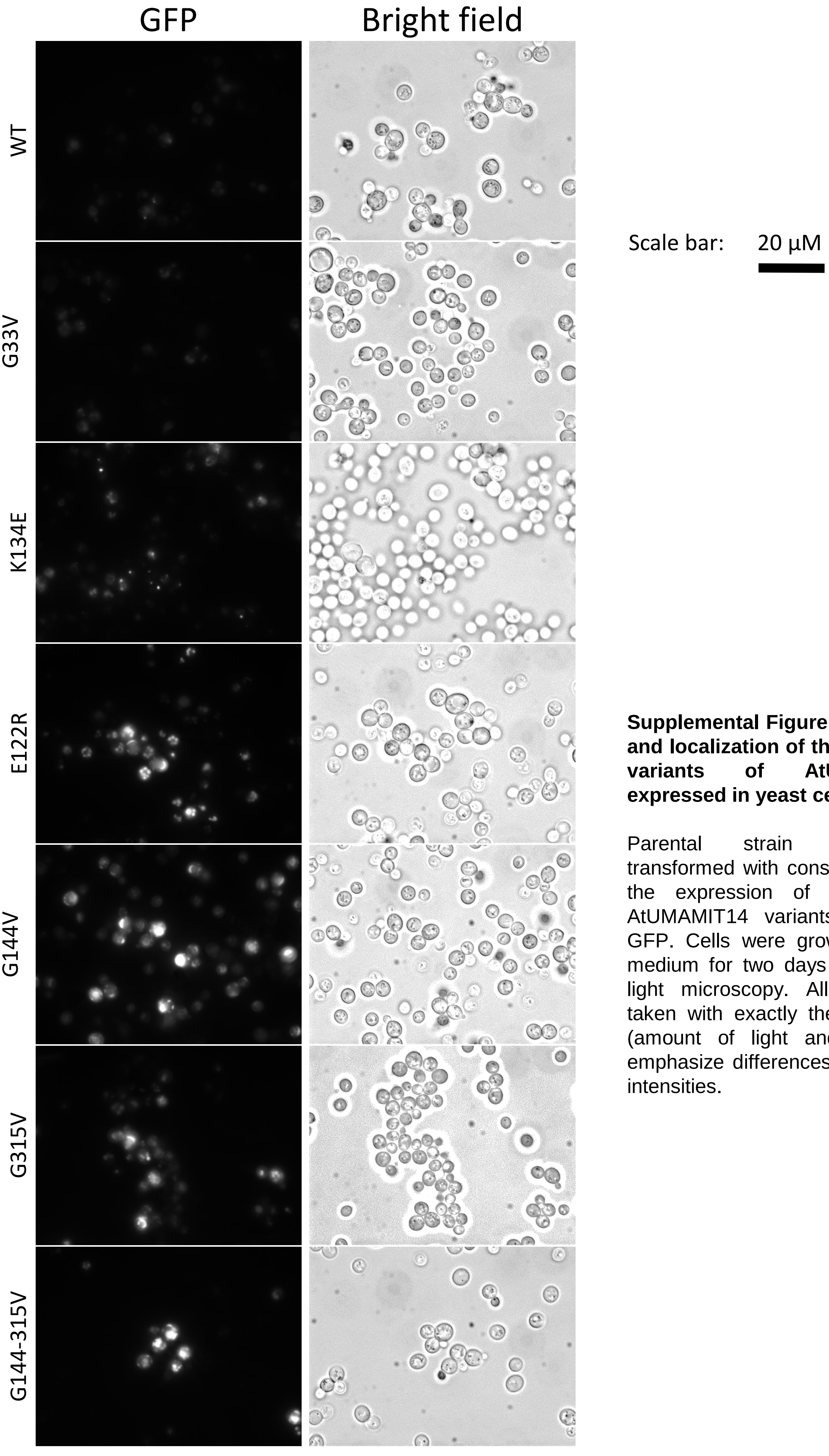


**Supplemental Figure 18. Expression and localization of the wild type and variants of AtUMAMIT14-GFP expressed in yeast cells.**

Parental strain 23344c was transformed with constructs leading to the expression of wild type and AtUMAMIT14 variants fused to the GFP. Cells were grown on solid SD medium for two days and imaged by light microscopy. All pictures were taken with exactly the same settings (amount of light and exposure) to emphasize differences in fluorescence intensities.
