## Supplemental Table for "Detailed characterization of the UMAMITs provides insight into their evolution, functional properties as amino acid transporters and role in the plant"

**Supplemental Table 1. Oligonucleotide sequences**

The reverse oligonucleotides used for the cloning of the AtUMAMIT CDS contain a Y (= C or T) at the position of the stop codon, so that fragments that have a TGG (encodes a Trp in the sense orientation) can be cleaved by *Kpn*I at that site, while the fragment that have a TAG (stop codon in the sense orientation) cannot. This enables the generation of clones with or without stop with one single oligonucleotide, one BP reaction, and screenable by restriction analysis with *Kpn*I.

| Name | Sequence (5'-3') | Goal |
|---|---|---|
| AtUMAMIT01 attB1 | GACAAGTTTGTACAAAAAAGCAGGCTCAAGAAGAAAAATGGCAGCTCC | CDS cloning |
| AtUMAMIT01 Kpn attB2 | GACCACTTTGTACAAGAAAGCTGGGTACYATTTACCAGTCTCATCTCTGTAAATGAA | CDS cloning |
| AtUMAMIT02 attB1 | GACAAGTTTGTACAAAAAAGCAGGCTCAATCGGAAAGATGACGGCG | CDS cloning |
| AtUMAMIT02 Kpn attB2 | GACCACTTTGTACAAGAAAGCTGGGTACYAGTCTGCAGATTTCACAGAGGAACT | CDS cloning |
| AtUMAMIT02p attB1 | GACAAGTTTGTACAAAAAAGCAGGCTCGGTTTGGTTTCAGTTTGGA | Promoter cloning |
| AtUMAMIT02p attB2 | GACCACTTTGTACAAGAAAGCTGGGTAGCCGTCATCTTTCCGATTT | Promoter cloning |
| AtUMAMIT03 attB1 | GACAAGTTTGTACAAAAAAGCAGGCTCAATGGAGAGTACGGTGGAAAGAGAG | CDS cloning |
| AtUMAMIT03 Kpn attB2 | GACCACTTTGTACAAGAAAGCTGGGTACYAATCATCGGATTTTGGTGAAGAGAT | CDS cloning |
| AtUMAMIT03p attB1 | GACAAGTTTGTACAAAAAAGCAGGCTCGACGCAGTTACGGCAA | Promoter cloning |
| AtUMAMIT03p attB2 | GACCACTTTGTACAAGAAAGCTGGGTATCTTTCCACCGTACTCTCCAT | Promoter cloning |
| AtUMAMIT04 attB1 | GACAAGTTTGTACAAAAAAGCAGGCTCAAAAGAGAAAATGGGGAAAGGAGTA | CDS cloning |
| AtUMAMIT04 Kpn attB2 | GACCACTTTGTACAAGAAAGCTGGGTACYACACTTCAGATTCAGAATTACTCTTCTTG | CDS cloning |
| AtUMAMIT04p attB1 | GACAAGTTTGTACAAAAAAGCAGGCTTTTGGGGTTTCTTGGTATCT | Promoter cloning |
| AtUMAMIT04p attB2 | GACCACTTTGTACAAGAAAGCTGGGTATGATACTACTCCTTTCCCCAT | Promoter cloning |
| AtUMAMIT05 attB1 | GACAAGTTTGTACAAAAAAGCAGGCTCAATGGCGGATAACACCGATAAT | CDS cloning |
| AtUMAMIT05 Kpn attB2 | GACCACTTTGTACAAGAAAGCTGGGTACYAAACATTGTCCGTTGACTGATGG | CDS cloning |
| AtUMAMIT06 attB1 | GACAAGTTTGTACAAAAAAGCAGGCTCAAGAACGATGTCTCCGATACCG | CDS cloning |
| AtUMAMIT06 Kpn attB2 | GACCACTTTGTACAAGAAAGCTGGGTACYAGGAAGAAATTAATGGTTGACTAATAGG | CDS cloning |
| AtUMAMIT07 attB1 | GACAAGTTTGTACAAAAAAGCAGGCTCAGCCATGCAATTTTTTTGTGTAAAT | CDS cloning |
| AtUMAMIT07 Kpn attB2 | GACCACTTTGTACAAGAAAGCTGGGTACYAAGACAAAAGAGGCTTCTTAGAGTCG | CDS cloning |
| AtUMAMIT08 attB1 | GACAAGTTTGTACAAAAAAGCAGGCTCATTTTGTGTTGAAAAAATGATGAAGA | CDS cloning |
| AtUMAMIT08 Kpn attB2 | GACCACTTTGTACAAGAAAGCTGGGTACYAAAGCAAGAGAGGCTTCTGAAGAT | CDS cloning |
| AtUMAMIT09 attB1 | GACAAGTTTGTACAAAAAAGCAGGCTCAATGTCGATGGAGGAGATTAGTAGC | CDS cloning |
| AtUMAMIT09 Kpn attB2 | GACCACTTTGTACAAGAAAGCTGGGTACYATGGCTCTTGGATAGTCTTTTTCA | CDS cloning |
| AtUMAMIT10 attB1 | GACAAGTTTGTACAAAAAAGCAGGCTCATTTGATATGGGGTTAAAGATGTCA | CDS cloning |
| AtUMAMIT10 Kpn attB2 | GACCACTTTGTACAAGAAAGCTGGGTACYACGAGAAACCCACCGCCACTTTA | CDS cloning |
| AtUMAMIT11 attB1 | GACAAGTTTGTACAAAAAAGCAGGCTCAAAGATGGGATTAAGGATGTCAGAA | CDS cloning |
| AtUMAMIT11 Kpn attB2 | GACCACTTTGTACAAGAAAGCTGGGTACYAGTTAGGTCGTGAAACGCTATGC | CDS cloning |
| AtUMAMIT11p attB1 | GACAAGTTTGTACAAAAAAGCAGGCTCGGATAAGCAATTCTAGGCAGA | Promoter cloning |
| AtUMAMIT11p attB2 | GACCACTTTGTACAAGAAAGCTGGGTATTCTGACATCCTTAATCCCATC | Promoter cloning |
| AtUMAMIT12 attB1 | GACAAGTTTGTACAAAAAAGCAGGCTCTTCAGTGATGGAGGAAGTAAAGA | CDS cloning |
| AtUMAMIT12 Kpn attB2 | GACCACTTTGTACAAGAAAGCTGGGTACYAACCGTTAAGTGAGACTGTTTCTACA | CDS cloning |
| AtUMAMIT12p attB1 | GACAAGTTTGTACAAAAAAGCAGGCTCCTTCCAAGTGCAGAATAATGA | Promoter cloning |
| AtUMAMIT12p attB2 | GACCACTTTGTACAAGAAAGCTGGGTACCTCTTCTTTACTTCCTCCATCA | Promoter cloning |
| AtUMAMIT13 attB1 | GACAAGTTTGTACAAAAAAGCAGGCTCAGATCTTGCAGAGATGGCATGTAT | CDS cloning |
| AtUMAMIT13 Kpn attB2 | GACCACTTTGTACAAGAAAGCTGGGTACYAGACTGATTCTATCACTGTTCTTCTTTG | CDS cloning |

| | | |
|---|---|---|
| AtUMAMIT14 attB1 | GACAAGTTTGTACAAAAAAGCAGGCTCAGATATGGCTTTAAAAACATGGAAG | CDS cloning |
| AtUMAMIT14 Kpn attB2 | GACCACTTTGTACAAGAAAGCTGGGTACYAGACTGATTCATTGGTGTTAGGCCT | CDS cloning |
| AtUMAMIT14 E122R | TGGCATGGATCTTTAGACTTCGTAAGGTTAACGTCAAGAAAAT | Mutagenesis |
| AtUMAMIT14 G144V | GAACAATTGTGACAGTTGGTGTTGCAATGCTTATGACTGTTGT | Mutagenesis |
| AtUMAMIT14 G315V | GAGCAATAGTGATAGTTCTTGTTCTCTATTCGGTTTTGTGGGG | Mutagenesis |
| AtUMAMIT14 G33V | CGAAATTCGCACTGAACCAAGTTATGAGTCCTCACGTTCTAGC | Mutagenesis |
| AtUMAMIT14 K134E | AGAAAATACACAGCCAAGCCGAAATTCTAGGAACAATTGTGAC | Mutagenesis |
| AtUMAMIT15 attB1 | GACAAGTTTGTACAAAAAAGCAGGCTCAAGAATGAAGTTTGAGAGAGCAAGG | CDS cloning |
| AtUMAMIT15 Kpn attB2 | GACCACTTTGTACAAGAAAGCTGGGTACYACTCTTGAGACCGAGCTGCTG | CDS cloning |
| AtUMAMIT17 attB1 | GACAAGTTTGTACAAAAAAGCAGGCTCAAGGAAGAAAATGAAAGGAGGAAAG | CDS cloning |
| AtUMAMIT17 Kpn attB2 | GACCACTTTGTACAAGAAAGCTGGGTACYACACTTGGGTATTGTTAGTCACACC | CDS cloning |
| AtUMAMIT18 attB1 | GACAAGTTTGTACAAAAAAGCAGGCTCAATAAAGATGAAAGGTGGAAGCATG | CDS cloning |
| AtUMAMIT18 Kpn attB2 | GACCACTTTGTACAAGAAAGCTGGGTACYAGGTACTGGTAACCACACCGTTAGT | CDS cloning |
| AtUMAMIT19 attB1 | GACAAGTTTGTACAAAAAAGCAGGCTCAATGGGGAGAGGGTTAATGAATAGT | CDS cloning |
| AtUMAMIT19 Kpn attB2 | GACCACTTTGTACAAGAAAGCTGGGTACYAAATCTCAACTTGAGTAGTAGTAGTAGCC | CDS cloning |
| AtUMAMIT20 attB1 | GACAAGTTTGTACAAAAAAGCAGGCTCATCAATAACCATGGAGGGTGTTAGT | CDS cloning |
| AtUMAMIT20 Kpn attB2 | GACCACTTTGTACAAGAAAGCTGGGTACYATATTCCAGATGTGTTACTATGCGC | CDS cloning |
| AtUMAMIT21 attB1 | GACAAGTTTGTACAAAAAAGCAGGCTCAATGGACATGGAGTCGAAGAAAC | CDS cloning |
| AtUMAMIT21 Kpn attB2 | GACCACTTTGTACAAGAAAGCTGGGTACYAACTAATGACAACCTTCACTTCATTG | CDS cloning |
| AtUMAMIT22 attB1 | GACAAGTTTGTACAAAAAAGCAGGCTCATATTACACGATGATGATGGAGCAC | CDS cloning |
| AtUMAMIT22 Kpn attB2 | GACCACTTTGTACAAGAAAGCTGGGTACYAGACGATGATAACTTCTTCTATAATTTCTTT | CDS cloning |
| AtUMAMIT22p attB1 | GACAAGTTTGTACAAAAAAGCAGGCTCTTTAGGAGCATCTCCAAGTGT | Promoter cloning |
| AtUMAMIT22p attB2 | GACCACTTTGTACAAGAAAGCTGGGTAGGCTTTGTGCTCCATCATC | Promoter cloning |
| AtUMAMIT23 attB1 | GACAAGTTTGTACAAAAAAGCAGGCTCAGAAATGAAAGATATAACGGCAATG | CDS cloning |
| AtUMAMIT23 Kpn attB2 | GACCACTTTGTACAAGAAAGCTGGGTACYAAGGGACATTTGTACTTAATGTTGG | CDS cloning |
| AtUMAMIT23p attB1 | GACAAGTTTGTACAAAAAAGCAGGCTAAATTGTTGGCAGGAACTAGAT | Promoter cloning |
| AtUMAMIT23p attB2 | GACCACTTTGTACAAGAAAGCTGGGTACATTGCCGTTATATCTTTCAT | Promoter cloning |
| AtUMAMIT24 attB1 | GACAAGTTTGTACAAAAAAGCAGGCTCAGGAGAAATGAAGAGTGTAGTTGCA | CDS cloning |
| AtUMAMIT24 Kpn attB2 | GACCACTTTGTACAAGAAAGCTGGGTACYAGGGGACATCTCTATTTACTGATGAA | CDS cloning |
| AtUMAMIT24p attB1 | GACAAGTTTGTACAAAAAAGCAGGCTCGTATAGGGAACAATAGGAACA | Promoter cloning |
| AtUMAMIT24p attB2 | GACCACTTTGTACAAGAAAGCTGGGTACTTCATTTCTCCCCACATT | Promoter cloning |
| AtUMAMIT25 attB1 | GACAAGTTTGTACAAAAAAGCAGGCTCAGAGATGGCTAAATCAGATATGTTGC | CDS cloning |
| AtUMAMIT25 Kpn attB2 | GACCACTTTGTACAAGAAAGCTGGGTACYAAGGCGATGTAGACCTTGTGG | CDS cloning |
| AtUMAMIT26 attB1 | GACAAGTTTGTACAAAAAAGCAGGCTCAATGGGTGAGGATATGCGAGTAGT | CDS cloning |
| AtUMAMIT26 Kpn attB2 | GACCACTTTGTACAAGAAAGCTGGGTACYATGCTTCCATATCTTTGTTCTTACC | CDS cloning |
| AtUMAMIT27 attB1 | GACAAGTTTGTACAAAAAAGCAGGCTCAGAGGGAGAGATGGGTGAGGATAT | CDS cloning |
| AtUMAMIT27 Kpn attB2 | GACCACTTTGTACAAGAAAGCTGGGTACYAGACTCGTTGATCTTCGTTGTTCAT | CDS cloning |
| AtUMAMIT28 attB1 | GACAAGTTTGTACAAAAAAGCAGGCTCAAGAGATATGGCTGGAGATATGCA | CDS cloning |
| AtUMAMIT28 Kpn attB2 | GACCACTTTGTACAAGAAAGCTGGGTACYAAACGGGCGACTTAGAGTCGTTAT | CDS cloning |
| AtUMAMIT28p attB1 | GACAAGTTTGTACAAAAAAGCAGGCTTCTCTTCTTTTGTAAGAGCCTT | Promoter cloning |
| AtUMAMIT28p attB2 | GACCACTTTGTACAAGAAAGCTGGGTATCCTTGCATATCTCCAGCCAT | Promoter cloning |
| AtUMAMIT29 attB1 | GACAAGTTTGTACAAAAAAGCAGGCTCAGAGAGAGAGAGAGGGATGATGAAG | CDS cloning |
| AtUMAMIT29 Kpn attB2 | GACCACTTTGTACAAGAAAGCTGGGTACYAAACGGGCAAATTAGTATCCTTATG | CDS cloning |

| | | |
|---|---|---|
| AtUMAMIT29p attB1 | GACAAGTTTGTACAAAAAAGCAGGCTCCGATTTCTCCAAATTCAGGTA | Promoter cloning |
| AtUMAMIT29p attB2 | GACCACTTTGTACAAGAAAGCTGGGTACCATTGTTCTTCCTTCATCAT | Promoter cloning |
| AtUMAMIT30 attB1 | GACAAGTTTGTACAAAAAAGCAGGCTCACTAATCATGGGTTTTATCGATGGA | CDS cloning |
| AtUMAMIT30 Kpn attB2 | GACCACTTTGTACAAGAAAGCTGGGTACYAAGGAGTCATCGGAATAACCAAAAG | CDS cloning |
| AtUMAMIT30p attB1 | GACAAGTTTGTACAAAAAAGCAGGCTTTTGTTTTGATGCAAGTAGTAGT | Promoter cloning |
| AtUMAMIT30p attB2 | GACCACTTTGTACAAGAAAGCTGGGTACTTTCCATCGATAAAACCCAT | Promoter cloning |
| AtUMAMIT31 attB1 | GACAAGTTTGTACAAAAAAGCAGGCTCAATAATGGGTTACTGCGATGGTAAA | CDS cloning |
| AtUMAMIT31 Kpn attB2 | GACCACTTTGTACAAGAAAGCTGGGTACYAAGGAGTCATTGGAACAACCATAAG | CDS cloning |
| AtUMAMIT32 attB1 | GACAAGTTTGTACAAAAAAGCAGGCTCAATGGTAAAGTTTGATACAAAACTATGGA | CDS cloning |
| AtUMAMIT32 Kpn attB2 | GACCACTTTGTACAAGAAAGCTGGGTACYATTTTGCAGACACTTGATGGG | CDS cloning |
| AtUMAMIT33 attB1 | GACAAGTTTGTACAAAAAAGCAGGCTCAGAGATGGAGATATCGAAATACAAGG | CDS cloning |
| AtUMAMIT33 Kpn attB2 | GACCACTTTGTACAAGAAAGCTGGGTACYACATGAGAAGAGGTTCTAATAGCTCC | CDS cloning |
| AtUMAMIT33p attB1 | GACAAGTTTGTACAAAAAAGCAGGCTTGAGTTCGTTGAAGGTTCGT | Promoter cloning |
| AtUMAMIT33p attB2 | GACCACTTTGTACAAGAAAGCTGGGTATATCTCCATCTCTCTCTCTCTCT | Promoter cloning |
| AtUMAMIT34 attB1 | GACAAGTTTGTACAAAAAAGCAGGCTCAATGGGGAAGATAGAGGAGTACAAG | CDS cloning |
| AtUMAMIT34 Kpn attB2 | GACCACTTTGTACAAGAAAGCTGGGTACYAGTATAGTTGTTGATGTGTTTGGATTTC | CDS cloning |
| AtUMAMIT34p attB1 | GACAAGTTTGTACAAAAAAGCAGGCTGACTCCCTCTCTGCACCAAGA | Promoter cloning |
| AtUMAMIT34p attB2 | GACCACTTTGTACAAGAAAGCTGGGTAGTACTCCTCTATCTTCCCCATGA | Promoter cloning |
| AtUMAMIT35 attB1 | GACAAGTTTGTACAAAAAAGCAGGCTCAATGGGATTAATGGATACAAGATGG | CDS cloning |
| AtUMAMIT35 Kpn attB2 | GACCACTTTGTACAAGAAAGCTGGGTACYAAACTTGATCAAAATCACCATTCG | CDS cloning |
| AtUMAMIT36 attB1 | GACAAGTTTGTACAAAAAAGCAGGCTCAATGGAGGTGAAGGTTAGAAGAGAT | CDS cloning |
| AtUMAMIT36 Kpn attB2 | GACCACTTTGTACAAGAAAGCTGGGTACYATACAGGACTTTCTTCCTGGTTAAT | CDS cloning |
| AtUMAMIT37 attB1 | GACAAGTTTGTACAAAAAAGCAGGCTCAATCAGATCTATGACAAGAGGAGCA | CDS cloning |
| AtUMAMIT37 Kpn attB2 | GACCACTTTGTACAAGAAAGCTGGGTACYAGTCACAGCTTAATGAGGCTTCTTC | CDS cloning |
| AtUMAMIT38 attB1 | GACAAGTTTGTACAAAAAAGCAGGCTCAATGAGAGAAGAGACAGTTTCTTGGA | CDS cloning |
| AtUMAMIT38 Kpn attB2 | GACCACTTTGTACAAGAAAGCTGGGTACYAATCTTCTTCTCTGTCGTGTGAAGG | CDS cloning |
| AtUMAMIT40 attB1 | GACAAGTTTGTACAAAAAAGCAGGCTCAATGAGAGAAGCAGGAGAAGAGAAA | CDS cloning |
| AtUMAMIT40 Kpn attB2 | GACCACTTTGTACAAGAAAGCTGGGTACYAGCTTAATGGAAAGGCTCCATCT | CDS cloning |
| AtUMAMIT41 attB1 | GACAAGTTTGTACAAAAAAGCAGGCTCAGAGAAAGAGAAGAAGATGGCGC | CDS cloning |
| AtUMAMIT41 Kpn attB2 | GACCACTTTGTACAAGAAAGCTGGGTACYATACATGTTCATCGTTCTTGTAACTTTC | CDS cloning |
| AtUMAMIT42 attB1 | GACAAGTTTGTACAAAAAAGCAGGCTCAAAAATGGTTCATGGAAGGTTATGT | CDS cloning |
| AtUMAMIT42 Kpn attB2 | GACCACTTTGTACAAGAAAGCTGGGTACYAGCTTTTAAAATTATCCAAGAGAGGA | CDS cloning |
| AtUMAMIT42p attB1 | GACAAGTTTGTACAAAAAAGCAGGCTGAAGCTCTGCAAATCTGGCTTA | Promoter cloning |
| AtUMAMIT42p attB2 | GACCACTTTGTACAAGAAAGCTGGGTAACATAACCTTCCATGAACCATT | Promoter cloning |
| AtUMAMIT44 attB1 | GACAAGTTTGTACAAAAAAGCAGGCTCAACAATGGCCTCCATTACTCTCC | CDS cloning |
| AtUMAMIT44 Kpn attB2 | GACCACTTTGTACAAGAAAGCTGGGTACYATATTCGGTCGTCTATGTTTTCGT | CDS cloning |
| AtUMAMIT45 attB1 | GACAAGTTTGTACAAAAAAGCAGGCTCATATCATTTTCCAATAATGGCCC | CDS cloning |
| AtUMAMIT45 Kpn attB2 | GACCACTTTGTACAAGAAAGCTGGGTACYAGCCGTTCAATAGAAGAGGGGT | CDS cloning |
| AtUMAMIT46 attB1 | GACAAGTTTGTACAAAAAAGCAGGCTCACAAATTTTTCTACCAATGCCAATA | CDS cloning |
| AtUMAMIT46 Kpn attB2 | GACCACTTTGTACAAGAAAGCTGGGTACYAAACTTGGTCGTTTTTGCGGTTTAA | CDS cloning |
| AtUMAMIT46p attB1 | GACAAGTTTGTACAAAAAAGCAGGCTTAAACCATACCTCAAAAAACTGTGG | Promoter cloning |
| AtUMAMIT46p attB2 | GACCACTTTGTACAAGAAAGCTGGGTATCCGGCCATTATTGGCAT | Promoter cloning |
| AtUMAMIT47 attB1 | GACAAGTTTGTACAAAAAAGCAGGCTCATTTCTACCATTTCCAATGGCC | CDS cloning |

| | | |
|---|---|---|
| AtUMAMIT47 Kpn attB2 | GACCACTTTGTACAAGAAAGCTGGGTACYAAATTTGGTCGTTTTTGCCG | CDS cloning |
| AtUMAMIT14-01 SOE f | CACTCTGCAATGACATTGGTTC | N-terminus exchange |
| AtUMAMIT14-01 SOE r | ACCAATGTCATTGCAGAGTGTGGCTTCCATGTTTTTAAAGC | N-terminus exchange |
| AtUMAMIT14-08 SOE f | ATCGGAGGATTGGCCG | N-terminus exchange |
| AtUMAMIT14-08 SOE r | GCTCCGGCCAATCCTCCGATTGGCTTCCATGTTTTTAAAGC | N-terminus exchange |
| AtUMAMIT14-33 SOE f | GTGCTAGCGTTAGTGATGTTGC | N-terminus exchange |
| AtUMAMIT14-33 SOE r | AACATCACTAACGCTAGCACTGGCTTCCATGTTTTTAAAGC | N-terminus exchange |
| AtUMAMIT14-34 SOE f | GTAATGGCGATGACGATGATT | N-terminus exchange |
| AtUMAMIT14-34 SOE r | ATCATCGTCATCGCCATTACTGGCTTCCATGTTTTTAAAGC | N-terminus exchange |
| AtUMAMIT14-40 SOE f | TTTGCTGCGATGTTTGCG | N-terminus exchange |
| AtUMAMIT14-40 SOE r | ACCGCAAACATCGCAGCAAATGGCTTCCATGTTTTTAAAGC | N-terminus exchange |
| AtUMAMIT14-44 SOE f | TTGACGGCCATGTTGGC | N-terminus exchange |
| AtUMAMIT14-44 SOE r | GTCGCCAACATGGCCGTCAATGGCTTCCATGTTTTTAAAGC | N-terminus exchange |
